# Following the fate of lytic polysaccharide monooxygenases (LPMOs) under oxidative conditions by NMR spectroscopy

**DOI:** 10.1101/2023.02.02.526831

**Authors:** Idd A. Christensen, Vincent G. H. Eijsink, Anton A. Stepnov, Gaston Courtade, Finn L. Aachmann

## Abstract

Lytic polysaccharide monooxygenases (LPMOs) are copper-dependent enzymes that catalyze oxidative cleavage of polysaccharides, such as cellulose and chitin. LPMO action is key to the efficient varlorization of biomass, but the instability of LPMOs in turnover conditions limits their efficiency. LPMO catalysis requires the presence of a reductant, such as ascorbic acid, and hydrogen peroxide, which can be generated *in situ* in the presence of molecular oxygen and various electron donors.. While it is known that reduced LPMOs are prone to auto-catalytic oxidative damage due to off-pathway reactions with the oxygen co-substrate, little is known about the structural consequences of such damage. Here, we present atomic-level insight into how the structure of the chitin-active *Sm*LPMO10A is affected by oxidative damage, using NMR and CD spectroscopy. Incubation with ascorbic acid, led to rearrangements of aromatic residues, followed by more profound structural changes near the copper active site and loss of activity. Longer incubation times induced changes in larger parts of the structure, indicative of progressing oxidative damage. Incubation with ascorbic acid in the presence of chitin led to similar changes in the observable (i.e., not substrate-bound) fraction of the enzyme. Upon subsequent addition of H_2_O_2_, which drastically speeds up chitin hydrolysis, NMR signals corresponding to seemingly intact *Sm*LPMO10A reappeared, indicating dissociation of catalytically competent LPMO. Activity assays confirmed that *Sm*LPMO10A retained catalytic activity when pre-incubated with chitin before being subjected to conditions that induce oxidative damage. Overall, this study provides structural insights into the process of oxidative damage of *Sm*LPMO10A and demonstrates the protective effect of the substrate. The impact of turnover conditions on aromatic residues in the core of the enzyme suggests a role for these residues in dealing with redox-active species generated in the copper center.

## Introduction

The discovery of lytic polysaccharide monooxygenases (LPMOs) more than a decade ago has changed our understanding of enzymatic biomass degradation^1–5^. LPMOs are monocopper enzymes that degrade recalcitrant polysaccharides like cellulose and chitin^4,6–10^ by disrupting the crystalline structure of these polysaccharides^4,6–12^. The reaction mechanism used by LPMOs is not fully uncovered ^13,14^, but involves either O_2_ or H_2_O_2_ as a co-substrate, and an external reductant^4,14–18^. The copper active site is part of a solvent-exposed substrate-binding surface. It comprises two conserved histidines (one of which is the N-terminal residue) that coordinate a single copper ion ^6,8–10^. LPMOs share similar three-dimensional structures, consisting of a β-sandwich core with β-strands linked through loops containing several short helices, believed to influence substrate specificity and oxidative regioselectivity ^19–21^.

Chitin- and cellulose-active LPMOs cleave the β(1,4) glycosidic bonds in their substrates in a regioselective manner, acting on the C1- and/or C4- carbon ^21,22^. The LPMO reaction (Figure 1) starts with the reduction of the copper atom in the active site from Cu(II) to Cu(I) by an external reductant. The solvent-exposed active site ^6,9^ enables copper reduction by both small-molecule reductants such as ascorbic acid and gallic acid ^4,6,23^ or by redox enzyme partners such as cellobiose dehydrogenase ^10,24^ and pyrroloquinoline-quinone dependent dehydrogenase ^25^. The activated LPMO then reacts with molecular O_2_ or H_2_O_2_ to create a reactive oxygen species capable of abstracting a hydrogen atom from the C1 or the C4 of the scissile glycosidic bond. Hydrogen abstraction is followed by hydroxylation through a rebound mechanism, which destabilizes the glycosidic bond and leads to its cleavage ^10,13–15,17,26^. The peroxygenase reaction with H_2_O_2_ is much faster than the monooxygenase reaction with O_2_ ^18,27–29^ and there is some debate in the field regarding the kinetic relevance of the latter^17,18^. Although copper reduction has been shown to increase substrate affinity ^9,30^ it is still unclear if LPMOs bind the oxygen co-substrate before or after they bind to the substrate. LPMO activity significantly boosts the performance of hydrolytic enzymes involved in the depolymerization of crystalline substrates ^2,4,5,31^. However, to efficiently harness the power of LPMOs in commercial enzyme cocktails for polysaccharide saccharification, challenges related to LPMO inactivation must be resolved ^32^.

**Figure 1:**
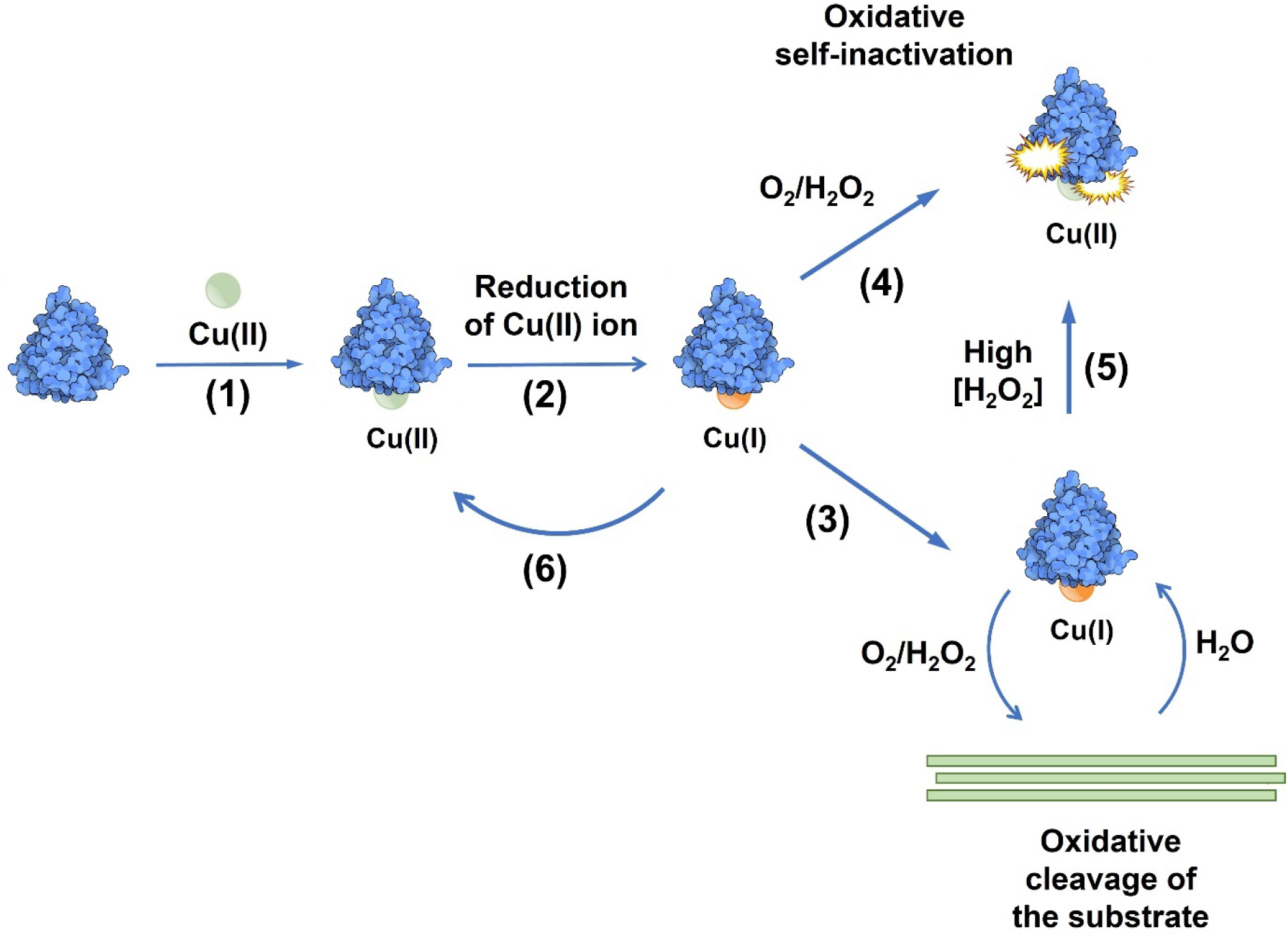
Scheme of the LPMO reaction and off-pathway oxidative self-inactivation. **(1)** A copper atom must be bound to the LPMO active site for the enzyme to be catalytically active. LPMO activity is also dependent on (**2**) a priming reaction where Cu(II) is reduced to Cu(I) by a reductant such as ascorbic acid. **(3)** In the presence of substrate, the reduced LPMO (provided with an oxygen containing co-substrate) catalyzes oxidative cleavage of glycosidic bonds in the substrate. **(4)** Alternatively, an off-pathway reaction occurs which may lead to oxidative damage by uncontrolled reactive oxygen species. **(5)** The ratio between on-pathway and off-pathway reactions depends on the concentrations of both substrate and H_2_O_2_. **(6)** The off-pathway reactions may lead to copper oxidation without enzyme damage, meaning that re-reduction of the LPMO is needed to become active. Of note, damage to the enzyme may lead to the release of free copper into solution.

In solution, reduced LPMOs are susceptible to inactivation caused by reactions between the activated LPMO and its co-substrate, in a process referred to as oxidative self-inactivation^15,33,34^. Oxidative self-inactivation occurs when H_2_O_2_ reacts with the non-substrate bound reduced LPMO, leading to off-pathway reactions that may cause oxidative damage to catalytically important residues ^15,33,35^. Even in LPMO reactions without added H_2_O_2_, this oxidant will be available due to several side-reactions. LPMOs will produce H_2_O_2_ when incubated with an appropriate reductant (like ascorbic acid) and O_2_ in the absence of their substrate ^15,36,37^. In addition, H_2_O_2_ may be generated through reactions between the reductant and O_2_ ^23,38^. The oxidation of certain reductants (including the much-used ascorbic acid) is strongly affected by the presence of transition metals in solution. A recent study showed that even low micromolar concentrations of free copper (i.e., concentrations similar to typically used LPMO concentrations) promote oxidation of ascorbic acid and concomitant production of H_2_O_2_, which may lead to inactivation of LPMOs^16^.. As LPMOs are efficient peroxygenases^15,27^, LPMO reactions can be fueled by extraneously supplied H_2_O_2_. When doing so, the supply of H_2_O_2_ must be carefully controlled to prevent LPMO inactivation^15,28,32^.

The protective effect of the substrate against oxidative self-inactivation has been shown by carrying out LPMO reactions with varying substrate concentrations ^15,37,39^. The presence of Carbohydrate Binding Modules (CBMs), which are believed to keep the LPMO in close proximity to its substate, also reduces LPMO inactivation^39^. Substrate binding will position the reactive oxygen species produced during catalysis in a way that promotes bond cleavage rather than damaging off-pathway reactions^26,30,40^. Interestingly, Bissaro *et al*. identified a tunnel connecting the active site of the chitin-bound *Sm*LPMO10A to the bulk solvent. This would make it possible for the reduced LPMO to activate O_2_ or H_2_O_2_ while substrate-bound ^40^, creating a controlled reaction environment.

Currently, limited information is available regarding the structural effects of oxidative off-pathway reactions in LPMOs. Mass spectrometry-based studies of oxidative damage in cellulose-active *Sc*LPMO10C have shown that oxidative damage primarily occurs close to the copper active site, with the two catalytic histidines and nearby aromatic residues being most exposed to oxidation ^15^. Other parts of the structure seemed largely unaffected ^15^. A more detailed molecular understanding of the oxidative self-inactivation of LPMOs is needed as it may eventually help optimize these enzymes for utilization in industrial biomass saccharification. In addition, one may speculate about the existence of possible protective hole hopping pathways ^41^ in LPMOs, which would be expected to largely involve aromatic residues. The propagation of oxidative damage through the LPMO molecule likely relates to such pathways.

Here we have studied structural changes in an LPMO that is particularly rich in aromatic residues near the catalytic center under oxidative conditions. NMR ^15^N-HSQC (Heteronuclear Single Quantum Coherence) spectra and CD spectra were used to monitor structural changes occurring in chitin-active *Sm*LPMO10A (also known as CBP21) incubated with ascorbic acid under aerobic conditions and/or H_2_O_2_ in the absence of substrate. Similar treatments in the presence of chitin were used to investigate the protective effect of the substrate against oxidative damage and self-inactivation.

## Materials and Methods

The methods and experimental design used to monitor structural changes in *Sm*LPMO10Aunder oxidative conditions in either the presence or absence of chitin are summarized in Figure 2 and Figure S1.

**Figure 2:**
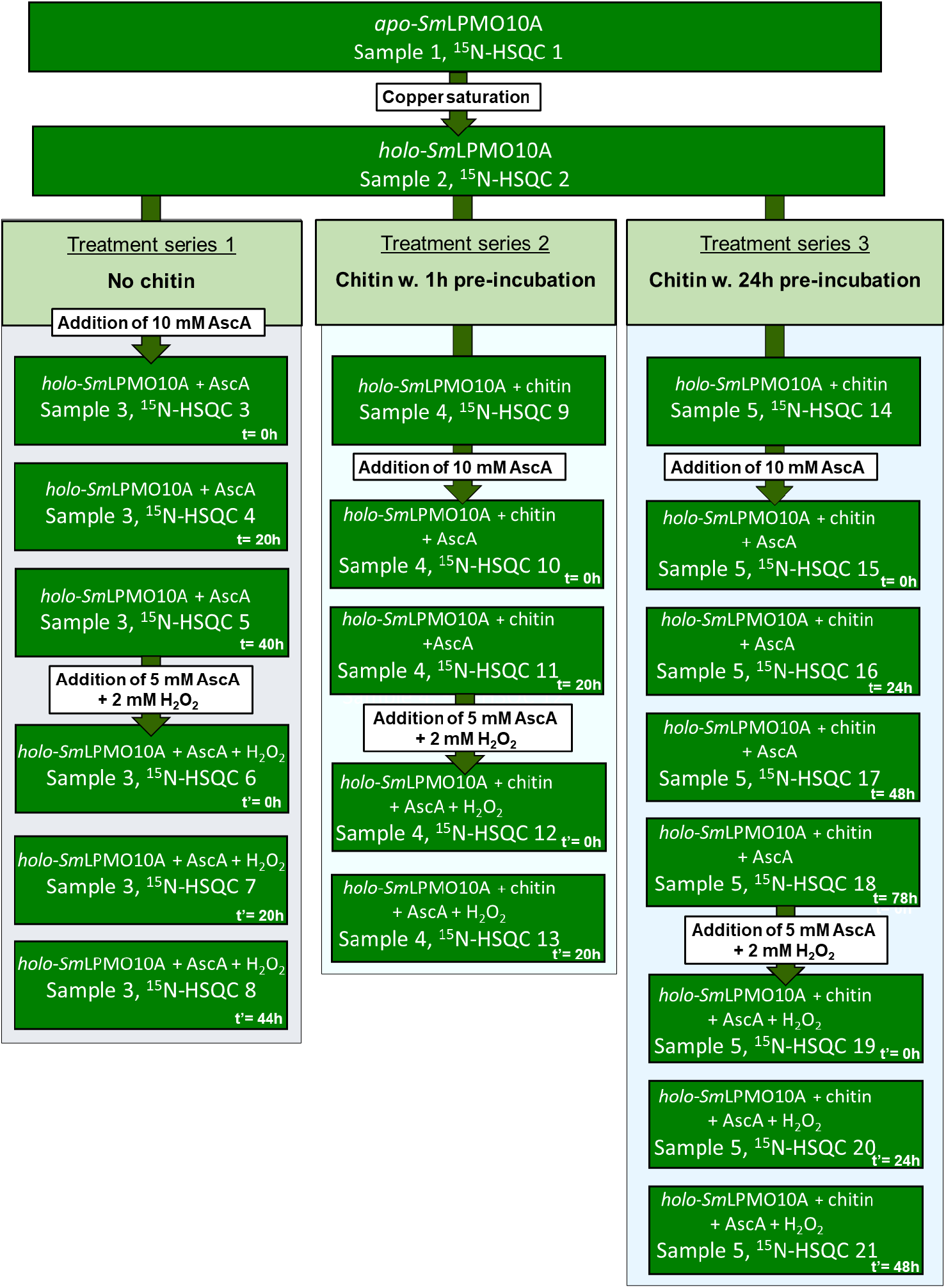
Flow diagram of the incubations and NMR experiments done to investigate copper active site reduction and oxidative self-inactivation of *Sm*LPMO10A. The rectangular dark green text boxes represent the time points when NMR spectra of *Sm*LPMO10A were recorded. The experiments were done using three treatment series. In all treatment series copper was added to a sample of *apo*-*Sm*LPMO10A in a 3:1 molar ratio to obtain *holo*-*Sm*LPMO10A. Note that excess (i.e. unbound) Cu(II) ions were removed from the samples by desalting and the samples were then concentrated to samples of ~ 150 μM protein. Ascorbic acid (AscA) was added as indicated, without (series 1) or with (series 2 & 3) prior addition of chitin. After approximately two days, ascorbic acid and hydrogen peroxide were added, as indicated, and time points from this second phase of the incubation experiment are referred to as t’ rather than t. Time points t = 0 and t’ = 0 refer to approximately 5 minutes after addition. The recording time for each ^15^N-HSQC spectrum was 40 minutes.

### Sample preparation of *apo-Sm*LPMO10A for NMR

Cloning, production, purification, and NMR sample preparation of both natural isotope abundance, ^15^N- and ^13^C,^15^N- isotopically enriched *Sm*LPMO10A samples (UniProt entry O83009, residues 28-197) was carried out as previously described ^42^. The protein concentration was determined by measuring the A_280nm_ of the samples using a Nanodrop ND-1000 spectrophotometer (NanoDrop). The A_280nm_ value was then used to calculate the protein concentration based on the theoretical extinction coefficient (ϵ = 35,200 M^−1^·cm^−1^) obtained using the ProtParam tool (https://web.expasy.org/protparam/).

To obtain the *apo* form of *Sm*LPMO10A, the purified enzyme samples were incubated at 4°C overnight in acetate buffer (25 mM sodium acetate, 10 mM NaCl, pH 5.0) with 2 mM EDTA. A buffer exchange with acetate buffer (25 mM sodium-acetate, 10 mM NaCl, pH 5.0) was performed to remove EDTA. A sample were then concentrated to ~ 150 μM with a volume of ~ 500 μL using first, a VivaSpin centrifugation filter (10 kDa cut-off, Sartorius), followed by an Amicon ®Ultra centrifugation filter (3 kDa cut-off, Merck). The sample was transferred to a 5 mm NMR-tube (Labscape™ Essence) and 10% D_2_O (99.9% D, Sigma-Aldrich) was added.

To obtain samples of *holo-Sm*LPMO10A, CuCl_2_ was added to *apo*-*Sm*LPM10A in a 3:1 molar ratio in acetate buffer (25 mM sodium-acetate, 10 mM NaCl, pH 5.0). The sample was then incubated for 30 minutes at room temperature and excess copper was removed by gel filtration using a PD MidiTrap G-25 desalting column (GE Healthcare; Uppsala, Sweden) equilibrated with acetate buffer (25 mM sodium-acetate, 10 mM NaCl, pH 5.0) ^43^. The resulting sample of *holo*-*Sm*LPMO10A was concentrated to ~ 150 μM using an Amicon ®Ultra centrifugation filter (3 kDa cut-off, Merck) and the sample was split into three equal volumes of ~ 500 μL. One of the resulting samples was transferred to a 5 mm NMR tube (LabScape™ Essence) and 10% D_2_O (99.9%, D, Sigma-Aldrich) was added.

Insoluble β-chitin fibers (Mahtani Chitosan) were mechanically treated by milling and sieving to a particle size of approximately 0.5 mm. 10 mg of these β-chitin particles were transferred into two separate 5 mm NMR tubes (LabScape™ Essence). The two remaining *holo*-*Sm*LPMO10A samples were then added to each of the NMR tubes and 10% D_2_O (99.9%, D, Sigma-Aldrich) was added to the samples.

### NMR spectroscopy

All NMR spectra were recorded at 25 °C on a Bruker Avance III HD 800 MHz spectrometer equipped with a 5-mm Z-gradient CP-TCI (H/C/N) cryogenic probe at the NV-NMR-Centre/Norwegian NMR Platform (NNP) at the Norwegian University of Science and Technology (NTNU). ^1^H chemical shifts were internally referenced to the water signal at 25°C, while ^15^N and ^13^C chemical shifts were indirectly referenced to the water signal, based on absolute frequency ratios (Zhang et al. 2003). The NMR data were processed using TopSpin version 4.0.7. The processed NMR spectra were analyzed using CARA version 1.5.5. The previously published NMR assignment of *Sm*LPMO10A (Aachmann et al., 2012) with BMRB entry number 17160 was used.

### *Sm*LPMO10A under oxidative conditions by NMR spectroscopy

The effects of oxidative conditions on the structure of *Sm*LPMO10A were investigated by NMR using the experimental designed outlined Figure 2. *Holo*-SmLPMO10A was subjected to conditions designed to promote oxidative self-inactivation in three treatment series. In treatment series 1, *holo*-SmLPMO10A was incubated in oxidative conditions without substrate, while in treatment series 2 and 3 the enzyme was pre-incubated with chitin for 1 and 24 hours, respectively, before being subjected to oxidative conditions (with the substrate still present).

### Copper binding

Cu(II) binding to the active site of *Sm*LPMO10A was evaluated by recording a ^15^N-HSQC spectrum after saturating with Cu(II) (Figure 2: ^15^N-HSQC 2) and comparing it to the ^15^N-HSQC spectrum of *apo*-*Sm*LPMO10A (Figure 2: ^15^N-HSQC 1) and previously published NMR data^9^.

### Treatment series 1: The effect of oxidative conditions on SmLPMO10A in the absence of substrate

Ascorbic acid (10 mM) was added to a ~ 150 μM sample of *holo*-*Sm*LPMO10A 500 μL in a 5 mm NMR tube (LabScape™ Essence) without chitin (Figure 2: ^15^N-HSQC 3). The effect of ascorbic acid was evaluated by recording ^1^H, ^15^N-HSQC, HNCA and HN(CO)CACB spectra of the sample over the course of 2 days (Figure 2: ^15^N-HSQC 3–5). At this point a ^1^H spectrum was recorded to verify that no more ascorbic acid could be observed. Additional ascorbic acid (5 mM) was added together with H_2_O_2_ (2 mM) to induce extensive oxidative damage of the LPMO. The effect of the combined addition of ascorbic acid and H_2_O_2_ was evaluated by recording ^1^H, ^15^N-HSQC, HNCA, and HN(CO)CACB spectra over the course of another 2 days (Figure 2: ^15^N-HSQC 6–8) after which, a ^1^H spectrum was recorded to verify that no more ascorbic acid could be detected in the sample.

### Treatment 2 and 3: The effect of oxidative conditions on *Sm*LPMO10A in the presence of chitin

To investigate the protective effect of the substrate against oxidative damage, two 500 μL samples of ~ 150 μM *holo*-*Sm*LPMO10A were pre-incubated with 10 mg β-chitin particles (~0.5 mm; Mahtani Chitosan) in an NMR tube for 1 h (treatment series 2) and 24h (treatment series 3), respectively. After the pre-incubation, ^15^N-HSQC spectra of both samples were recorded (Figure 2: ^15^N-HSQCs 9 and 14), before adding 10 mM ascorbic acid to both samples.

In treatment series 2 (1h pre-incubation with chitin),^1^H, ^15^N-HSQC, HNCA, and HNCACB spectra were recorded over the course of 1 day (Figure 2: ^15^N-HSQC 10–11). At this point, a ^1^H spectrum was recorded to verify that no more ascorbic acid could be observed. Additional ascorbic acid (5 mM) was added together with H_2_O_2_ (2 mM) to potentially induce extensive oxidative damage, and ^1^H,^15^N-HSQC, HNCA, and HNCACB spectra were recorded over the course of one day (Figure 2: ^15^N-HSQC 12–13). Finally, a ^1^H spectrum was recorded to verify that no more ascorbic acid could be observed.

In treatment series 3 (24h pre-incubation with chitin), ^1^H and ^15^N-HSQC spectra were recorded every 24 hours for a total of three days (Figure 2: ^15^N-HSCQ 15–18), and ^1^H spectrum was recorded to verify that no more ascorbic acid could be observed. Additional ascorbic acid (5 mM) was added together with H_2_O_2_ (2 mM) to potentially induce extensive oxidative damage. Following this step ^15^N-HSQC spectra were recorded every 24 hours over a period of two days (Figure 2: ^15^N-HSQC 19–21). Finally, a ^1^H spectrum was recorded to verify that no more ascorbic acid could be observed.

### Chemical shift perturbations in response to oxidative conditions

Chemical shift perturbations (CSPs) were analyzed for ^15^N-HSQC spectra of *Sm*LPMO10A by monitoring changes in the ^1^H-^15^N signals. CSP were calculated using the combined chemical shift change (Δδ_comb_) from the equation: 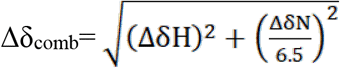, ^44,45^ Δ*δH* = Δ*δH_obs_* − Δ*δH_free_* and Δ*δN* = Δ*δN_obs_* − Δ*δN_free_* are the absolute changes in the chemical shifts in parts per million (ppm) of the amide proton and amide nitrogen, respectively^45^. CSPs were calculated for pairs of consecutively recorded ^15^N-HSQC spectra in each of the treatment series and between each recorded ^15^N-HSQC spectrum and the ^15^N-HSQC reference spectrum of *apo*-*Sm*LPMO10A (^15^N-HSQC 1).

### Linewidth from signal intensity calculations

The linewidths of ^1^H-^15^N signals in all the recorded ^15^N-HSQC spectra of both *apo*- and *holo*-*Sm*LPMO10A were measured by integration of the signal using the CARA software ^46^. The relative change in linewidths were then estimated by calculating the intensity ratios between ^15^N-HSQC spectra using the equation: 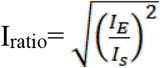, where *I_E_* is the ^1^H-^15^N intensity in each ^15^N-HSQC spectrum corresponding to the sample treatments shown in Figure 2 and *I_S_* is the ^1^H-^15^N intensity in either the previously recorded ^15^N-HSQC spectrum in a treatment series (as shown in Figure 2), or the ^1^H-^15^N intensity in the reference spectrum of *apo*-*Sm*LPMO10A (^15^N-HSQC 1).

### Changes in mobility in response to oxidative conditions

Heteronuclear {^1^H}-^15^N NOEs, *T_1_*- and *T_2_*-relaxation time experiments were recorded for *apo*-*Sm*LPMO10A and for *holo*-*Sm*LPMO10A after a 96-hour incubation time with 10 mM ascorbic acid. A 96-hour incubation period was chosen to ensure that structural changes in *Sm*LPMO10A happening in response to ascorbic acid were completed before recording the spectra. After 96-hours a ^1^H NMR spectrum was recorded to verify that the initially added amount of ascorbic acid was exhausted. Additional ascorbic acid (5 mM) was added together with H_2_O_2_ (2 mM) to induce extensive oxidative damage, and new NOEs, *T_1_*- and *T_2_*-relaxation time experiments were acquired after 72-hours of incubation. The heteronuclear {^1^H}-^15^N heteronuclear NOEs as well as *T_1_* and *T_2_* relaxation data were derived from the spectra using Dynamics Center software version 2.3.1 (Bruker BioSpin).

### Spectroscopy

CD spectroscopy was used to monitor structural changes for treatment series 1 (Figure 2), using samples prepared correspondingly to NMR experiments. Aliquots were collected from samples of 100 μM *apo*-*Sm*LPMO10A alone, 100 μM *apo*-*Sm*LPMO10A after addition of 10 mM ascorbic acid (at t= 4min, 24h, and 48h), and *holo*-*Sm*LPM10A after addition of 10 mM ascorbic acid (at t= 4 min, 30 min, 4h, 20h, 24h, 45h, and 48h). All samples were in acetate buffer (25 mM sodium-acetate, 10 mM NaCl, pH 5.0). The collected aliquots were diluted to a protein concentration of 2.5 μM using acetate buffer and far-UV Circular Dichroism (CD) spectra and UV absorption spectra were recorded using a Chirascan™ qCD spectropolarimeter (Applied Photophysics Ltd. Leatherhead, Great Britain). A 0.1 cm path length quartz cuvette (Sigma-Aldrich) was employed. Reported CD spectra are based on measuring ellipticity the 185-260 nm, and absorbance in range of 185-340 nm, (resolution of 2 nm, a step-size of 0.5 nm and 1.3s per timepoint) and represent the average of 8 scans. All data was recorded at 23 °C. The content of secondary structural elements was analyzed using the BeStSel (Beta Structure selection) server (http://bestsel.elte.hu/index.php) ^47,48^.

### Functional studies of *Sm*LPMO10A under oxidative conditions

Activity tests were performed to assess the catalytic capacity of *holo*-*Sm*LPMO10A after being subjected to treatments like those in treatment series 1 and 3 shown in Figure 2. Two samples of ~ 100 μM *holo*-*Sm*LPMO10A in acetate buffer (25 mM sodium acetate, 10 mM NaCl, pH 5.0), with a volume of 500 μL, were prepared as described above. The samples were each transferred to 5 mm NMR tubes (LabScape™ Essence). To study the protective effect of chitin, 10 mg mechanically milled β-chitin particles with a size of ~ 0.5 mm (Mahtani Chitosan) were added to one of the two NMR tubes while no substrate was added to the other NMR tube. The samples were then pre-incubated for 24 hours at room temperature, after which ascorbic acid (10 mM) was added to both samples. 10 μL aliquots were collected from the NMR tubes at t= 0h (i.e., 30 seconds after addition of ascorbic acid) and at t= 24h, and diluted with 990 μL acetate buffer (25 mM sodium acetate, 10 mM NaCl, pH 5.0) containing 10 mg chitin from shrimp shells (Sigma-Aldrich, practical grade powder). Ascorbic acid was then added to the reactions (1 mM final concentration) followed by incubation for 24 hours at 40 °C in an Eppendorf thermomixer set to 800 RPM. The reactions were stopped by separating the enzyme and soluble products from the insoluble chitin substrate by filtration using 96-well filter plates (Merck). The concentration of oxidized LPMO products was also determined in aliquots taken from the pre-incubation reaction with β-chitin to allow accounting for carry-over of products potentially formed in the pre-incubation phase.

To measure the maximum (i.e., 100%) enzyme activity of *holo*-*Sm*LPMO10A, a reaction with 1 μM LPMO and 10 mg shrimp chitin was carried out in 500 μL 25 mM sodium acetate buffer, pH 5.0, at 40 °C and 800 RPM using an Eppendorf thermomixer. The enzyme was pre-incubated with the substrate for 24 hours at room temperature. The reaction was initiated by adding ascorbic acid to 1 mM final concentration and quenched after 24 hours by filtration as described above. A control reaction lacking enzyme was prepared by incubating 10 mg chitin particles from shrimp shells in 500 μL of 25 mM sodium acetate buffer, pH 5.0 at 40°C and 800 RPM for 24 hours using an Eppendorf thermomixer.

### Quantification of soluble oxidized LPMO products by HPAEC-PAD

Soluble oxidized products generated upon incubating chitin with the LPMO were analysed by high-performance anion-exchange chromatography with pulsed amperometric detection (HPAEC-PAD) using a Dionex ICS5000 system (Thermo Scientific, San Jose, CA, USA) equipped with a CarboPac PA200 analytical column. Prior to analysis, the reaction samples were diluted 2 times with 50 mM sodium-phosphate buffer, pH 6.0 and treated with 1 μM *Sm*GH20 chitobiase (30 °C, overnight)^43^ to convert native and oxidized chito-oligosaccharide products to a mixture of *N*-acetylglucosamine (GlcNAc) and chitobionic acid (GlcNAcGlcNAC1A). A stepwise gradient with an increasing amount of eluent B (eluent B: 0.1 M NaOH and 1 M NaOAc; eluent A: 0.1 M NaOH) was applied to the column at 0.5 ml/min flow rate according to the following program: 0–10% B over 5 min, 10–100% B over 4.5 min, 100-0% B over 6 s, 0% B over 13.5 min. Chromeleon 7.0 software was used for data analysis. Standard solutions of chitobionic acid were prepared in-house as previously described^43^.

## Results

### Exposure to ascorbic acid in the absence of chitin affects residues near the copper active site

The addition of Cu(II) to *apo*-*Sm*LPMO10A (^15^N-HSQC 1 & 2) was accompanied by rapid relaxation of ^1^H-^15^N signals belonging to residues located within ~10 Å from the copper active site (Figure S2). The increased relaxation is due to PRE^49^ (Paramagnetic Relaxation Enhancement) caused by Cu(II) binding to the histidine brace formed by H28 and H114, and is in agreement with previous results obtained by Aachmann *et al*.^9^.

To follow the effect of ascorbic acid on the structure of *holo*-*Sm*LPMO10A (treatment series 1), a total of three ^15^N-HSQC spectra were recorded at time points t= 0h (i.e., just after addition), 20h, and 40h. The immediate (t= 0h) effect of ascorbic acid (Figure S3, ^15^N-HSQC 3) was the loss of PRE on ^1^H-^15^N signals belonging to residues ~10 Å from the active site indicating that the copper was reduced. As PRE influenced the NMR spectra of Cu(II) *holo*-*Sm*LPMO10A, all the ^15^N-HSQC spectra of *holo*-SmLPMO10A recorded after the addition of ascorbic acid were compared with the reference spectrum of *apo*-*Sm*LPMO10A (^15^N-HSQC 1) which is not affected by PRE.

Compared with *apo*-*Sm*LPMO10A, copper binding and addition of ascorbic acid (i.e., binding of reduced copper) produced changes in both the chemical shifts and linewidths for several of the ^1^H-^15^N signals in the HSQC spectrum (Figure S3, ^15^N-HSQC 3). Chemical Shift Perturbations (CSPs) indicate changes in the chemical environment of the nuclei giving rise to the signal ^50^, whereas changes in linewidth (seen by signal intensity) reflect changes in residue dynamics ^51^. Relative to the *apo* enzyme, the reduced, copper containing *holo* enzyme showed large CSPs (>50 Hz) for residues near the active site (Figure S4) immediately (t= 0h) after the addition of ascorbic acid (Figure 3A,C). Among these were residues known to be involved in substrate binding and residues in or near the copper site, such as T111, H114, D182, and F187 ^3,9^. The binding of reduced copper, and surplus concentrations of ascorbic acid also affected larger parts of the protein, including residues, such as A63, A72, G150, and A71 whose signals were not affected by PRE (Fig. 3A), in accordance with previous studies^9^.

**Figure 3:**
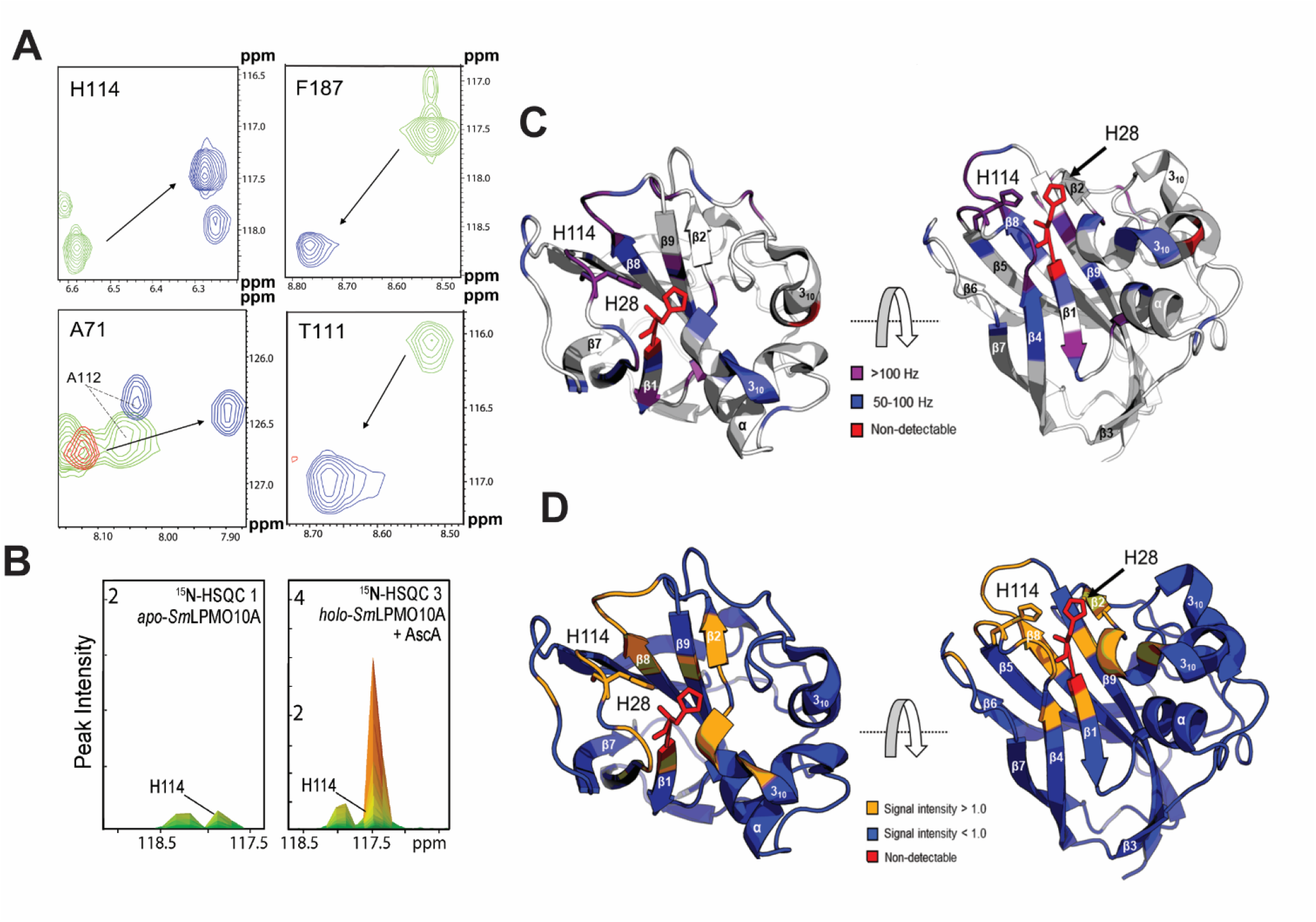
Immediate (t= 0h) effect of incubating *holo*-*Sm*LPMO10A with 10 mM ascorbic acid under aerobic conditions (treatment series 1, ^15^N-HSQC 3). **A)** Selected parts of the ^15^N-HSQC spectra displaying chemical shifts for *apo*-*Sm*LPMO10A (green), *holo*-*Sm*LPMO10A (red), and *holo*-*Sm*LPMO10A immediately (t= 0h) after addition of ascorbic acid (blue). The arrows indicate the direction of the change in chemical shifts upon copper binding and reduction. Note that there are no (red) signals for residues within ~ 10 Å due to the Paramagnetic Relaxation Enhancement caused by Cu(II). **B)** Change in the linewidth, seen as an increase in the signal intensity (found by peak integration), for H114 upon addition of ascorbic acid (at t= 0h) compared to *apo*-*Sm*LPM10A. **C)** Chemical Shift Perturbations (CSPs) in response to addition of ascorbic acid (a t= 0h) and copper reduction compared to *apo*-*Sm*LPMO10A highlighted on the X-ray crystal structure of *Sm*LPMO10A (PDB ID: 2BEM). The magnitude of the change in Hz is indicated by the color scheme. Non-detectable residues are shown in red. No CSPs > 50 Hz were observed for the grey colored areas of the structure. **D)** Changes in linewidths in response to addition of ascorbic acid (at t= 0h) and copper reduction compared with *apo*-*Sm*LPMO10A highlighted on the X-ray crystal structure of *Sm*LPMO10A. An increase in signal intensity is shown in orange, while a decrease is shown in blue. Non-detectable residues are shown in red.

Addition of ascorbic acid also led to narrower linewidths (relative to the *apo*-enzyme), seen by increased signal intensities, for ^1^H-^15^N signals of residues near the copper active site (Figure S4). These increased intensities were accompanied by a relative decrease in signal intensity for the remaining parts of the structure, as shown in Figure 3D. Linewidth analysis can provide information about protein dynamics ^44,51^. The increased ^1^H-^15^N signal intensities near the copper active site are intriguing as they could indicate slow exchange between different conformations of *holo*-SmLPMO10A ^44^, or, interestingly, increased local mobility, following the addition of ascorbic acid.

Taken together, the CSP and the changes in signal intensity show that exposure to ascorbic acid has a major effect on the copper-binding region of the LPMO. These changes exceed the structural effect of copper binding, which is not expected to increase structural flexibility based on previous results ^9^. The most likely explanation for these changes is oxidative damage of the catalytic center. Indeed, LPMO activity measurements showed reduced activity at t = 0 (measured at approximately 5 minute after adding ascorbic acid), whereas LPMO activity was completely abolished at later timepoints, demonstrating that the LPMO is rapidly damaged when exposed to a high concentration of reductant in the absence of substrate.

Longer incubation times (t> 0h) with ascorbic acid did not result in additional CSPs compared to those observed at t= 0h. The number of amino acid residues with narrow linewidths relative to *apo*-*Sm*LPMO10A first grew (Figure S4) at t= 20h after adding ascorbic acid (Figure S5, ^15^N-HSQC 4), potentially meaning that a larger part of the structure displayed local increases in mobility. Further evidence of structural changes was observed at t= 40h (Figure S6, ^15^N-HSQC 5) by line broadening (Figures 4B,C), which is indicative of structural changes in the intermediate exchange regime^51^ and may reflect a decline in the protein’s structural stability ^52^. As a result, the number of residues with narrow linewidths/high signal intensities decreased and was lower than the intensities found at t= 0h sample (Figure 4A). It is conceivable that extended exposure to ascorbic acid in the presence of free copper ions released from damaged LPMOs leads to abiotic formation of reactive-oxygen species that can damage proteins^53^.

**Figure 4:**
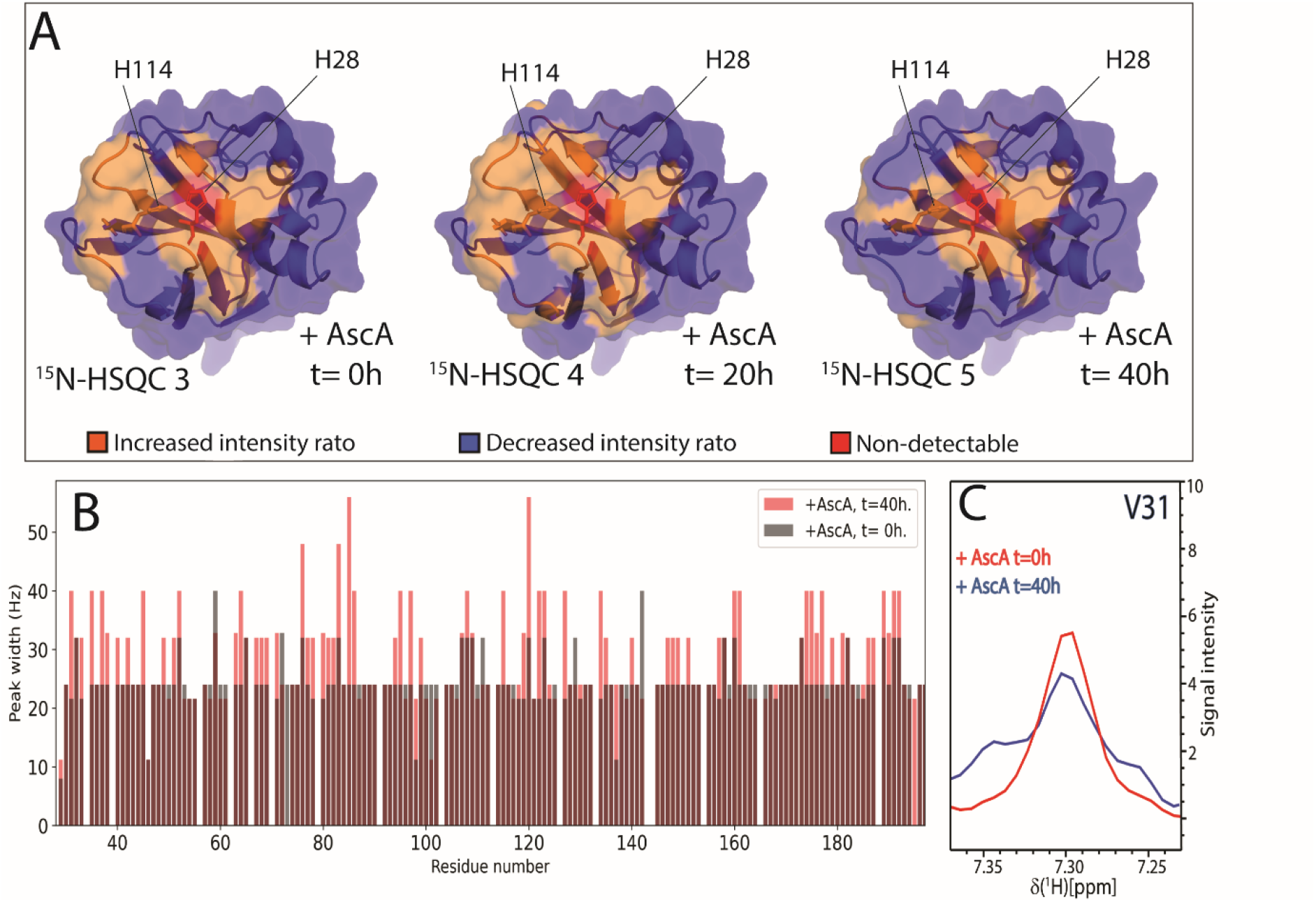
Observed changes in ^1^H-^15^N signal intensities in response to incubation of *holo*-*Sm*LPMO10A with 10 mM ascorbic acid from t= 0h to t= 40h under aerobic conditions (treatment series 1). **A)** Changes in line shape, seen as changes in signal intensity, relative to *apo*-*Sm*LPMO10A highlighted on the structure of *Sm*LPMO10A (PDB ID: 2BEM). The orange color indicates an increase in signal intensity, while the blue color indicates a decrease. Non-detectable residues are shown in red. **B)** ^1^H resonance linewidths for all residues of *Sm*LPMO10A at t= 0h and t= 40h after addition of ascorbic acid. **C)** Line broadening of the ^1^H resonance illustrated with the residue V31 in response to incubation with ascorbic acid.

Subsequent incubation with fresh ascorbic acid and H_2_O_2_ resulted in the signals of several residues disappearing from the ^15^N-HSQC spectra (Figure S7–S9, ^15^N-HSQC 6–8). This indicates that the protein has been damaged and is unfolding/denatured when exposed to a combination of free copper (leaking from damaged LPMOs), ascorbic acid and H_2_O_2_. Immediately after adding H_2_O_2_ (Figure S7, ^15^N-HSQC 6), signals of residues primarily located near the copper active site became undetectable while new signals appeared in the region between 8-9 ppm in the spectrum (Figure 5), indicating partial loss of *Sm*LPMO10As structural integrity ^54^. More pronounced changes, no longer limited to the copper-binding region, were observed in the ^15^N-HSQC spectrum recorded at t’= 44h after the addition of H_2_O_2_ (Figure S9, ^15^N-HSQC 8). Firstly, around 25% of the signals for the protein could no longer be observed in the spectrum. Secondly, the linewidth of many of the remaining detectable signals became broader (Figure S10) and there was an overall decrease in signal dispersion with signals accumulating in the range between 8-9 ppm in the spectrum (Figure 5 and S11). Thirdly, narrow linewidths and CSPs> 50 Hz were observed for still detectable residues near the active site (Figure S4). These spectral changes, taken together, strongly suggest a loss of 3D structure and protein unfolding.

**Figure 5:**
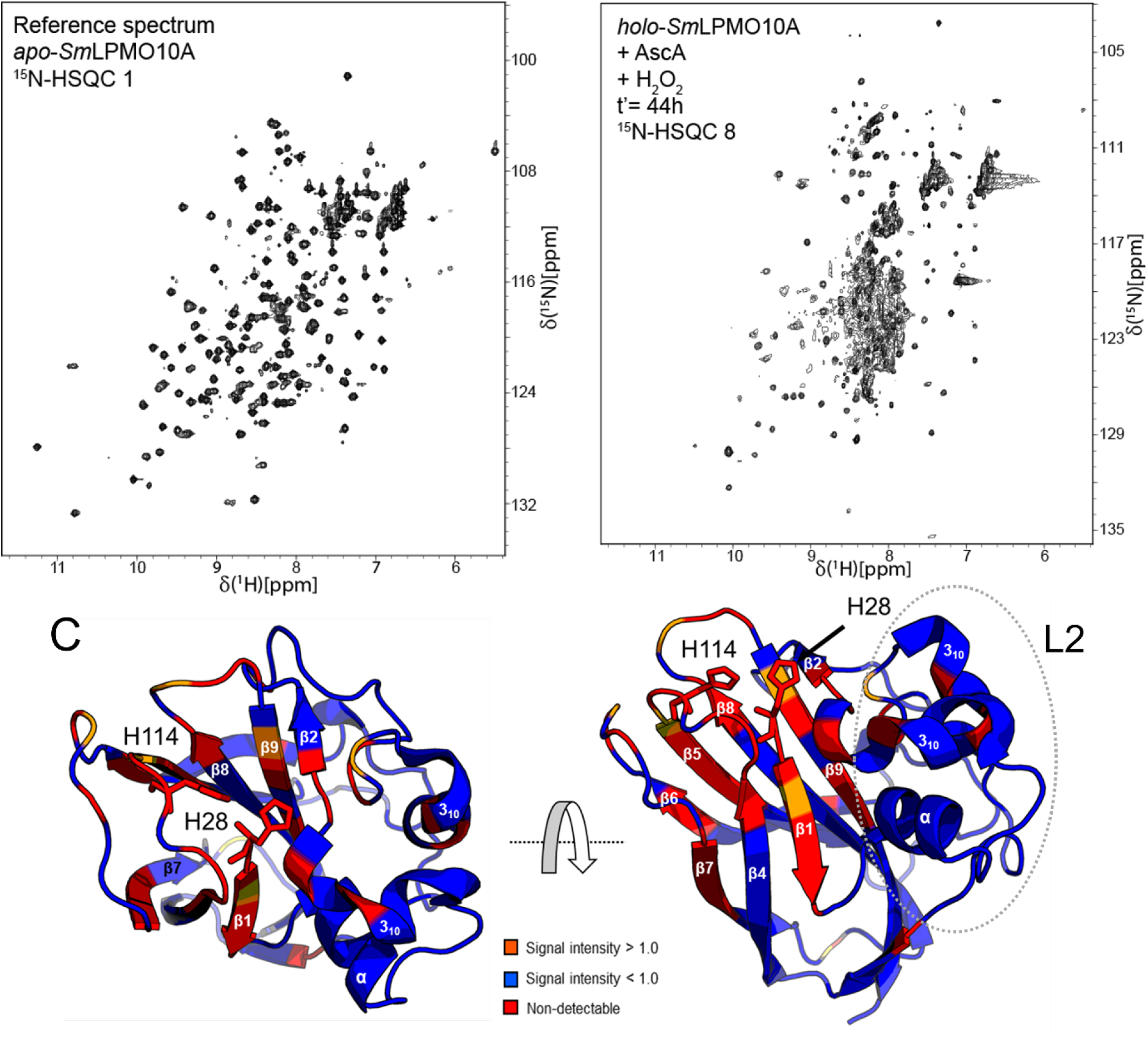
Effect of long-term (t’= 44h) incubation with H_2_O_2_. **A)** The reference ^15^N-HSQC spectrum of *apo*-*Sm*LPMO10A. The ^1^H-^15^N-signals in the spectrum are narrow and dispersed (indicated by the arrows) in both dimensions, indicating that *Sm*LPMO10A is correctly folded. **B)** The ^1^H-^15^N-HSQC spectrum of *holo*-*Sm*LPMO10A that was incubated for 40 hours with 10 mM ascorbic acid and then for 44 hours with freshly added 5 mM ascorbic acid and 2 mM H_2_O_2_ (treatment series 1). Signs of protein degradation, namely reduced signal dispersion (signals aggregating between 8 and 9 ppm) and increased line broadening, are visible. **B)** Changes in signal intensities in response to the 44-h incubation with ascorbic acid and H_2_O_2_ highlighted on the structure of *Sm*LPMO10A (PDB ID: 2BEM). An increase in signal intensity is shown in orange, while a decrease is shown in blue. Non-detectable residues are shown in red.

### CD spectroscopy reveals conformational rearrangement of aromatic residues near the active site in response to ascorbic acid

CD spectroscopy in the far UV range (185-260 nm) was used to investigate changes in the overall structure of holo-SmLPMO10A in response to incubation with 10 mM ascorbic acid in the absence of chitin (same conditions as in treatment series 1), while UV absorbance at 250 nm was used to monitor ascorbic acid consumption ^55^.

The CD spectrum of *apo*-*Sm*LPMO10A was in agreement with previous findings ^54^ (Figure S12), showing a negative maximum at 219 nm, and a zero-ellipticity crossover at 214 nm indicative of a structure with a high degree of β-strands^57^. The spectrum also featured a negative band between 185-187 nm indicating some random coils^57^, and a positive maximum at 232 nm. The positive maximum signal at 232 nm is considered to arise from π-π^*^ excitation coupling between the side-chains of tryptophan, phenylalanine, and tyrosine residues that are less than 1 nm apart ^58^ and has been shown to disappear when *Sm*LPMO10A is unfolded ^56^. Indeed, *Sm*LPMO10A contains a cluster of aromatic residues just beneath the copper site (Figure S12). The spectrum for the *holo*-enzyme showed only minor changes relative to the apo-enzyme, which can be due to the expected rigidification at the copper site (Figure S13) ^9^.

Addition of 10 mM ascorbic acid resulted in increased intensity of the positive maximum at 233 nm and appearance of a second positive band at 240-255 nm for both *holo-Sm*LPMO10A (Fig. 6) and catalytically inactive *apo-Sm*LPMO10A (Fig. S14). Control spectra of ascorbic acid alone (Figure 6 and S14) showed that these spectral changes observed immediately after adding ascorbic acid are to a large extend due to absorbance of ascorbic acid.

**Figure 6:**
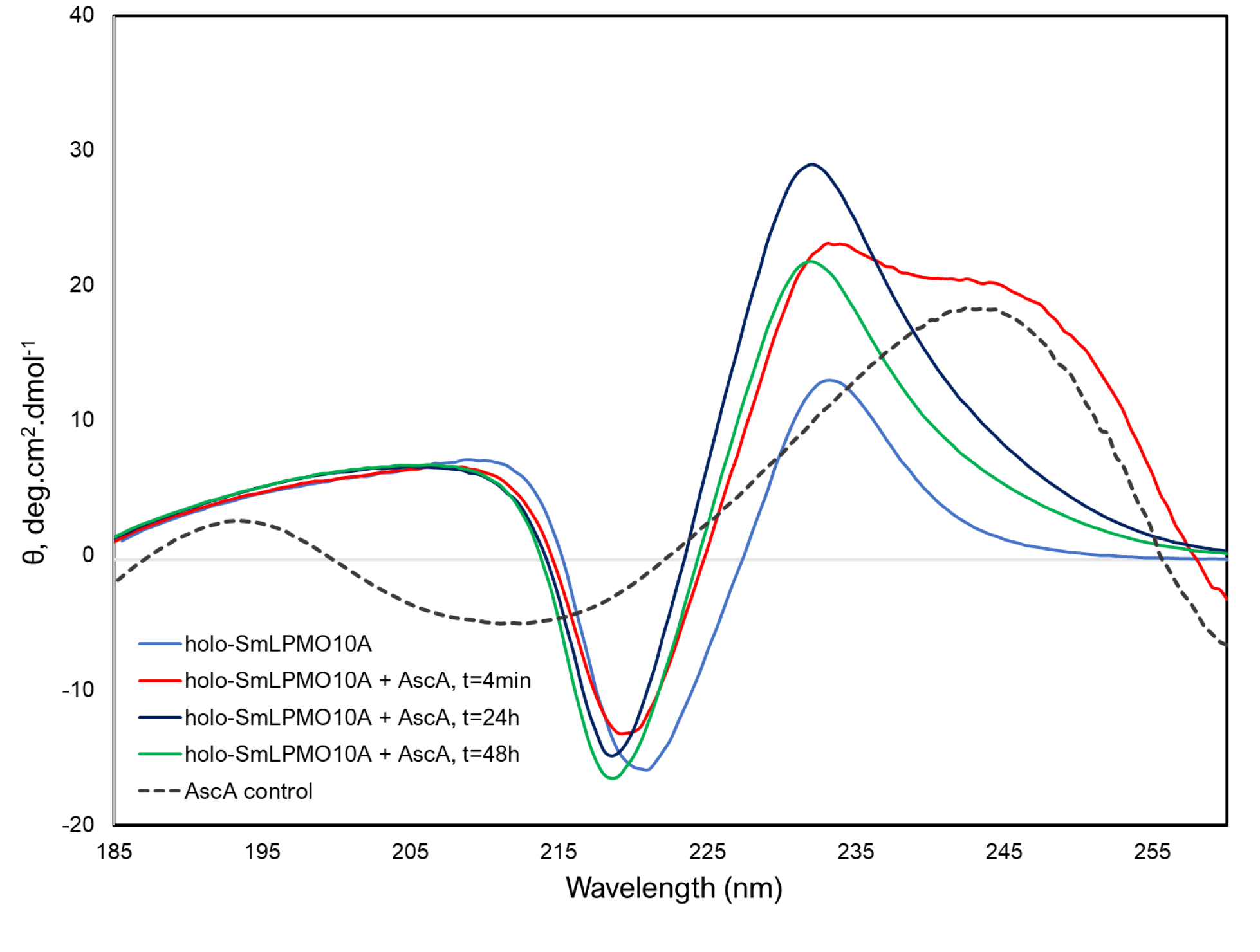
Far UV CD analysis of *holo*-*Sm*LPMO10A. The graph shows a spectrum of *holo*-*Sm*LPMO10A and spectra of *holo*-SmLPMO10A incubated with 10 mM ascorbic acid for 4min, 24h, and 48h, in the absence of chitin, at 23 °C. The protein concentration in all analyzed samples was 2.5 μM and the protein was dissolved in acetate buffer (25 mM sodium-acetate, 10 mM NaCl, pH 5.0). The dashed black curve shows a spectrum recorded for 10 mM ascorbic acid in the same buffer.

Interestingly, only in the case of *holo*-*Sm*LPMO10A, the positive maximum at ~233 nm remained increased (Figure 6), even after all ascorbic acid had been consumed (Figure S15). In contrast, the spectrum of *apo*-*Sm*LPMO10A at t= 48h after addition of ascorbic acid was very similar to the spectrum recorded after 4min (Figure S14). These findings show that the addition of ascorbic acid leads to changes in the structure of *holo-Sm*LPMO10A that change the orientation of aromatic side chains with respect to each other.

### Backbone Dynamics in the fast exchange regime

No conclutions regarding change in the mobility of *holo*-SmLPMO10A on the pico- and nanosecond time scales was reaced based on the recorded heteronuclear {^1^H}-^15^N NOEs, *T_1_*- and *T_2_*-relaxation time experiments, as shown in Figure S16, after addition of ascorbic acid. Clear changes in the structural flexibility of *holo*-*Sm*LPMO10A would be observed in the same area of the structure in all three experiments. As this was not the case, no definitive conclusion about mobility at the pico-and nanosecond timescales could be obtained.

### Chitin protects *Sm*LPMO10A against oxidative damage

The impact of the substrate was investigated by pre-incubating *holo*-*Sm*LPMO10A with β-chitin fibers for 1h before adding ascorbic acid and, subsequently, fresh ascorbic acid and H_2_O_2_ (treatment series 2). As in treatment series 1, ^15^N-HSQC spectra were recorded at varying time points (see Figure 2 for an overview). In these experiments the binding of the LPMO to chitin is expected to result in reduced signal intensities, as the substrate-bound LPMO is undetectable by solution state NMR. Consequently, the observations described below concern the non-substrate-bound fraction of the LPMO.

Changes indicative of large structural changes and increased structural flexibility were observed in the recorded ^15^N-HSQC spectrum immediately (t= 0h) after adding ascorbic acid in the presence of chitin (Figure S17, ^15^N-HSQC 10). Compared to *apo*-*Sm*LPMO10A, many residues showed narrower line widths, observed as an increase in the intensity of their ^1^H-^15^N signals (Figure 7B). The affected residues were similar to those shown in Fig. 4A (i.e., effects of ascorbic acid in the absence of chitin), but the total number of affected residues was slightly higher (Figure S18 and S19). The affected residues include E60, H114, W178, D182 and F187, found near the bound copper (Figure 7A), and the 150-160 region which comprises a loop adjacent to the loop with H114. Structural changes were also reflected by CSP >50 Hz observed for residues near the copper active site, and residues G29, K63, and A181 becoming non-detectable (Figure S18). As a whole, these combined changes seem to point at a high degree of structural changes in the ascorbic acid exposed, copper containing enzyme, as also seen in Treatment series 1.

**Figure 7:**
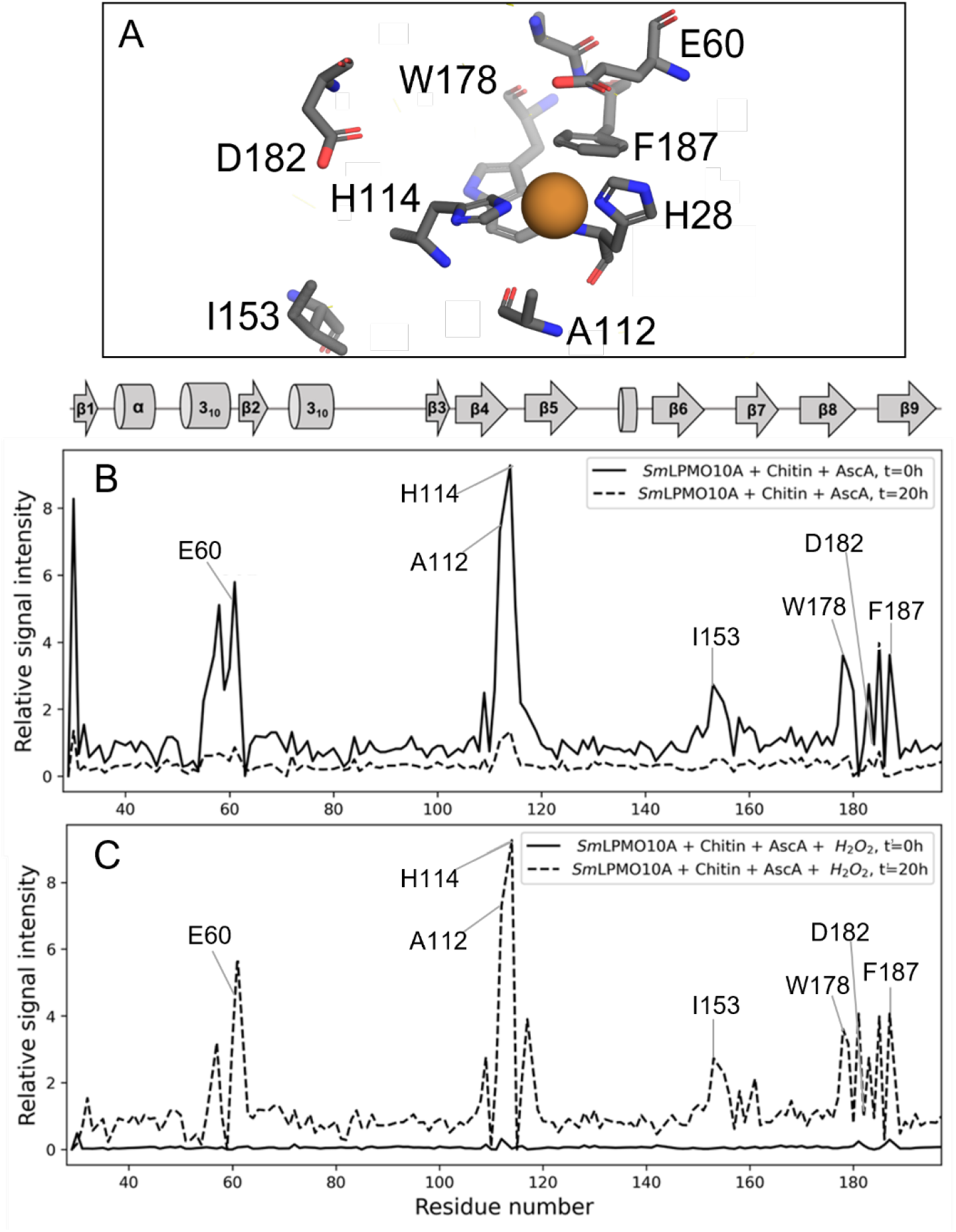
The effect of adding ascorbic acid in the presence of substrate. **A)** Stick representation of active site residues with Cu^1+^ bound (PDB 2BEM). **B)** Plots showing the relative signal intensity in response to addition of, first, 10 mM ascorbic acid (at t= 0h and t= 20h). **C)** Plots showing the relative signal intensity after a 20h incubation periode with 10 mM ascorbic acid, and, later, 5 mM fresh ascorbic acid and 2 mM H_2_O_2_ to *holo*-*Sm*LPMO10A in the presence of milled β-chitin particles with a diameter of ~ 0.5 mm (t’ =0h and t’= 20h). The relative signal intensities were calculated by taking the absolute values of the intensities of residues in each ^15^N-HSQC spectrum (^15^N-HSQCs 10–13) and dividing these by the corresponding intensities of ^1^H-^15^N signals in the reference ^15^N-HSQC spectrum of *apo*-*Sm*LPMO10A (^15^N-HSQC 1).

In contrast to what was observed in treatment series 1 (Figure 4A), only two residues (Y30 and H114) kept displaying narrower linewidths compared with the reference spectrum of *apo*-*Sm*LPMO10A (Figure S20) at t= 20h after the addition of ascorbic acid (Figure S21, ^15^N-HSQC 11). The remaining residues showed a sharp decrease in signal intensity or became undetectable (Figure S18 and S20). The decrease in signal intensities can in part be explained by line broadening (Figure S22) resulting from loss of structural integrity, as could be expected based on the results of treatment series 1. However, comparing the average signal intensities of the recorded spectra (which differ from the local variations in signal intensities discussed above), the overall reduction is larger than what was seen in treatment series 1 (Figure 8). The most likely explanation for these findings is that ascorbic acid, i.e., reduction of the LPMO combined with the availability of in situ generated H_2_O_2_, promotes binding of the LPMO to chitin, making the enzyme undetectable by solution state NMR.

**Figure 8:**
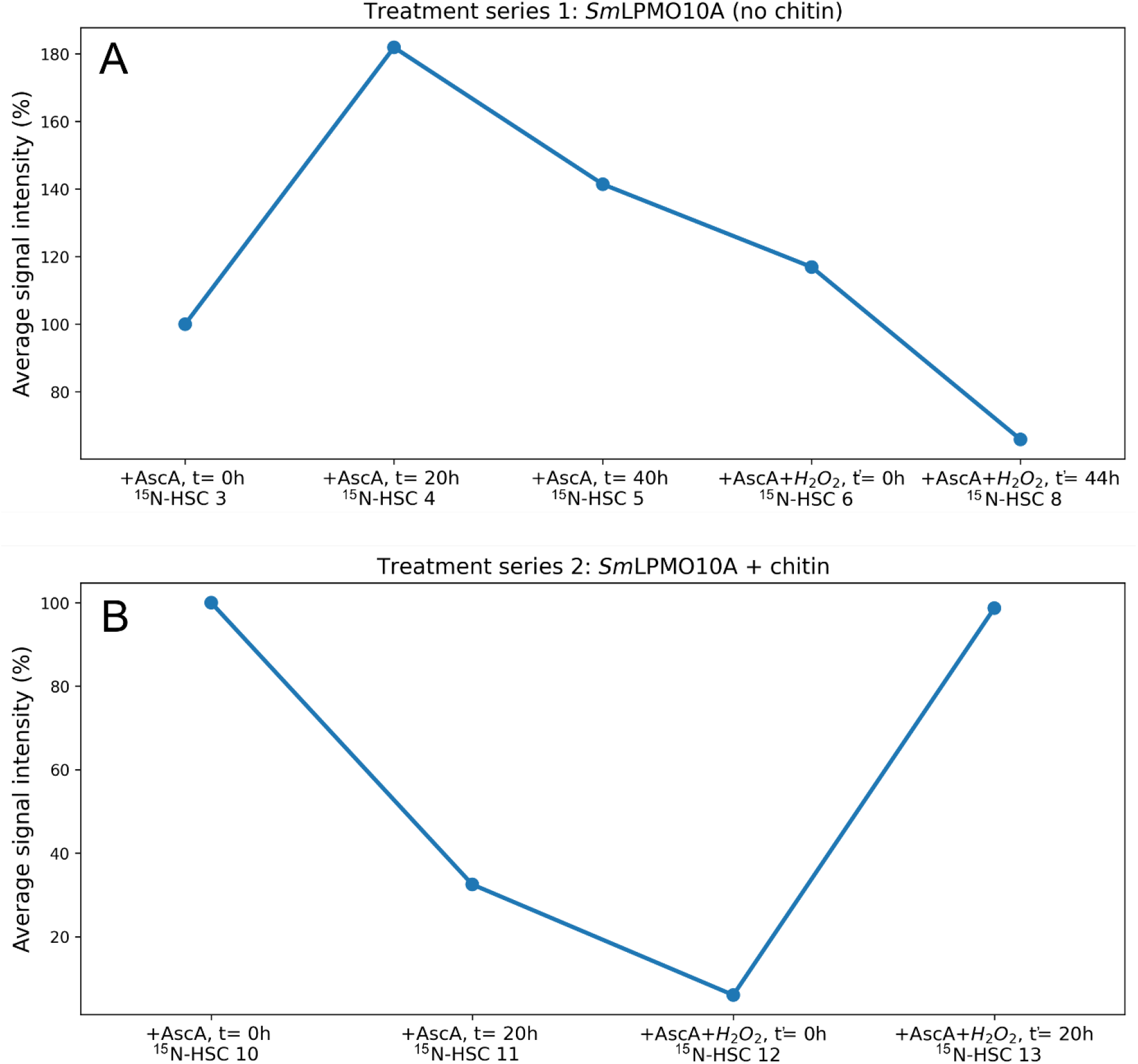
Average signal intensities for ^15^N-HSQC spectra recorded in treatment series 1 and 2. A) Average ^1^H-signal intensities in ^15^N-HSQC spectra of *holo*-*Sm*LPMO10A recorded in treatment series 1 (^15^N-HSQCs 3–8) showing the response to incubation with 10 mM ascorbic acid alone, and, subsequently, with 5 mM ascorbic acid with 2 mM H_2_O. B) Average ^1^H-signal intensities in ^15^N-HSQC spectra recorded of *holo*-*Sm*LPMO10A in treatment series 2 (^15^N-HSQCs 10–13) showing the response to incubation with 10 mM ascorbic acid alone, and, subsequently with 5 mM ascorbic acid and 2 mM H_2_O_2_, in the presence of milled β-chitin particles with a diameter of ~ 0.5 mm.

Underpinning the very different situation in the presence of substrate, activity measurements showed that, in the presence of the substrate the LPMO was still active after 20 h of incubation with ascorbic acid and chitin. In contrast, in treatment series 1, addition of ascorbic acid led to almost immediate enzyme inactivation. Of note, the activity assay does not allow a full quantitative description of a gradual decrease in catalytically competent LPMOs, since catalysis in the activity assay is to a large extent limited by access to in situ generated H_2_O_2_ and not only related to the amount of catalytically competent enzyme. It is thus possible to reconcile the observations that, on the one hand, a fraction of the LPMOs is being damaged, as suggested by the changes observed immediately after adding ascorbic acid, while, on the other hand, a fraction of the LPMOs remains active.

Addition of fresh ascorbic acid together with H_2_O_2_ was expected to boost the ongoing chitin degradation reaction but also considerable enzyme inactivation. Previous work has shown that high concentrations of ascorbic acid and H_2_O_2_ make the LPMO reaction very fast ^29^ while increasing the risk of oxidative damage ^32^. Initially, addition of ascorbic acid and H_2_O_2_ (Figure S23, ^15^N-HSQC 12) led to a further reduction of the average signal intensity (Fig. 8B) and left 9 residues undetectable (Figure S18). This is likely due to a combination of increased substrate binding and loss of signal due to structural deterioration of soluble enzyme. Upon addition of ascorbic acid and H_2_O_2_ the number of residues with large CSPs (50> Hz) also grew (Figure S18), indicating that, indeed, structural changes were taking place.

Interestingly, longer incubation times (t’= 20h) with ascorbic acid and H_2_O_2_ (Figure S24, ^15^N-HSQC 13) restored the average signal intensity (Figure 8B). Furthermore, residues that were undetectable in the previously recorded spectrum (Figure S23, ^15^N-HSQC 12) reappeared and the overall resolution of the spectrum was improved, seen by well-dispersed and narrow ^1^H-^15^N-signals (Fig. 9).

**Figure 9:**
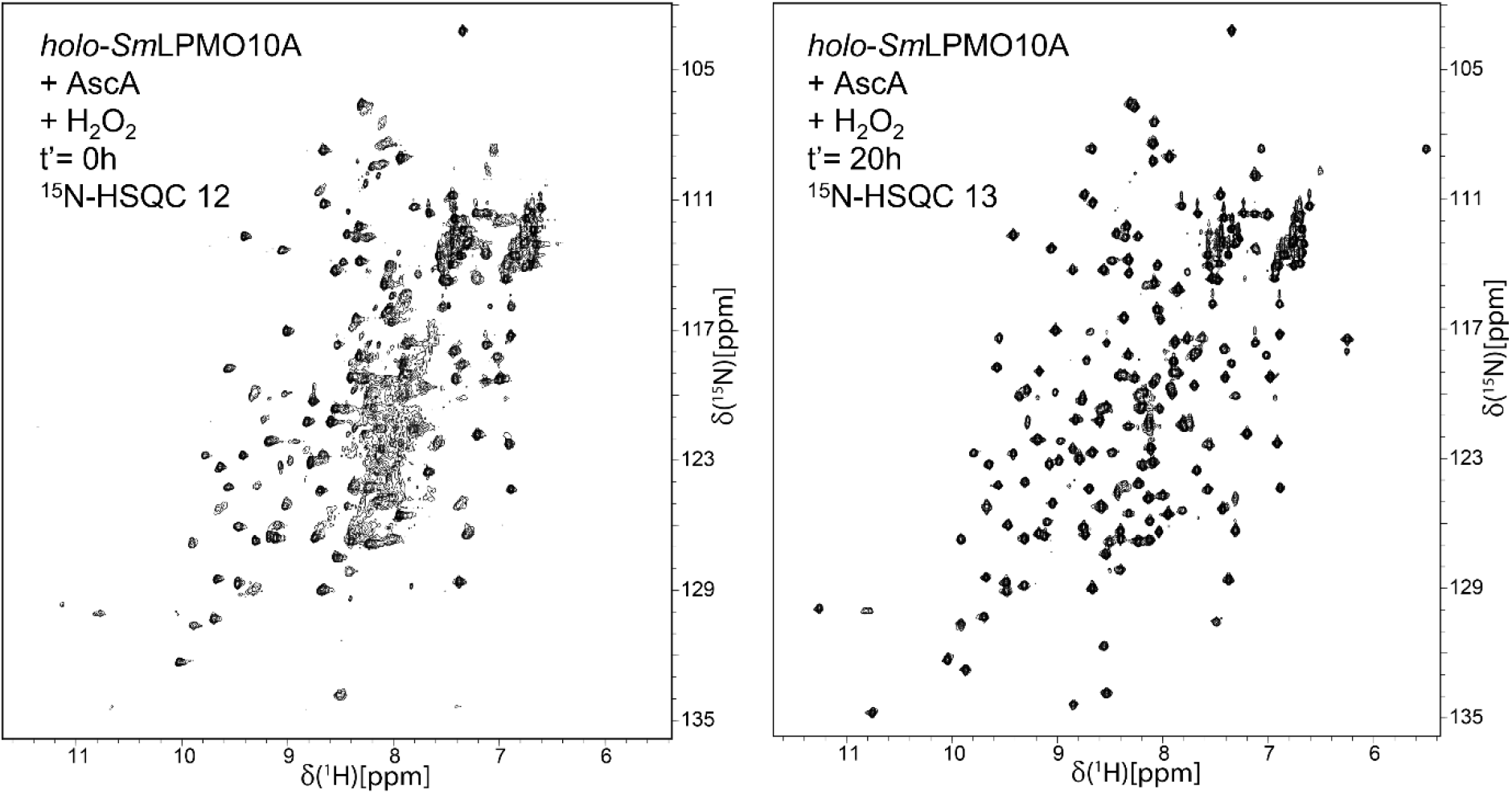
^15^N-HSQC spectra of *holo*-*Sm*LPMO10A first incubated with 10 mM ascorbic acid β-chitin, and then supplemented with 5 mM ascorbic acid, 2 mM H_2_O_2_. Spectra were recorded immediately after the addition of ascorbic acid and H_2_O_2_ (t’= 0h) and after 20 hours (t’= 20h).

Despite the restoration of signals corresponding to the folded form of *Sm*LPMO10A, the spectrum obtained 20 h after addition of ascorbic acid and H_2_O_2_ displayed numerous indications of structural changes. First of all, multiple residues showed narrower linewidths similar to what was seen right after addition of ascorbic acid (Figure S18), again pointing at damage near the catalytic copper. In addition, there were multiple CSPs >50Hz, particularly in the environment of the copper site (Figure S18). Nevertheless, the protective effect of the substrate against oxidative self-inactivation is evident when comparing the sample exposed to ascorbic acid and H_2_O_2_ in treatment series 1 and 2 (^15^N-HSQCs 7 and 13). In treatment series 2, no evidence of structural unfolding could be observed, with ^1^H-^15^N signals in the spectrum remaining dispersed and narrow (Figure 9B) and with residues staying detectable. Figure 10 illustrates the large difference in the number of residues ending up as being undetectable between treatment series 1 and 2.

**Figure 10:**
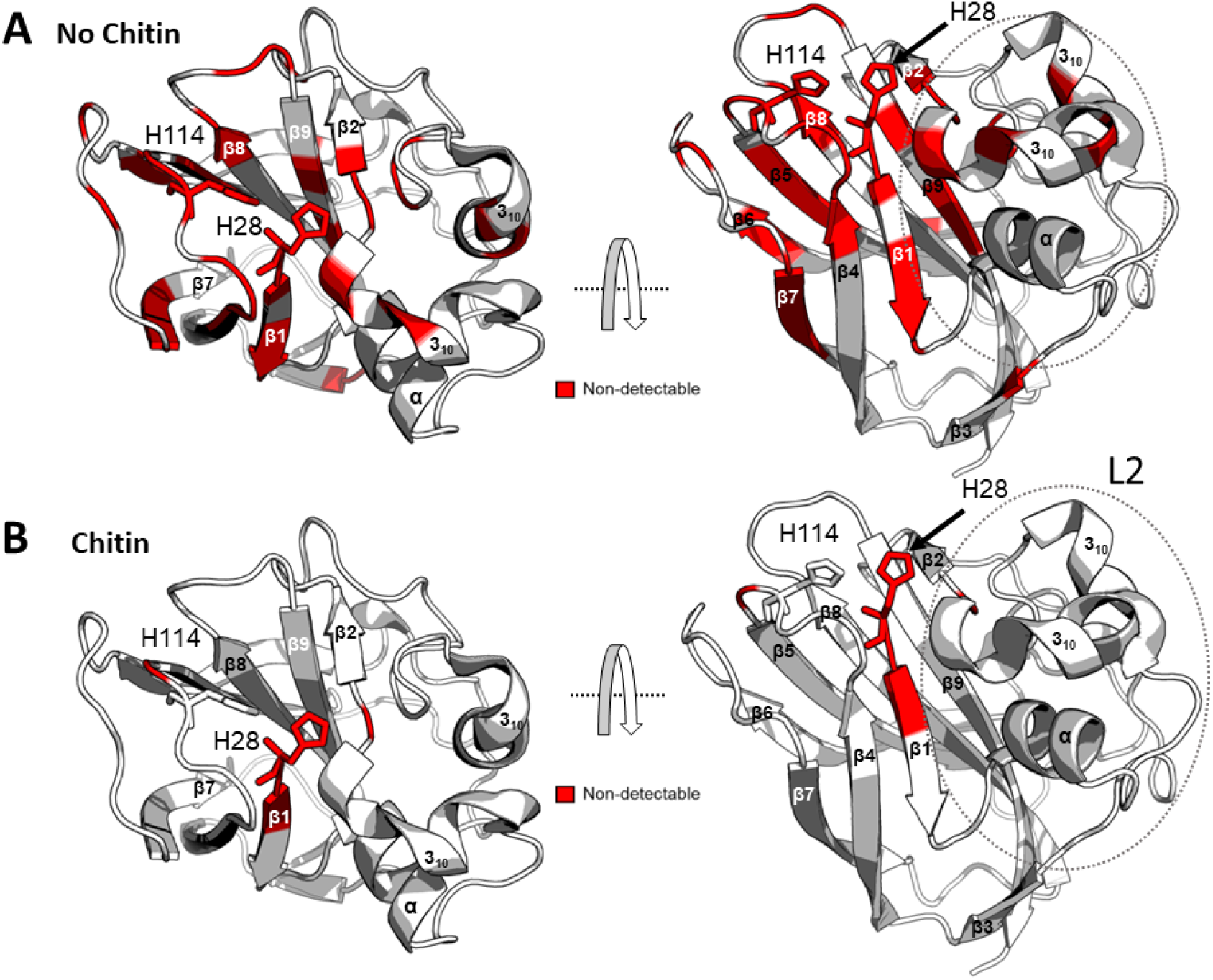
Comparison of undetectable residues in response to incubation of *holo*-*Sm*LPMO10A with 5 mM ascorbic acid and 2 mM H_2_O_2_ under aerobic conditions and after a preceding incubation with only 10 mM ascorbic acid, in **A)** the absence (treatment series 1) or **B)** presence (treatment series 2) of chitin. Residues whose signals disappeared after a 20h of incubation with ascorbic acid and H_2_O_2_ are highlighted in red on the structure of *Sm*LPMO10A (PDB ID: 2BEM).

To verify the results of treatment series 2, treatment series 3 was performed with minor differences; the pre-incubation period with chitin prior to addition of ascorbic acid was extended to 24 h. ^15^N-HSQC spectra were recorded over a longer period, compared to treatment series 2 (0–78 h with ascorbic acid alone, followed by 0 – 48 h after addition of fresh reductant and H_2_O_2_: Figure S25–32,^15^N-HSQCs 14–21). When monitoring the effect of adding ascorbic acid alone over this long time, indications of protein degradation became increasingly evident (Figure S33). Furthermore, the average signal intensity decreased with time and by t= 78h the signals of many residues were undetectable (Figure S33). The addition of fresh ascorbic acid and of H_2_O_2_ (t’= 0h) to treatment series 3 resulted in signals corresponding to the active form of *holo*-*Sm*LPMO10A to seemingly reappear (Figure S30,^15^N-HSQCs 19), as evidenced by missing residues reappearing and restoration of the signal dispersion in the spectrum (Figure S34) and an overall increase in the average signal intensities (Figure S35), although it was less pronounced than in treatment series 2. The latter is compatible with the prolonged preceding incubation with ascorbic acid only, which would lead to a larger degree of enzyme damage, as was indeed observed (Figure S35). Longer incubation (t’ > 0h) resulted in a drastic decrease in the average signal intensity due to the enzyme suffering further oxidative damage.

## Discussion

LPMOs are prone to oxidative self-inactivation caused by off-pathway reactions involving the reduced LPMO and oxygen or H_2_O_2_^15,35^. Reactions with the oxygen co-substrate generate reactive oxygen species, including hydroxyl radicals, that may emerge upon homolytic cleavage of H_2_O_2_^15,26,59^ As the substrate helps confining the reactive oxygen species, off-pathway reactions are exacerbated in the absence of substrate^15,29,33,60^. The chemical details of the destructive redox chemistry that happens in LPMO active sites remain partly unknown. It has been shown that the copper-binding histidine’s and nearby aromatic residues are particularly vulnerable^15^. In experiments with ascorbic acid only (as in the present study) H_2_O_2_ is generated through the reductant oxidase activity of the LPMO^36^ and through abiotic oxidation of the reductant^16^. In reactions with bacterial family AA10 LPMOs, such as *Sm*LPMO10A, such *in situ* production of H_2_O_2_ will be rather slow^61^, meaning that there is a large difference between the incubations with and without H_2_O_2_ added.

NMR enables real-time observation of changes in the overall structure of proteins and of changes in individual residues, while at the same time providing dynamic information. NMR thus provides the possibility to gain insight into the structural changes that accompany oxidative self-inactivation of LPMOs. In the present study, we have used ^15^N-HSQC ‘fingerprint’ spectra as reporters of structural changes during treatment series (summarized in Figure 2) that were designed to induce oxidative damage. Importantly, the set-up of this study was such that we observed both autocatalytic damages, as one would expect during the initial phase of incubating with ascorbic acid, and more general damage by reactive-oxygen species, which happens when the LPMO is exposed to high concentrations of hydrogen peroxide in the presence of transition metals. Changes in the structure and dynamics of *Sm*LPMO10A were monitored by CSPs and ^1^H-^15^N signal linewidths, respectively, while overall structural integrity was monitored by the overall chemical shift dispersion. It is important to note that ^15^N-HSQC spectra only reveal changes that affect the backbone H^N^ and N atoms of *Sm*LPMO10A. Further insight into the changes induced by ascorbic acid was achieved using CD spectroscopy.

In agreement with previous findings^9^, signals belonging to residues near (~10 Å) the copper active site became undetectable due to PRE caused by binding of Cu(II) to *apo*-*Sm*LPMO10A. Addition of ascorbic acid resulted in the reappearance of signals near the copper site due to reduction of the copper to diamagnetic Cu(I). Importantly, oxidative damage to the copper-coordinating histidines, H28 and H114, likely leads to release of the copper atom from the active site^53^. Thus, in all treatment series, the continuous persistence of signals sensitive to PRE may mean that the copper stays reduced but also that the copper dissociates from the enzyme.

In this study, large amounts of ascorbic acid were added, which in addition to reducing the LPMO and relieving PRE, would result in *in situ* generation of H_2_O_2_, causing autocatalytic inactivation of the reduced enzyme. Data from treatment series 1, showed that addition of ascorbic acid to Cu(II) *holo*-*Sm*LPMO10A caused residues near the copper active site to display CSPs >50Hz and narrower linewidths, when comparing with *apo-Sm*LPMO10A (Figure 3). Both effects indicate changes in the backbone structure of *Sm*LPMO10A. Minor conformational rearrangements and reduced conformational flexibility brought about by copper binding have been reported in the literature^9,62^. However, while copper binding is expected to result in broadening due to reduced conformational flexibility, the presence of ascorbic acid led to the opposite, namely narrowing of linewidths for residues near the copper center. This narrowing could result from a slow exchange regime between multiple structural states of *holo*-*Sm*LPMO10A or result from increased local flexibility near the active site. This indicates structural changes in the catalytic center that are likely due to oxidative damage caused by an autocatalytic process, aligning well with an observed rapid decrease in catalytic activity. Longer incubations with ascorbic acid led to general line broadening, suggesting that oxidative damage propagated through the enzyme and that overall structural integrity was being lost. In accordance with these structural signs of enzyme damage, activity measurements showed that the enzyme was completely inactive after 24 h of incubation with ascorbic acid.

Conformational rearrangements in response to ascorbic acid were also observed by CD-spectroscopy. The CD-spectrum of both *apo*- and *holo*-*Sm*LPMO10A showed an unusual positive maximum at 233 nm, which can be explained by π-π^*^ excitation coupling between aromatic side-chains ^58^, which causes a strong negative peak at 213 nm and positive peak around 230nm ^63–65^. The increased intensity of the positive maximum at 233 indicates a stronger contributions from π-π^*^ excitation couplings between aromatic side-chains following incubation with ascorbic acid ^64,65^. *Sm*LPMO10A contains at least three pairs of aromatic residues able to exhibit π-π^*^ excitation coupling: Y30-Y39, W108-W119, and W178-F187. In addition, excitation coupling between W119 and W178 or F187 is possible (Figure S12). Based on the NMR data, conformational changes in these aromatic pairs are most likely to occur for the W119-F178 pair, since both involved residues showed CSPs >50 Hz in response to the addition of ascorbic acid (Figure 3). The pair is located directly below the copper active site (Figure S12). Interestingly, it has been speculated that conserved tyrosine and tryptophan residues near the active site are protecting LPMOs from inactivation through a ‘hole hopping’ pathways^28,41,66^ and the changes observed here through CD measurements and NMR could relate to such pathways. In this respect, it is worth noting that CSPs > 50 Hz were also observed for residues Y30 and W108, meaning that conformational rearrangements of these residues also may contribute to increased π-π^*^ excitation couplings. The residue Y39 was not assigned, meaning that no structural information about this residue was obtained, while W119 did not show a significant CSP.

After the incubation with ascorbic acid and enzyme inactivation, there will be free copper in the solution, that means that subsequent addition of fresh ascorbic acid and H_2_O_2_ will lead to a highly damaging environment allowing, for example, Fenton-like chemistry catalyzed by reduced free copper. Since free copper promotes oxidation of ascorbic acid, hydrogen peroxide levels will be high. This situation likely leads to massive enzyme damage which is no longer autocatalytic. Indeed, the ^15^N-HSQC spectra (Figure S7–S9, ^15^N-HSQC 6–8) showed features commonly associated with protein unfolding ^52^, which became increasingly pronounced as time progressed (Figure 5). Looking at which residues became undetectable, it would seem that β-strands 1 and 5 were particularly prone to damage and (partial) unfolding. Structural elements that seemed to remain intact include parts of the β-sandwich core and the distal parts of the *L2*-loop (~ 13 Å from the active site), as shown in Figure 5.

In treatment series 2, addition of ascorbic acid after pre-incubation with chitin (Figure 2, ^15^N-HSQC 10) was again immediately (t= 0h) accompanied by CSPs and indications of increased structural flexibility. Longer incubation with ascorbic acid led to a notable reduction in the average signal intensity, which was mainly due to binding of the LPMO to the substrate and not to structural disintegration, since signal intensities were restored later on in the treatment series, upon addition of fresh ascorbic acid and H_2_O_2_ (Fig. 8). Adding β-chitin particles to the samples creates a biphasic system where the substrate-bound *Sm*LPMO10A becomes invisible^9^. Together with activity data, the NMR data show that reduction and the presence of *in situ* generated H_2_O_2_ promote substrate binding and that such binding protects the LPMO from damage and inactivation. The signs of protein degradation, a process that eventually also will contribute to a decrease in signal intensities, observed in response to ascorbic acid originate from unbound *Sm*LPMO10A, which suffers from oxidative damage. The results of treatment series 3 led to a similar conclusion; in this case, damage during the ascorbic acid-only phase was more extensive, due to the much longer incubation time.

The addition of fresh ascorbic acid and H_2_O_2_ in treatment series 2 and 3 (i.e., with the presence of chitin) led to an increase in the signal intensities of nearly all observable residues. Signals that were unobservable before the addition of H_2_O_2_ also became visible again. Moreover, the overall appearance of the spectra in terms of signal dispersion became more similar to the spectra collected before the initial addition of ascorbic acid. The reappearance of signals for intact, reduced LPMO, are compatible with the notion that addition of H_2_O_2_ drastically speeds up the rate of chitin cleavage, which could lead to release of catalytically competent LPMO into solution, which is be visible to NMR again.

Based on these results, we propose the chain of events outlined in Figure 11. When the copper active site of *Sm*LPMO10A is reduced, the substrate affinity of the enzyme increases and the equilibrium shifts towards more protein being substrate-bound, as previously demonstrated by Kracher and colleagues ^30,37,67^. The addition of externally supplied H_2_O_2_ increases catalytic activity, resulting in substrate oxidation and the release of intact LPMO back into solution. This chain of events implies that the presence of chitin, i.e., a good substrate, protects the LPMO against self-oxidative damage, as was confirmed by activity measurements showing that loss of enzyme activity was much reduced in the reactions with chitin. It has been shown that binding of the LPMO to substrate increases the enzyme’s reactivity towards H_2_O_2_^26,35^. Thus, the presence of substrate has multiple effects. On the one hand, reduced LPMOs are removed from solution, which limits the risk of futile turnover of H_2_O_2_, which could lead to enzyme damage. On the other hand, the increased consumption of H_2_O_2_ in the enzyme substrate complex removes available H_2_O_2_ from solution, further reducing the chance of futile, potentially damaging turnover by non-substrate bound LPMOs.

**Figure 11:**
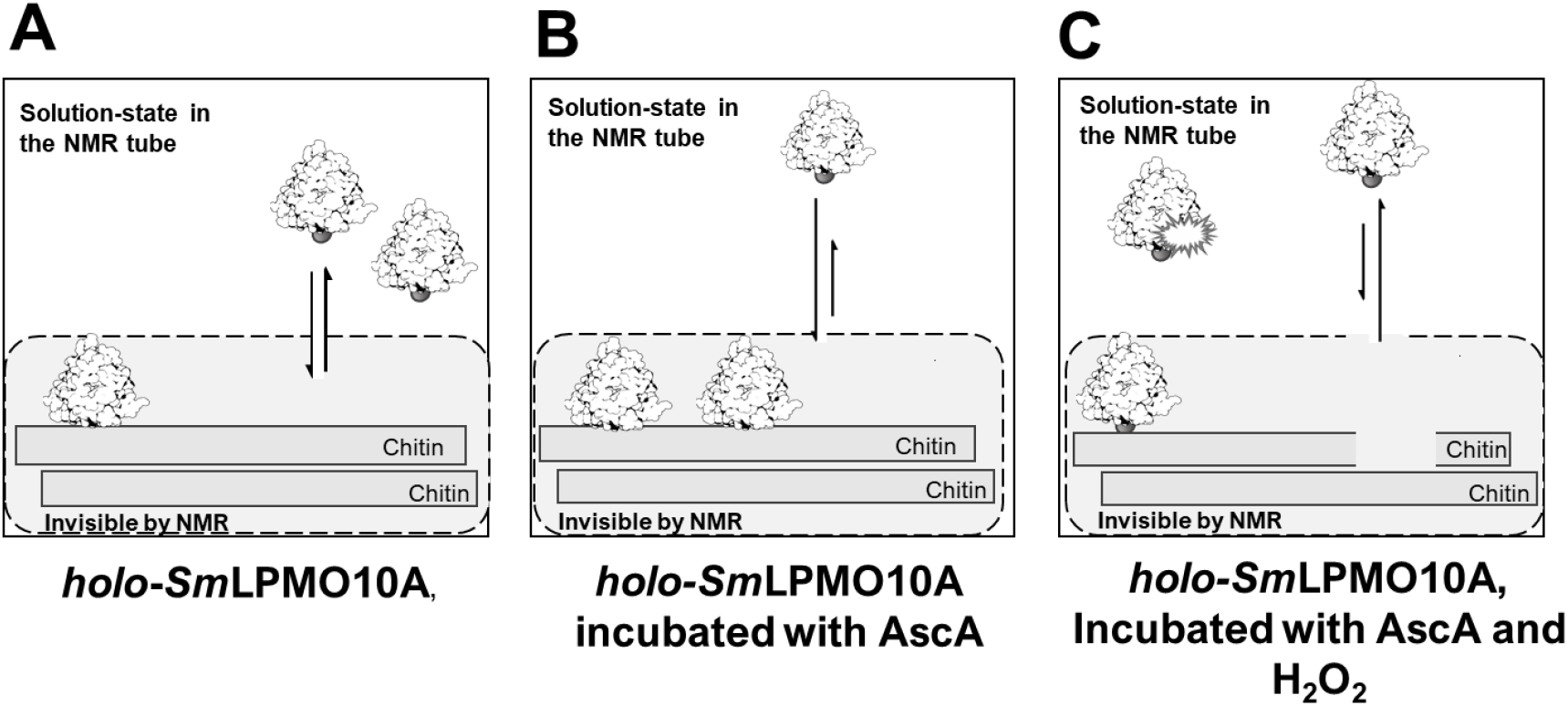
The effect of ascorbic acid and H_2_O_2_ on *holo*-*Sm*LPMO10A in the presence of chitin. **A)** *holo*-*Sm*LPMO10A in absence of ascorbic acid and H_2_O_2_; the enzyme is in the Cu(II) state **B)** *holo*-*Sm*LPMO10A incubated with ascorbic acid. Chitin is insoluble and therefore the chitin-bound LPMO is not detectable by solution-state NMR. Consequently, *Sm*LPM10A bound to the substrate will not be detected. Substrate-binding is promoted by reduction of the LPMO. **C)** The immediate effect of adding fresh ascorbic acid and H_2_O_2_ to the sample described in (B). H_2_O_2_ can diffuse into the active site of substrate-bound *Sm*LPMO10A and its presence leads to a drastic increase in the rate of chitin cleavage. Upon cleaving the substrate, *Sm*LPMO10A is released back into solution where it is detectable by solution-state NMR.

## Conclusion

Overall, by using NMR spectroscopy, CD spectroscopy, and activity assays, our results shed light on the process of oxidative self-inactivation of *Sm*LPMO10A over time. Whereas CSPs place the process of oxidative damage in a structural context, by providing direct insights into the chemical environment of the observed nuclei, changes in signal intensities indicate structural flexibility and /or that the native tertiary structure of *Sm*LPMO10A is partially lost. Using these tools, we show how oxidative damage first happens near the copper site and then propagates through the protein, and we show that chitin protects *Sm*LPMO10A from oxidative self-inactivation. Our CD studies suggest that aromatic residues in the core of *Sm*LPMO10A play a role in determining the fate of redox-active species generated at the copper site and further studies of the potential role of these residues in protecting the enzyme from autocatalytic inactivation would be of interest.

## Acknowledgements

The Norwegian Research Council funded this work through the OXYMOD (269408) and Norwegian NMR Platform (226244) projects. Gaston Courtade received funding from the Novo Nordisk Foundation through grant number NNF18OC0032242.

