## Supplementary files for "Following the fate of lytic polysaccharide monooxygenases (LPMOs) under oxidative conditions by NMR spectroscopy"

ORCIDs

Idd A. Christensen: 0000-0001-6229-2211

Vincent G. H. Eijsink: 0000-0002-9220-8743

Anton Stepnov: 0000-0002-6754-5852

Gaston Courtade: 0000-0002-1644-3223

Finn L. Aachmann: 0000-0003-1613-4663

*** To whom correspondence should be addressed.**
Finn L. Aachmann
NOBIPOL, Department of Biotechnology and Food Science, NTNU Norwegian University of Science and Technology, Sem Sælands vei 6/8, N-7491 Trondheim, Norway

Vincent G. H. Eijsink
Faculty of Chemistry, Biotechnology and Food Science, NMBU - Norwegian University of Life Sciences, 1432 Ås, Norway

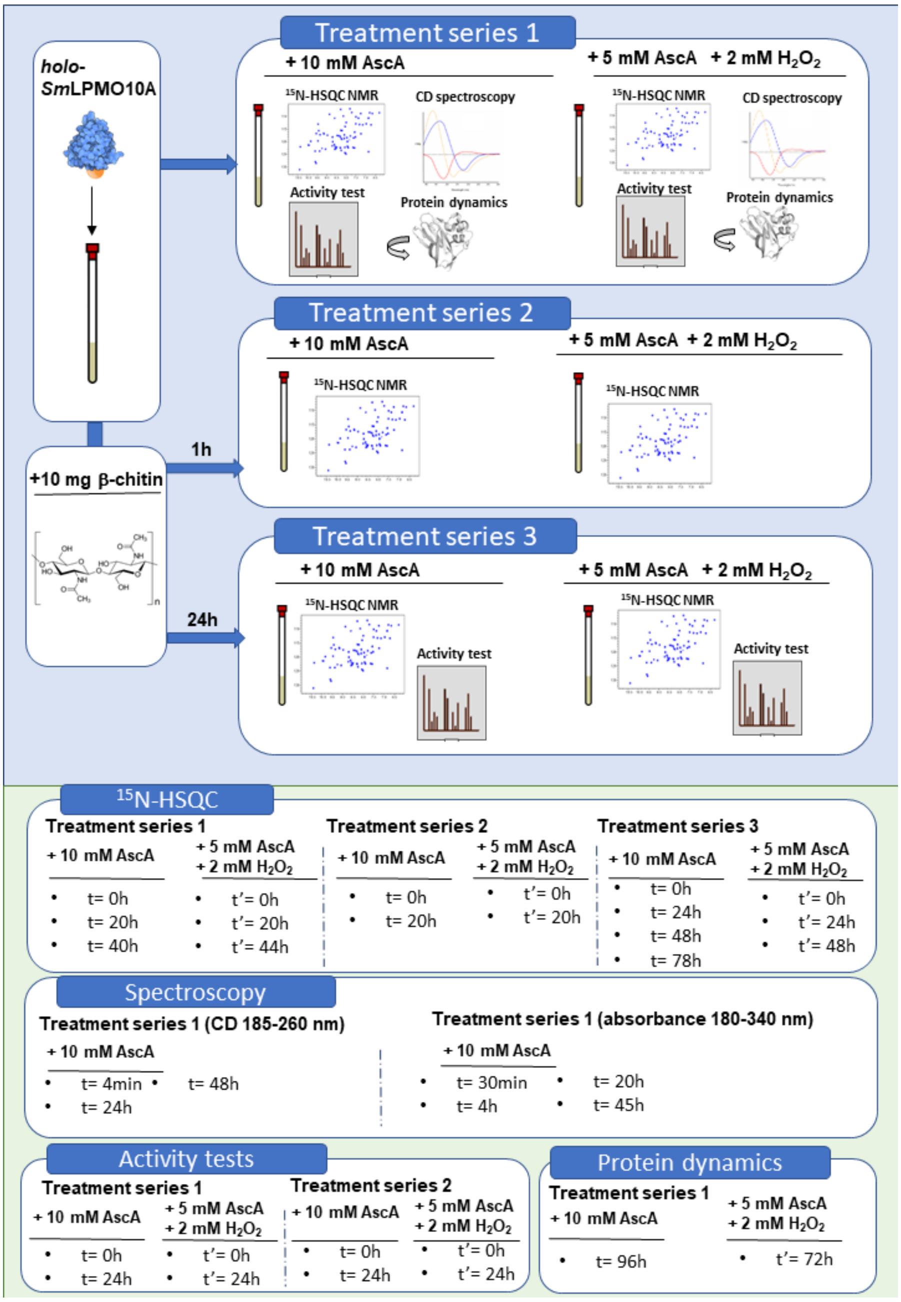


**Figure S1:** Overview of the experimental work done to observe the structural effects of oxidative conditions on *holo*-*Sm*LPMO10A. The effects of oxidative conditions were followed in three treatment series. Treatment series 1 was carried out in the absence of the substrate chitin, while *holo*-*Sm*LPMO10A was pre-incubated with 10 mg β-chitin particles for 1h and 24h in treatment series 2 and 3, respectively. Oxidative conditions were induced in all three treatment series by, first, the addition of 10 mM ascorbic acid and, later, the addition of 5 mM ascorbic acid and 2 mM $H_2O_2$. The methods used to monitor the effects of the oxidative conditions are indicated by icons for each treatment series, and the time-points for the various measurements are provided for each treatment series in the panels below. All samples were prepared in acetate buffer (25 mM sodium-acetate, 25 mM acetic acid, 10 mM NaCl, pH 5.0) with ~ 100-150 µM *holo*-*Sm*LPMO10A. Individual samples were prepared for each of the methods used in each treatment series.

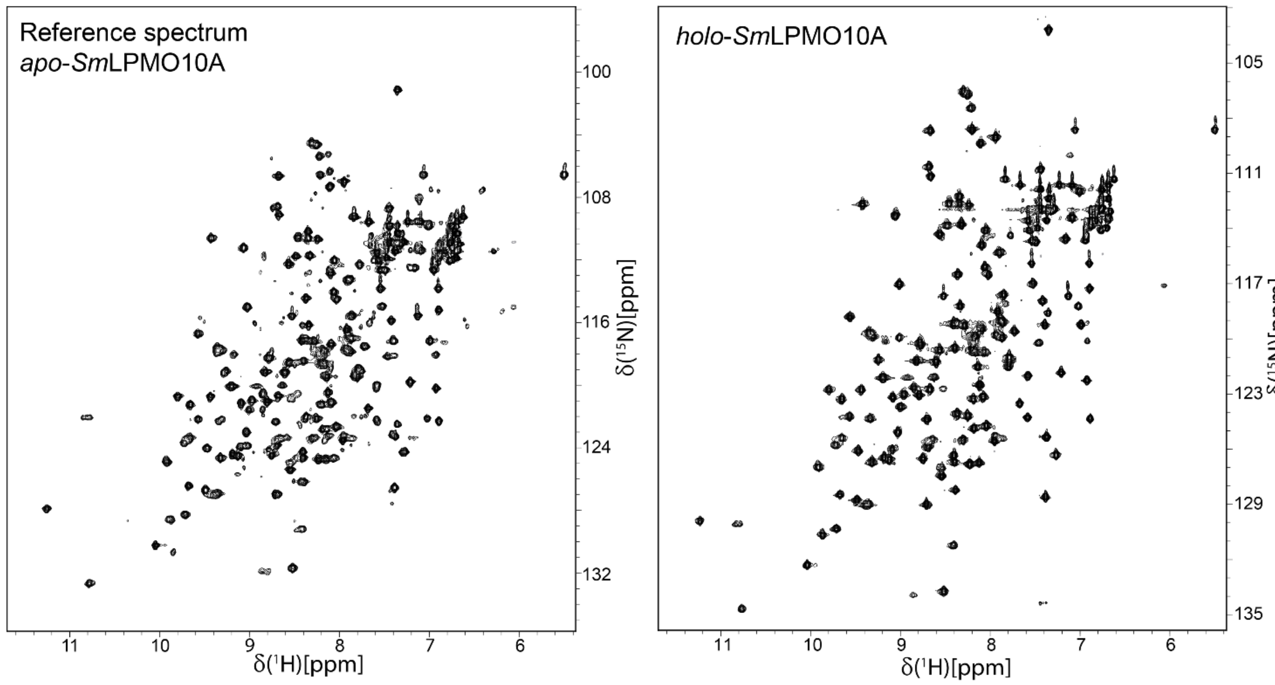


**Figure S2**: $^{15}$N-HSQC spectra of *apo-Sm*LPMO10A and *holo-Sm*LPMO10A in acetate buffer (25 mM sodium-acetate, 10 mM NaCl, pH 5.0). The $^{1}$H-$^{15}$N-signals in the spectrum are narrow and dispersed in both dimensions indicating that *Sm*LPMO10A is correctly folded. In the case of *holo-Sm*LPMO10A $^{1}$H-$^{15}$N signals of residues within ~10Å from the copper active site relax too fast to be detected in the spectrum, due to Paramagnetic Relaxation Enhancement caused by binding of Cu(II) to the active site.

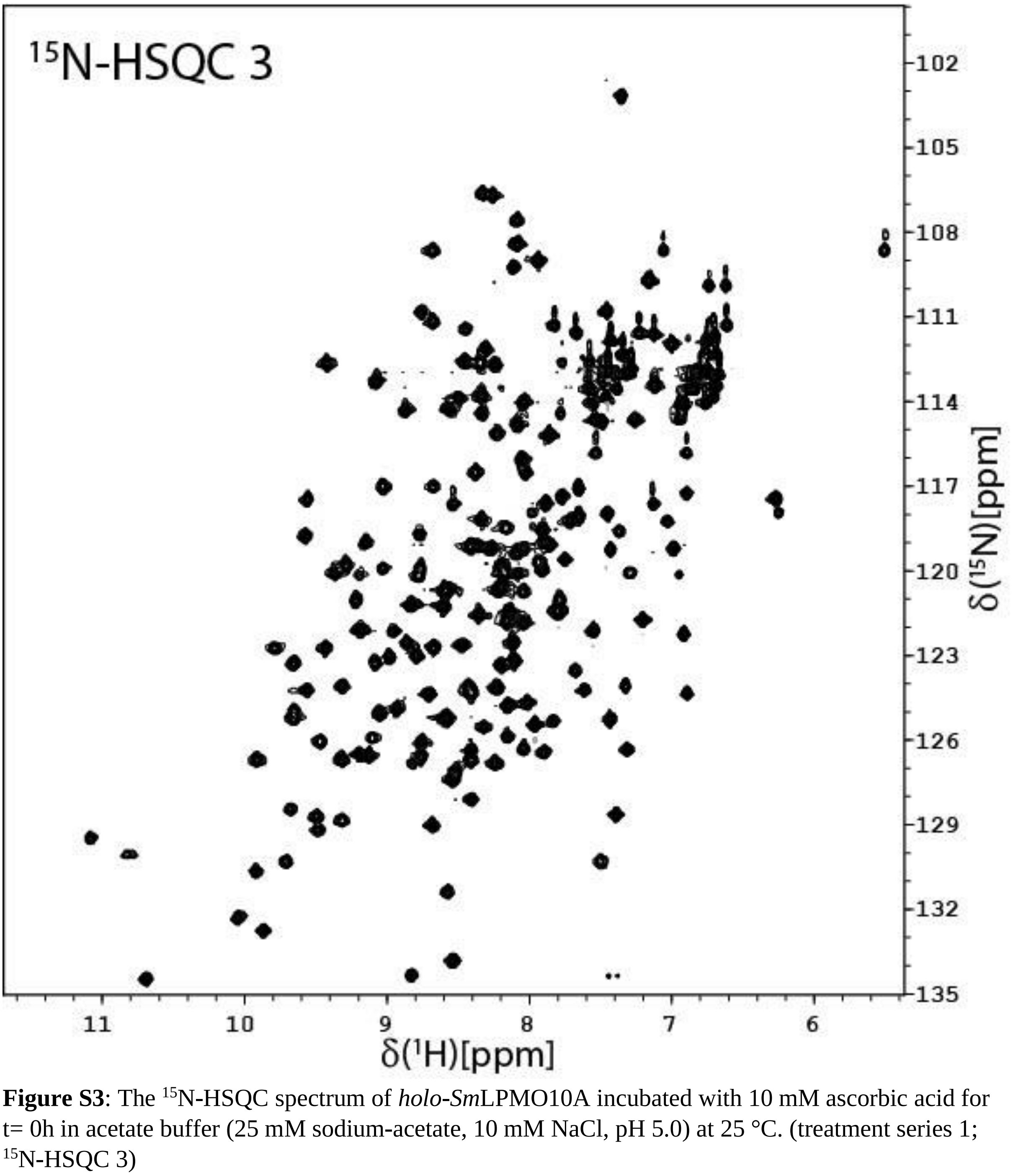


**Figure S3**: The $^{15}$N-HSQC spectrum of *holo*-*Sm*LPMO10A incubated with 10 mM ascorbic acid for t= 0h in acetate buffer (25 mM sodium-acetate, 10 mM NaCl, pH 5.0) at 25 °C. (treatment series 1; $^{15}$N-HSQC 3)

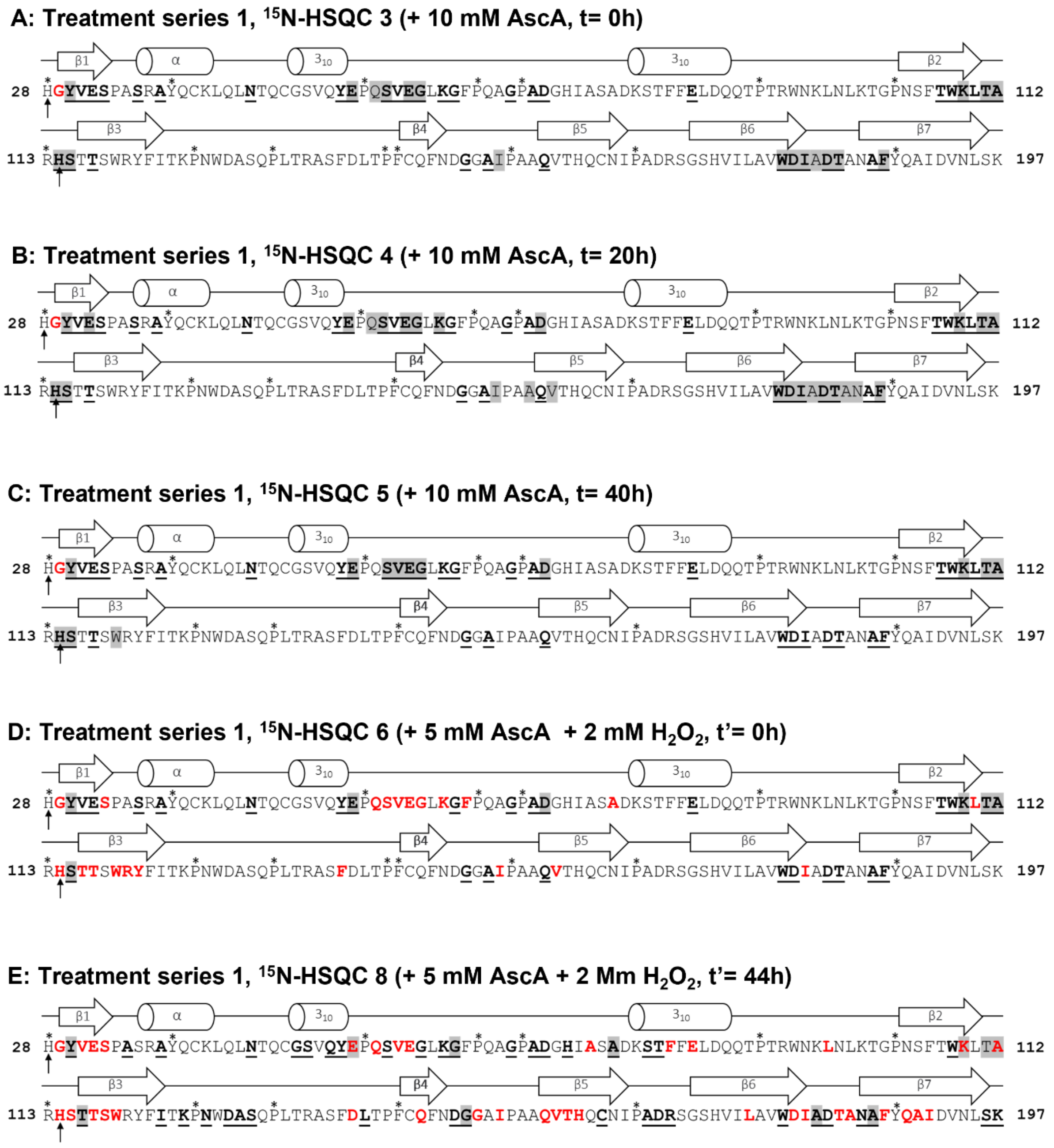


**Figure S4**: The combined structural changes to *holo*-*Sm*LPMO10A observed in treatment series 1, relative to the reference spectrum of *apo*-*Sm*LPMO10A. The active site histidines are indicated by arrows, while residues not assigned in by NMR are indicated by asterixis. Chemical shift perturbations >50Hz are shown as by bold and underlined residues, narrower linewidths (i.e increased signal intensities) are shown by residues highlighted in grey. Non-detectable $^{1}H$-$^{15}N$ signals are shown by red residues.

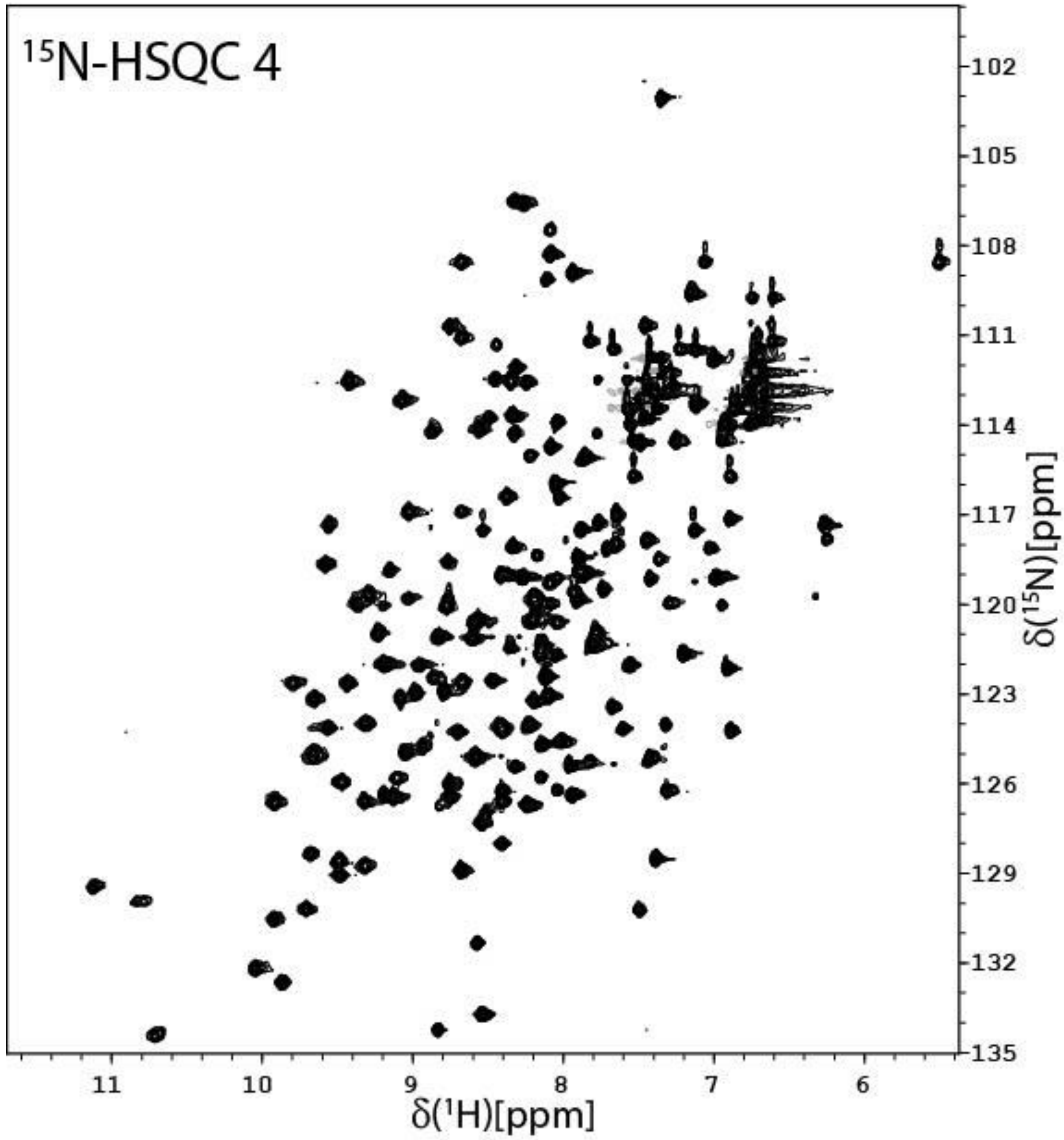


**Figure S5**: The $^{15}$N-HSQC spectrum of *holo-Sm*LPMO10A incubated with 10 mM ascorbic acid for t= 20h in acetate buffer (25 mM sodium-acetate, 10 mM NaCl, pH 5.0) at 25 °C (treatment series 1; $^{15}$N-HSQC 4).

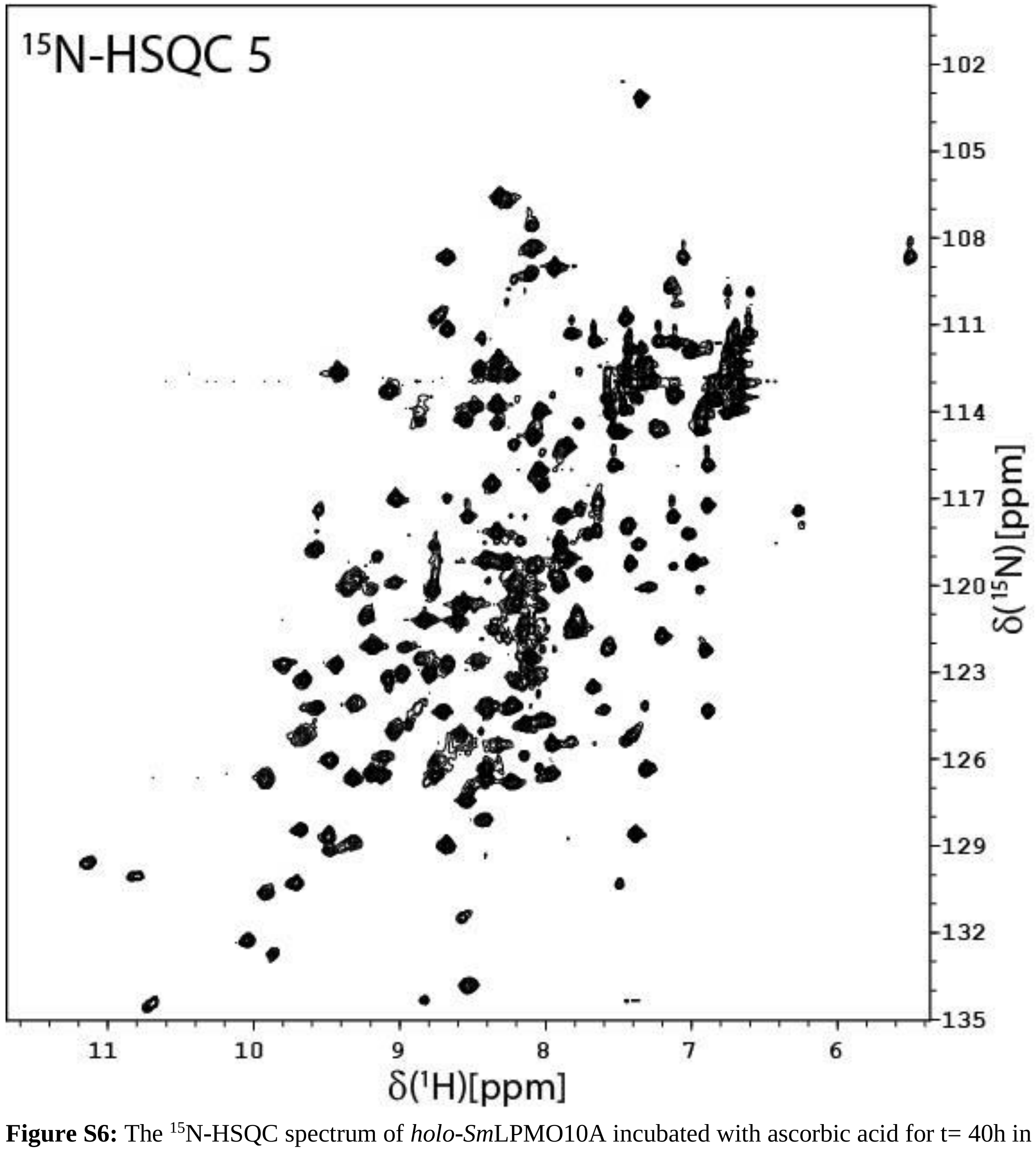


**Figure S6:** The $^{15}$N-HSQC spectrum of *holo*-*Sm*LPMO10A incubated with ascorbic acid for t= 40h in acetate buffer (25 mM sodium-acetate, 10 mM NaCl, pH 5.0) at 25 °C. (treatment series 1; $^{15}$N-HSQC 5).

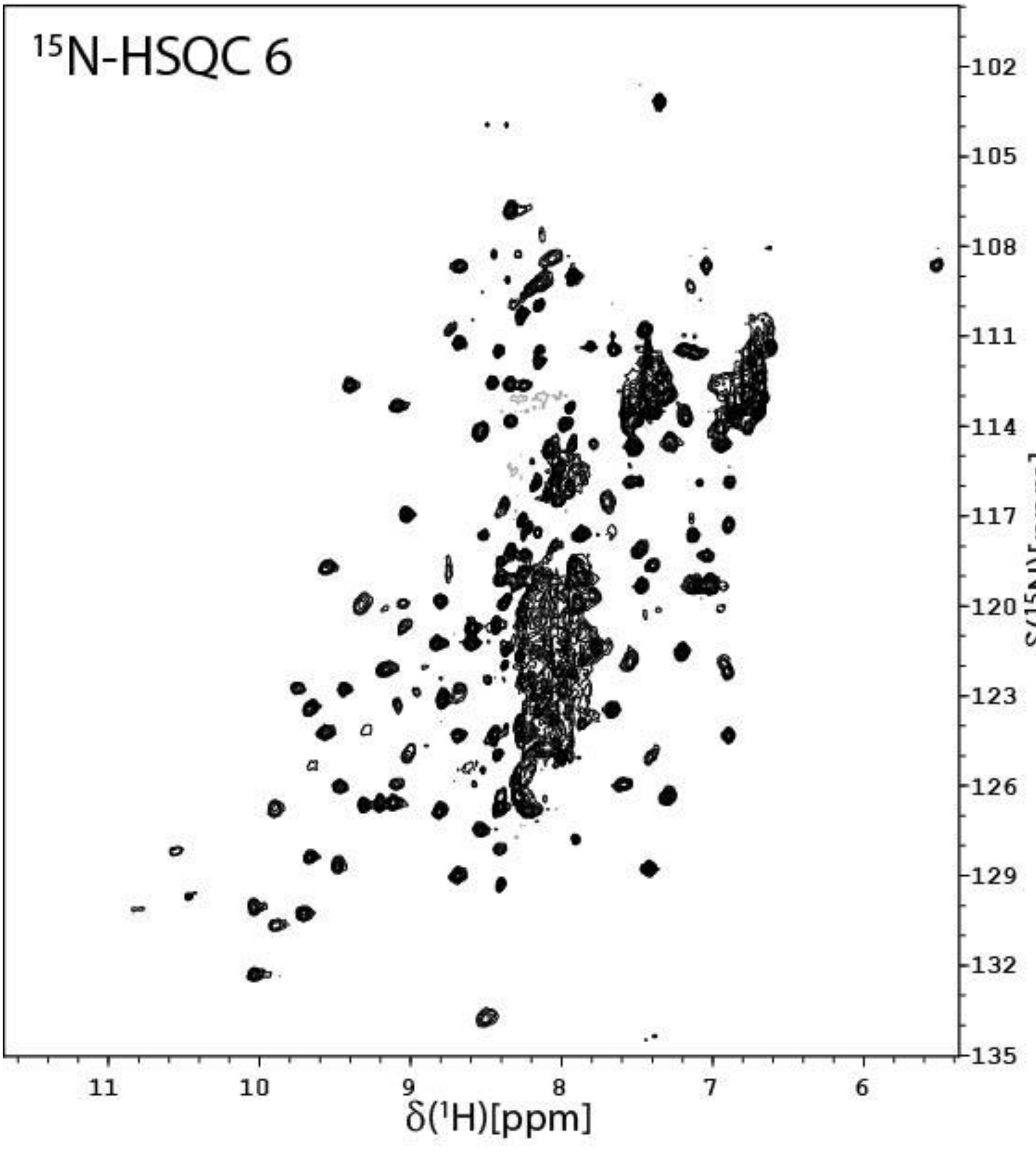


**Figure S7:** The [15]N-HSQC spectrum of *holo*-*Sm*LPMO10A incubated with 10 mM ascorbic acid for 40h and, subsequently, with 5 mM ascorbic acid and 2 mM $H_2O_2$ for t'= 0h in acetate buffer (25 sodium-acetate, 10 mM NaCl, pH 5.0) at 25 °C (treatment series 1; [15]N-HSQC 6).

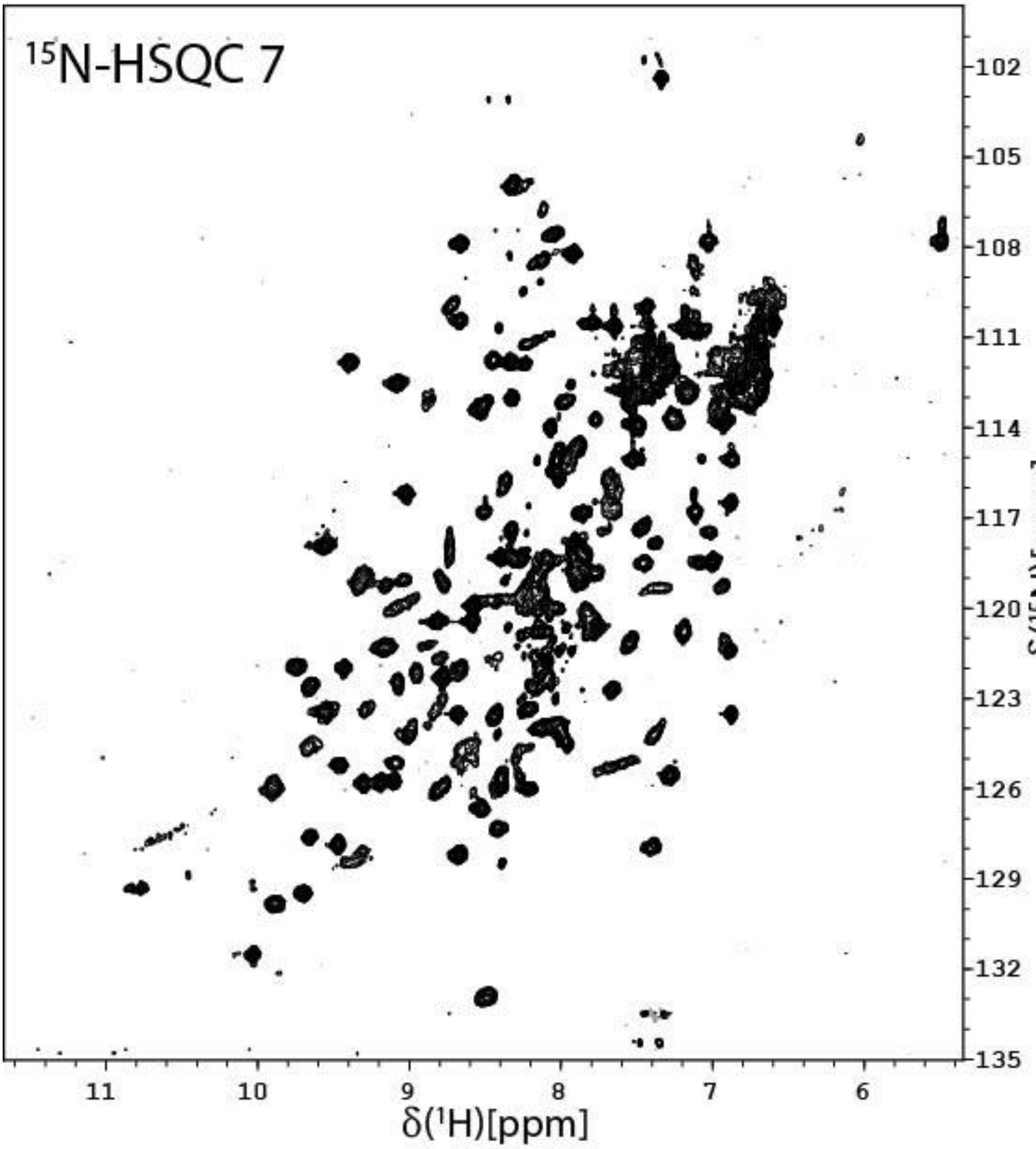


**Figure S8**: The $^{15}N$-HSQC spectrum of *holo-Sm*LPMO10A incubated with 10 mM ascorbic acid for 40h and, subsequently, with 5 mM ascorbic acid and 2 mM $H_2O_2$ for t'= 20h in acetate buffer (25 mM sodium-acetate, 10 mM NaCl, pH 5.0) at 25 °C (treatment series 1; $^{15}N$-HSQC 7).

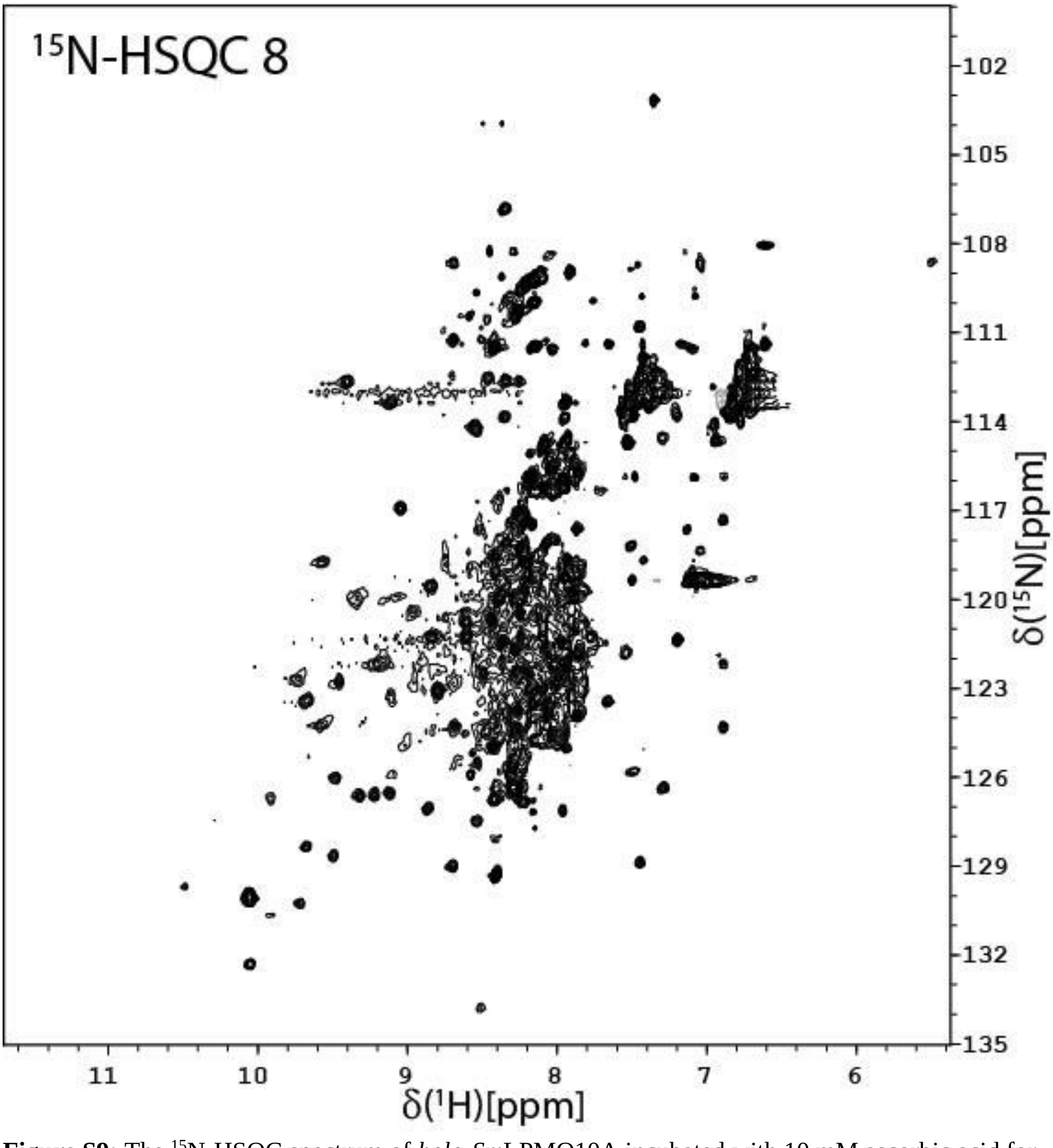


**Figure S9:** The $^{15}N$-HSQC spectrum of *holo*-*Sm*LPMO10A incubated with 10 mM ascorbic acid for 40h and, subsequently, with 5 mM ascorbic acid and 2 mM $H_2O_2$ for t'= 44h in acetate buffer (25 mM sodium-acetate, 10 mM NaCl, pH 5.0) at 25 °C (treatment series 1; $^{15}N$-HSQC 8).

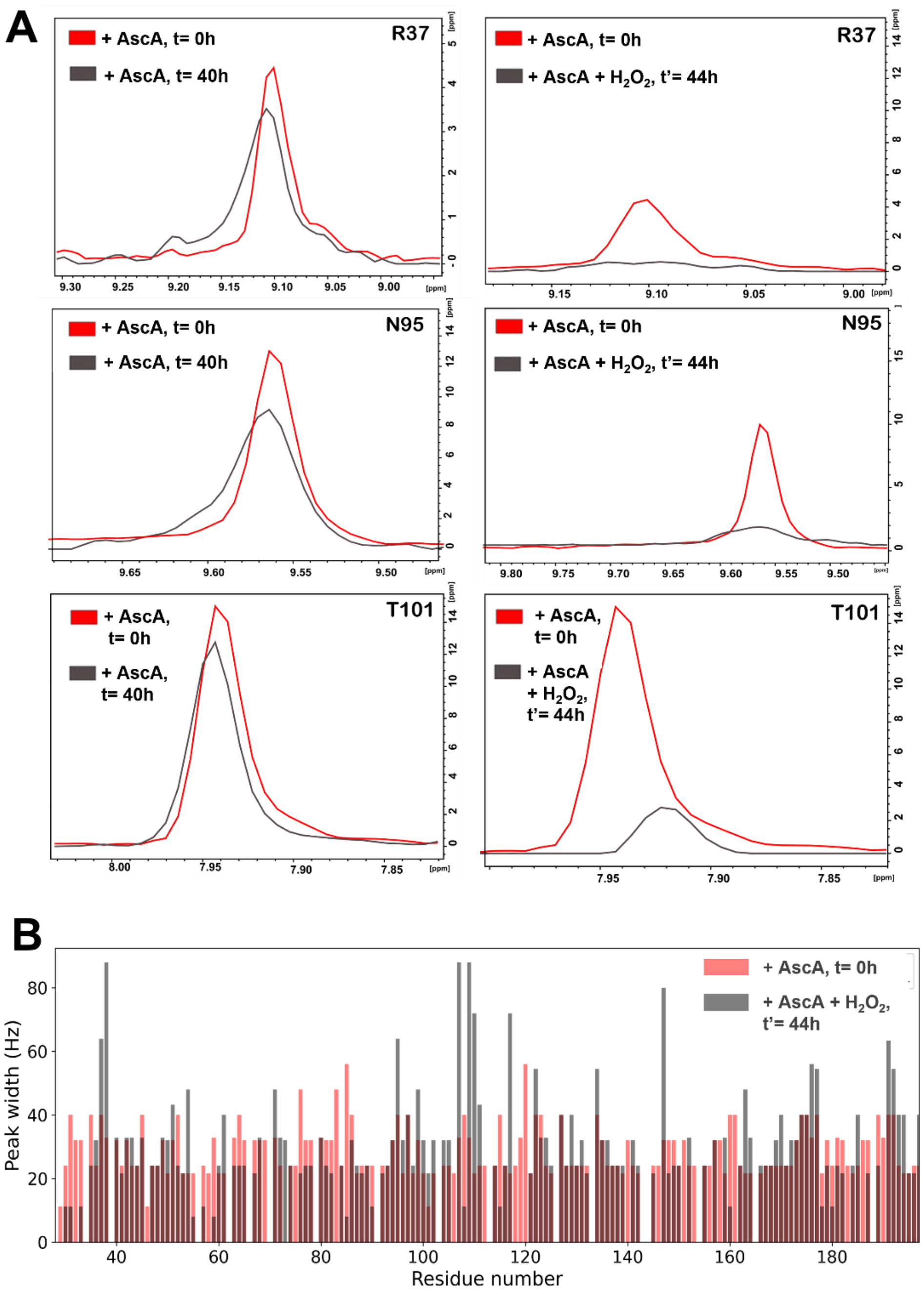


**Figure S10:** Changes in the [1]H resonance linewidths of *Sm*LPMO10A in response to incubation with ascorbic acid and, subsequently, ascorbic acid and $H_2O_2$, in the absence of chitin. **(A)** Changes in the [1]H resonances of residues R37, N95, and T101 in response to incubation with ascorbic acid alone (t= 0h or t= 40h), and, subsequently, ascorbic acid together with $H_2O_2$ (t'= 44h). The t= 0 timepoint is after the addition of ascorbic acid until t= 40min, which is the acquisition time for the $^{15}$N-HSQC spectrum. The linewidths of R37 and N95 increase in response to the treatments, while the linewidth of T101 remains unchanged; addition of $H_2O_2$ next to ascorbic acid speeds up line widening. **(B)** Changes in the [1]H resonance linewidth of all residues of *Sm*LPMO10A in response to incubation with 10 mM ascorbic acid (t= 40h) and, subsequently, with 5 mM ascorbic acid and 2 mM $H_2O_2$ (t'= 44h).

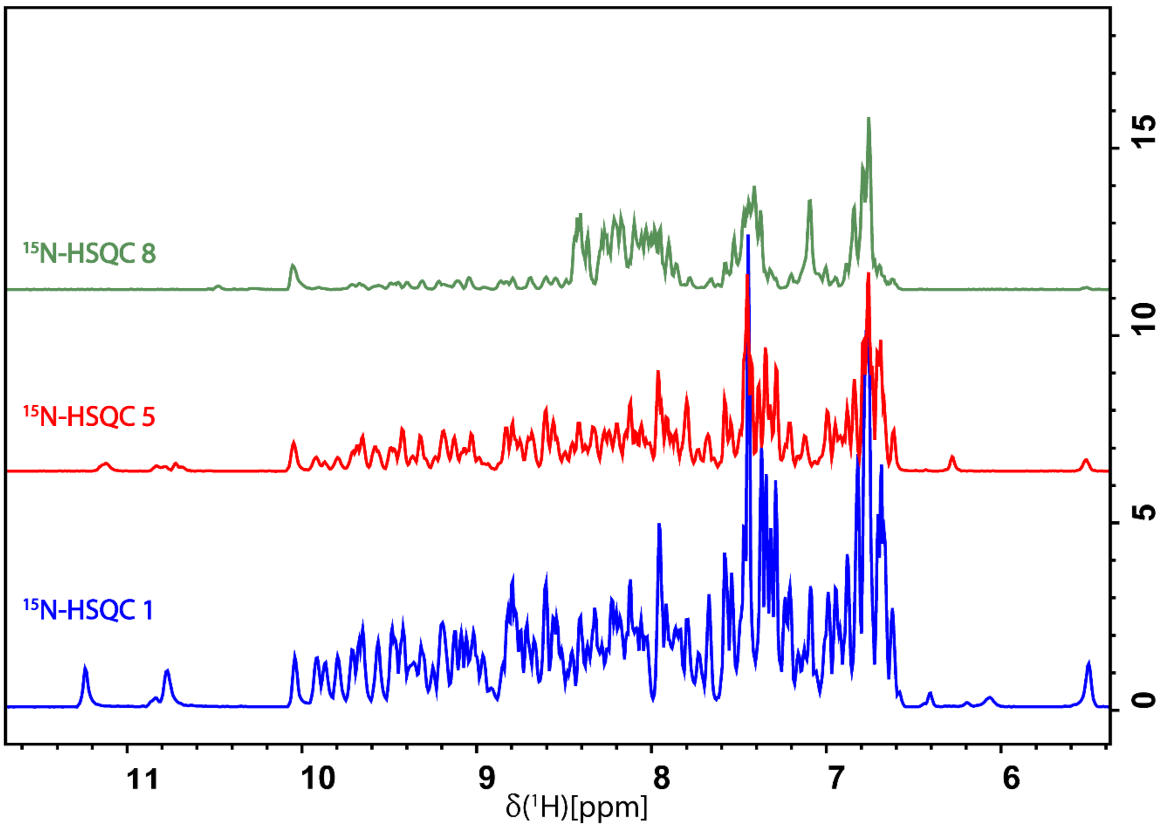


**Figure S11**: $^{1}$H projections from the 2D $^{15}$N-HSQC spectra of *apo*-*Sm*LPMO10A ($^{15}$N-HSQC 1), *holo*-*Sm*LPMO10A incubated with 10 mM ascorbic acid for t= 40h in the absence of the substrate chitin ($^{15}$N-HSQC 5), and *holo*-*Sm*LPMO10A, subsequently incubated with 5 mM ascorbic acid and 2 mM $H_2O_2$ for t'= 44h ($^{15}$N-HSQC 8).

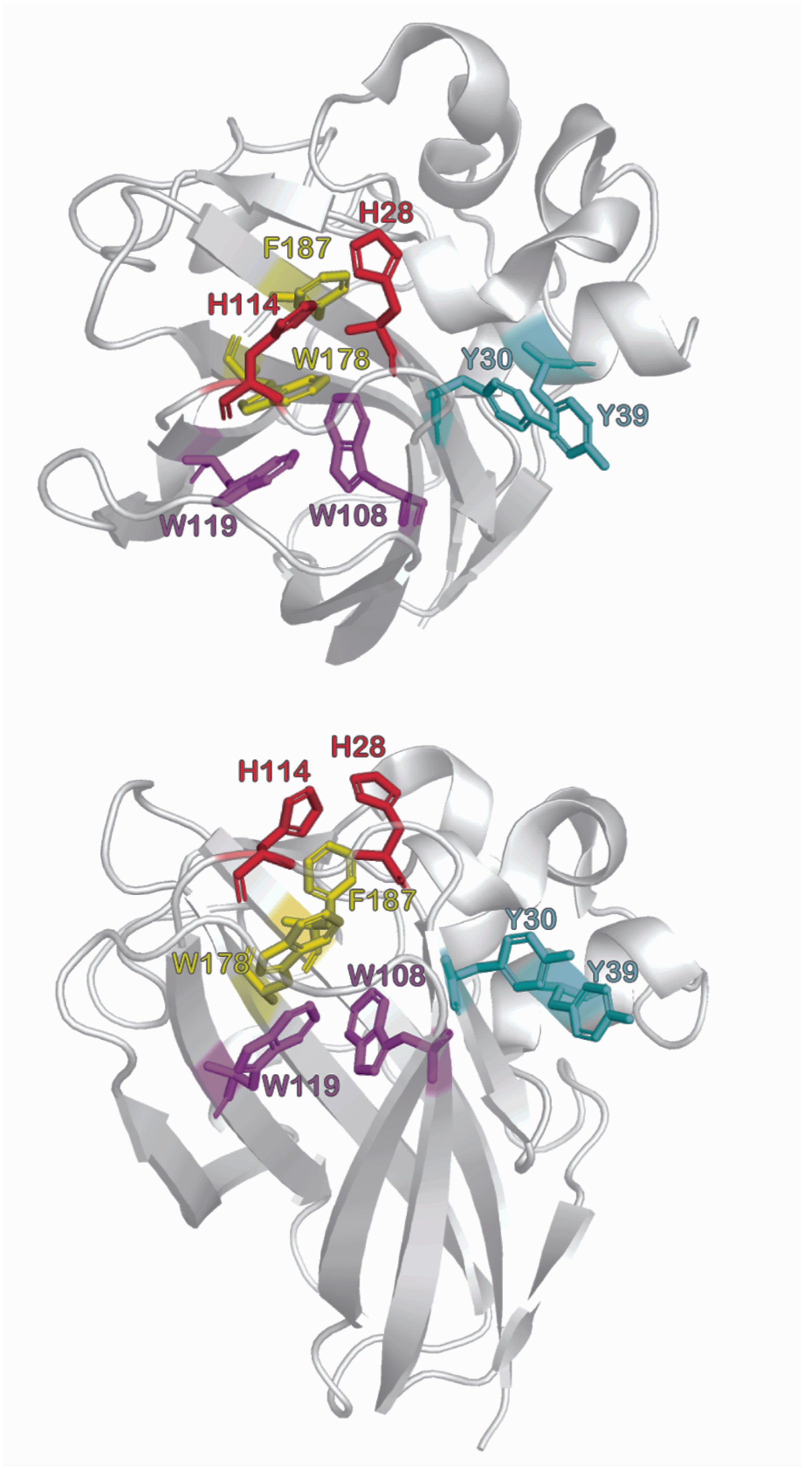


**Figure S12:** Aromatic pairs likely to give rise to π-π* excitation coupling in the CD spectra of SmLPMO10A. The three aromatic pairs Y30-Y39 (cyan), W108-W119 (magenta), and W178-F187 (yellow) are positioned in a way that allows for π-π* excitation coupling. In addition, π-π* excitation coupling is possible between residues W119 and W178 or F187. The aromatic residues are shown on a ribbon diagram of *Sm*LPMO10A (PDB ID: 2BEM).

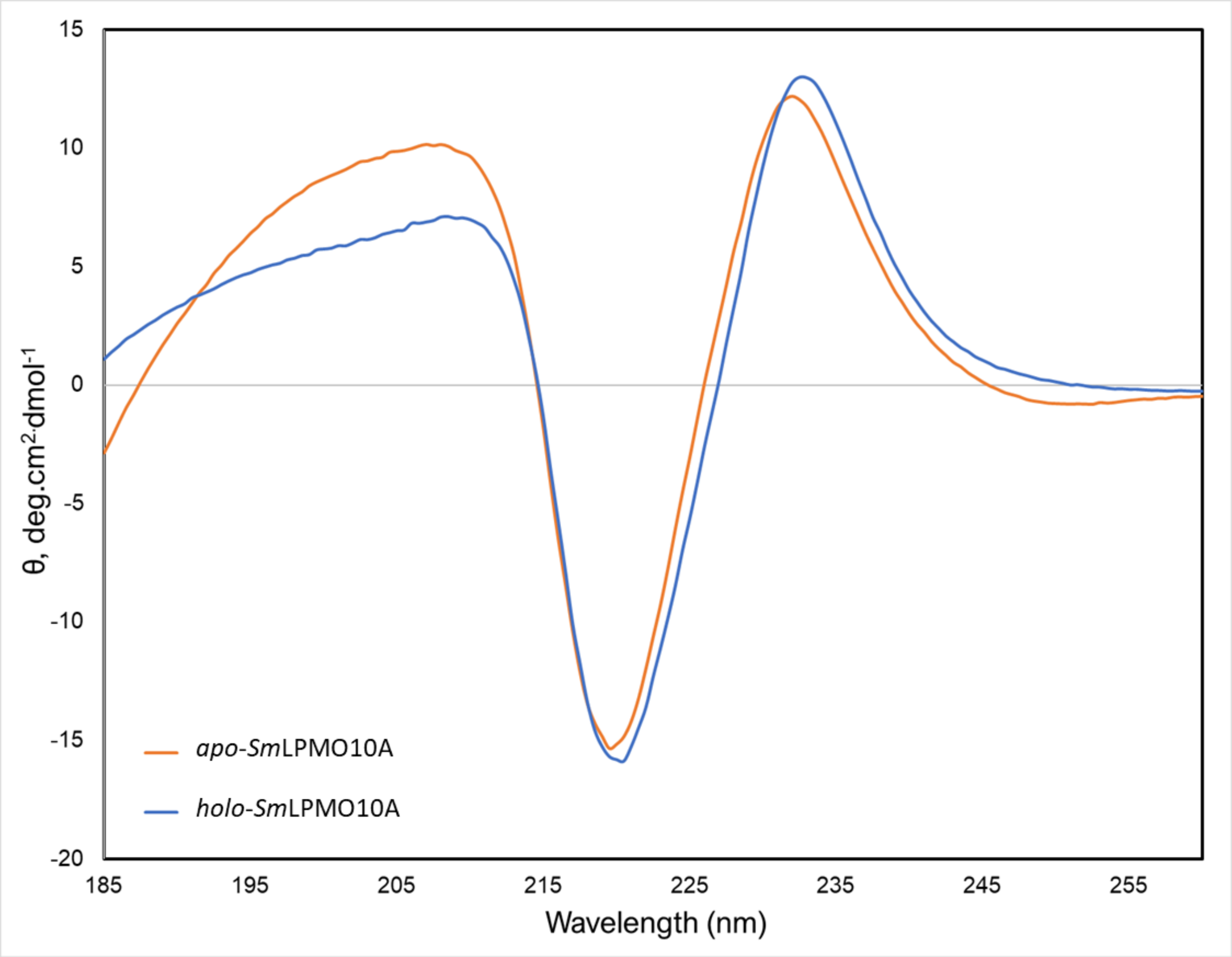


**Figure S13:** Far UV CD spectra of *apo-Sm*LPMO10A and *holo*-SmLPMO10A measured 23 °C. The protein concentration was 2.5 µM in all cases, and the protein was dissolved in acetate buffer (25 mM sodium-acetate, 10 mM NaCl, pH 5.0).

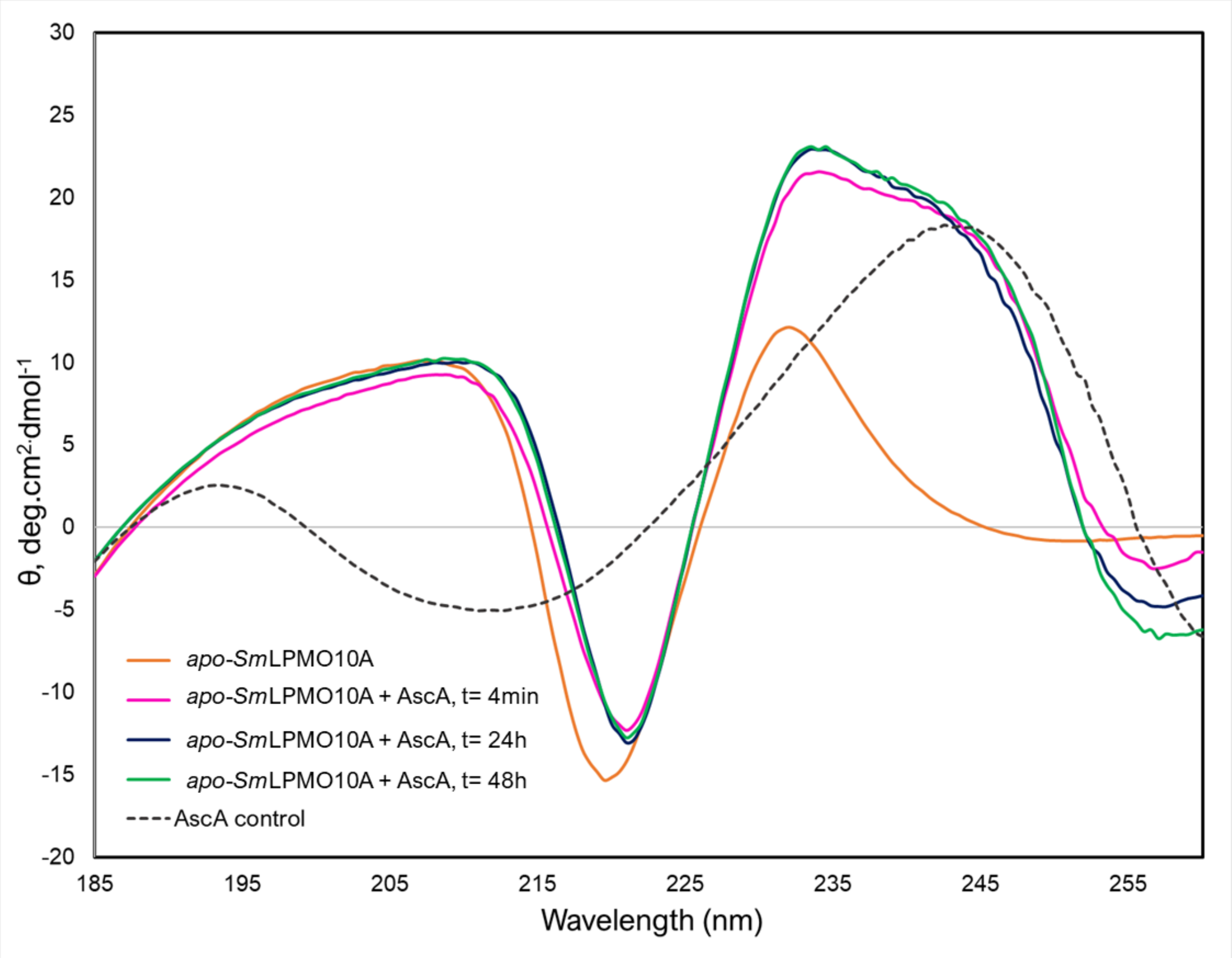


**Figure S14:** Far UV CD spectra of *apo-Sm*LPMO10A and *apo-Sm*LPMO10A incubated with 10 mM ascorbic acid for 4min, 24h, or 48h in the absence of the substrate chitin at 23 °C. The protein concentration was 2.5 µM in all cases and the protein was dissolved in acetate buffer (25 mM sodium-acetate, 10 mM NaCl, pH 5.0).

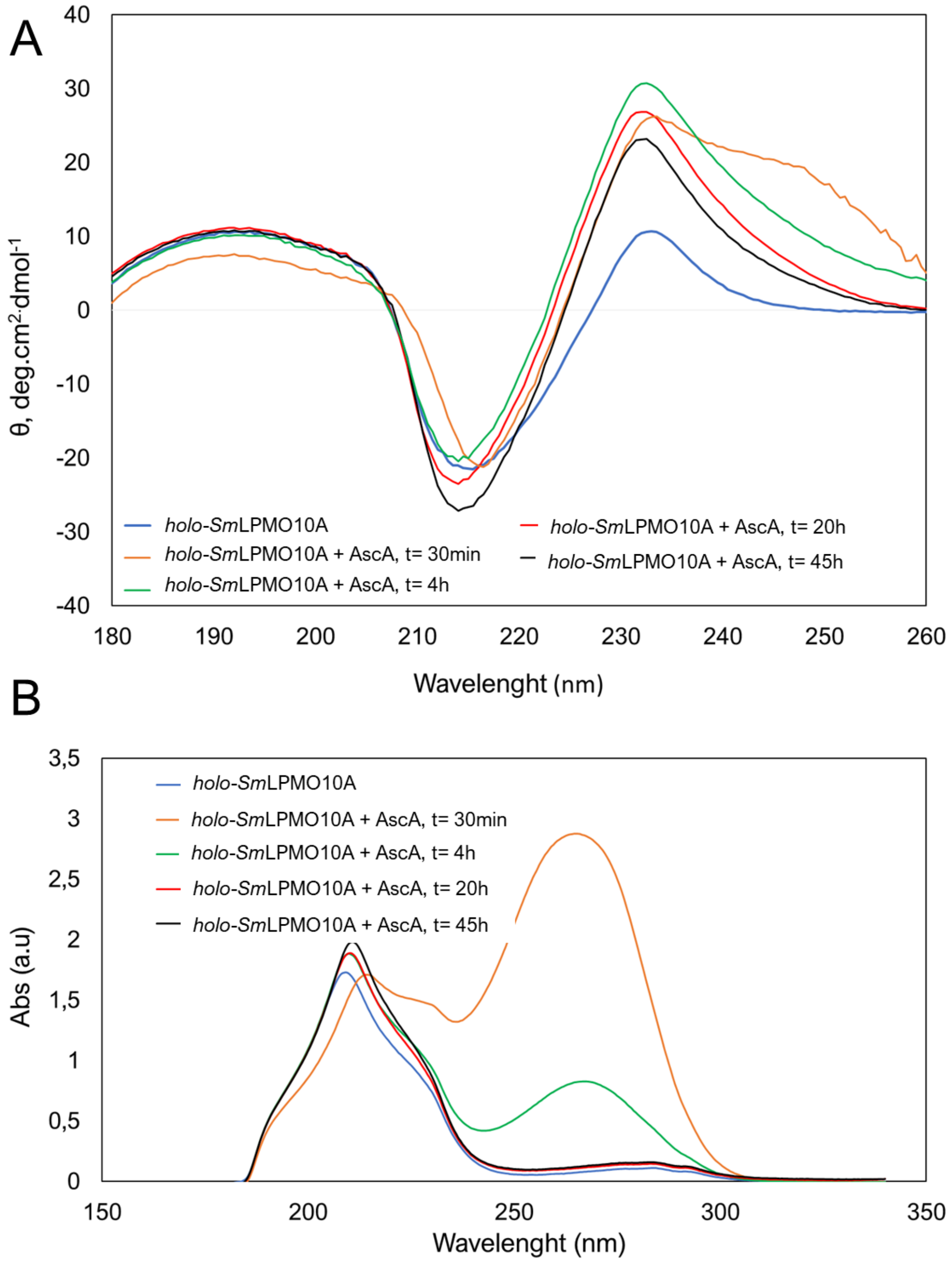


**Figure S15:** Far UV CD (A) and UV absorbance (B) spectra of *holo*-*Sm*LPMO10A and *holo*-SmLPMO10A incubated with 10 mM ascorbic acid for 30min, 4h, 20h, or 45h in the absence of chitin at 23 °C. The protein concentration was 2.5 µM in all cases and the protein was dissolved in acetate buffer (25 mM sodium-acetate, 10 mM NaCl, pH 5.0).

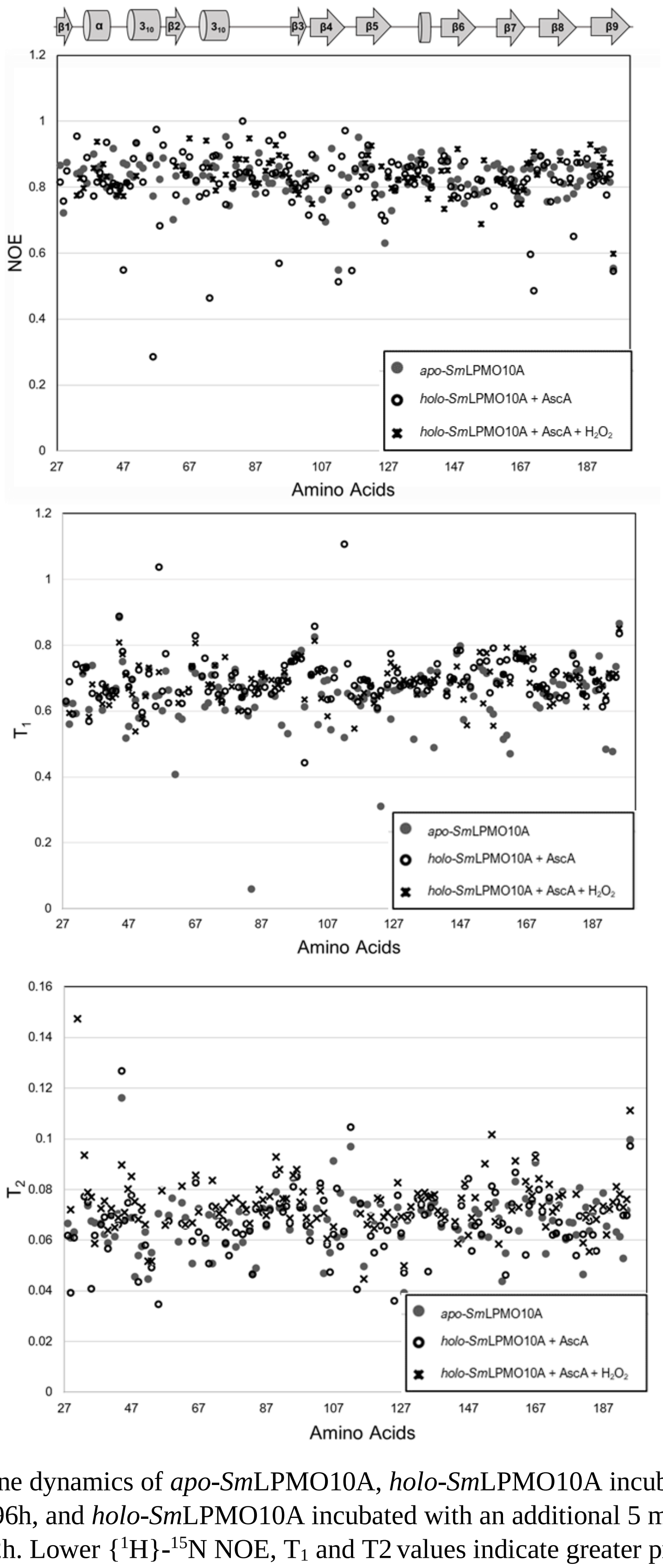


**Figure S16**: Backbone dynamics of *apo*-*Sm*LPMO10A, *holo*-*Sm*LPMO10A incubated with 10 µM ascorbic acid for t= 96h, and *holo*-*Sm*LPMO10A incubated with an additional 5 mM ascorbic acid and 2 mM $H_2O_2$ for t= 72h. Lower {$^1$H}-$^{15}$N NOE, $T_1$ and T2 values indicate greater protein backbone flexibility. However, clear changes in the structure of *Sm*LPMO10A in response to incubation with ascorbic acid and $H_2O_2$ would be seen in the same areas of the structure in all three experiments. No such conformational changes were clear in the displayed results which were ambiguous.

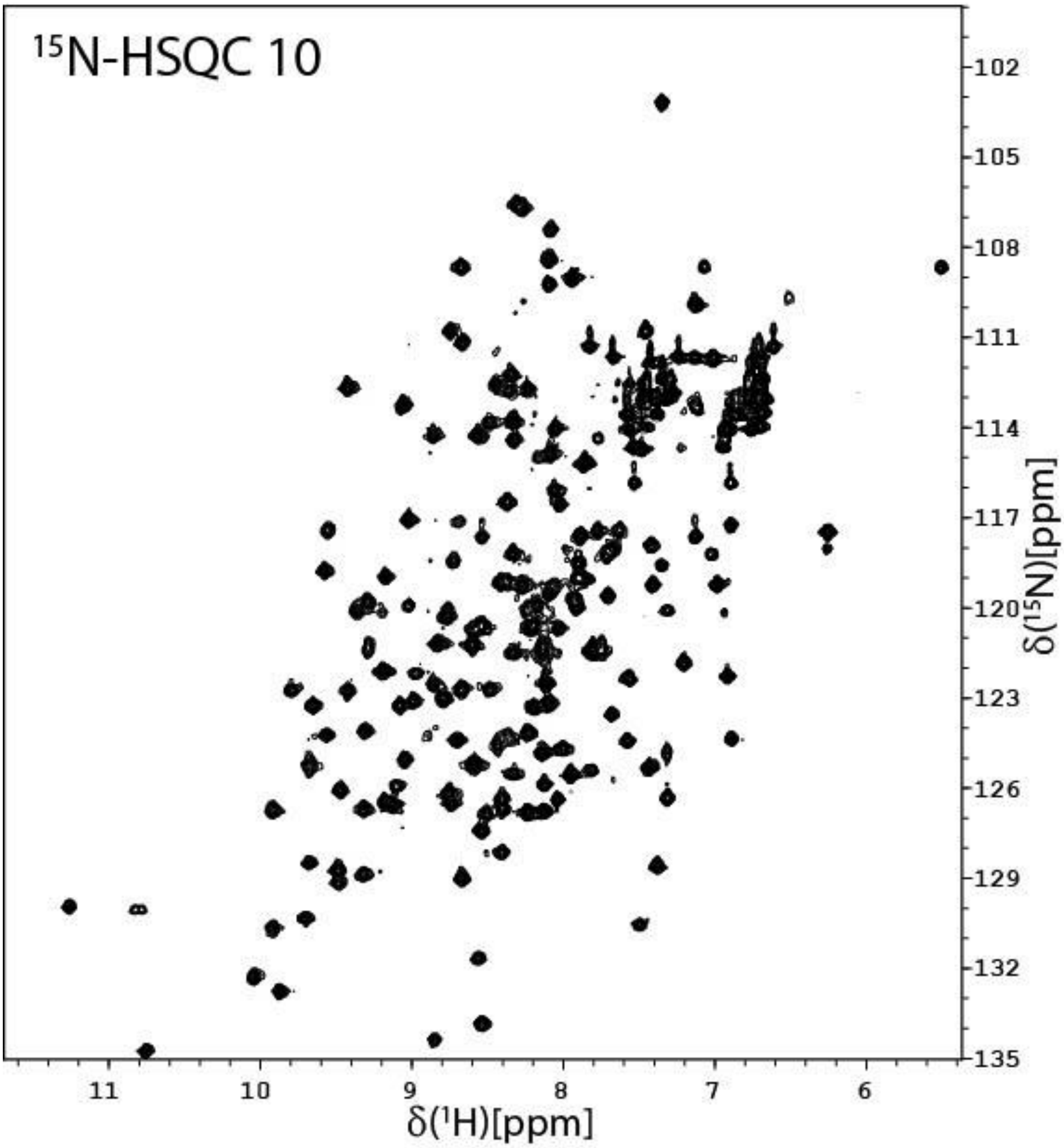


**Figure S17:** The $^{15}$N-HSQC spectrum of *holo*-*Sm*LPMO10A incubated with 10 mM ascorbic acid for t= 0h in acetate buffer (25 mM sodium-acetate, 10 mM NaCl, pH 5.0) in the presence of 10 mg β-chitin particles with a size of ~ 0.5 mm (1 h preincubation with chitin prior to addition of ascorbic acid) at 25 °C (treatment series 2; $^{15}$N-HSQC 10).

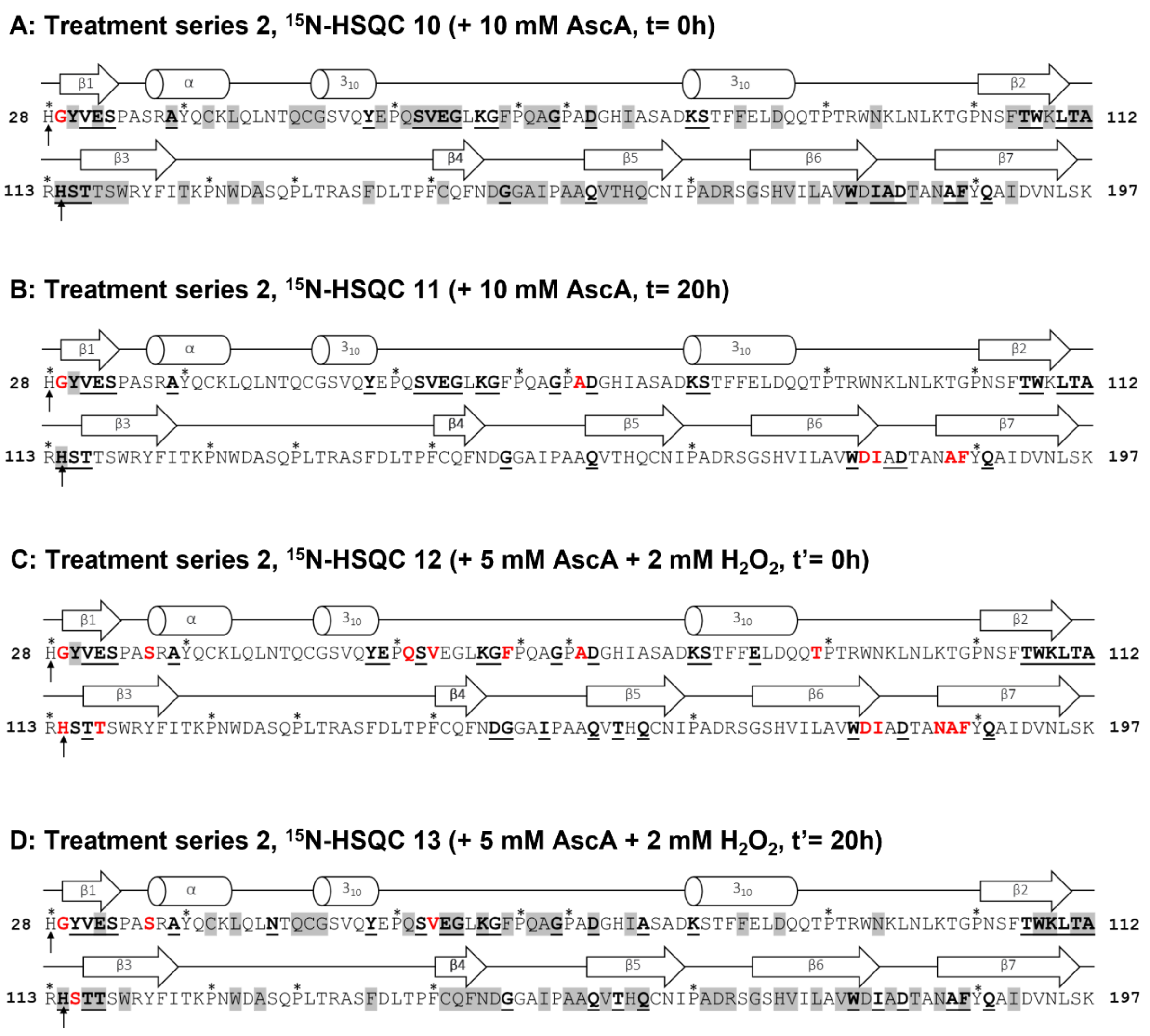


**Figure S18**: The combined structural changes to *holo-Sm*LPMO10A observed in treatment series 2 in the presence of the substrate β-chitin (1h pre-incubation), relative to the reference spectrum of *apo-Sm*LPMO10A. The active site histedines are indicated by arrows, while residues not assigned in by NMR are indicated by asterixis. Chemical shift perturbations >50Hz are shown as by bold and underlined residues, narrower linewidths (i.e increased signal intensities) are shown by residues highlighted in grey. Non-detectable $^{1}H$-$^{15}N$ signals are shown by red residues.

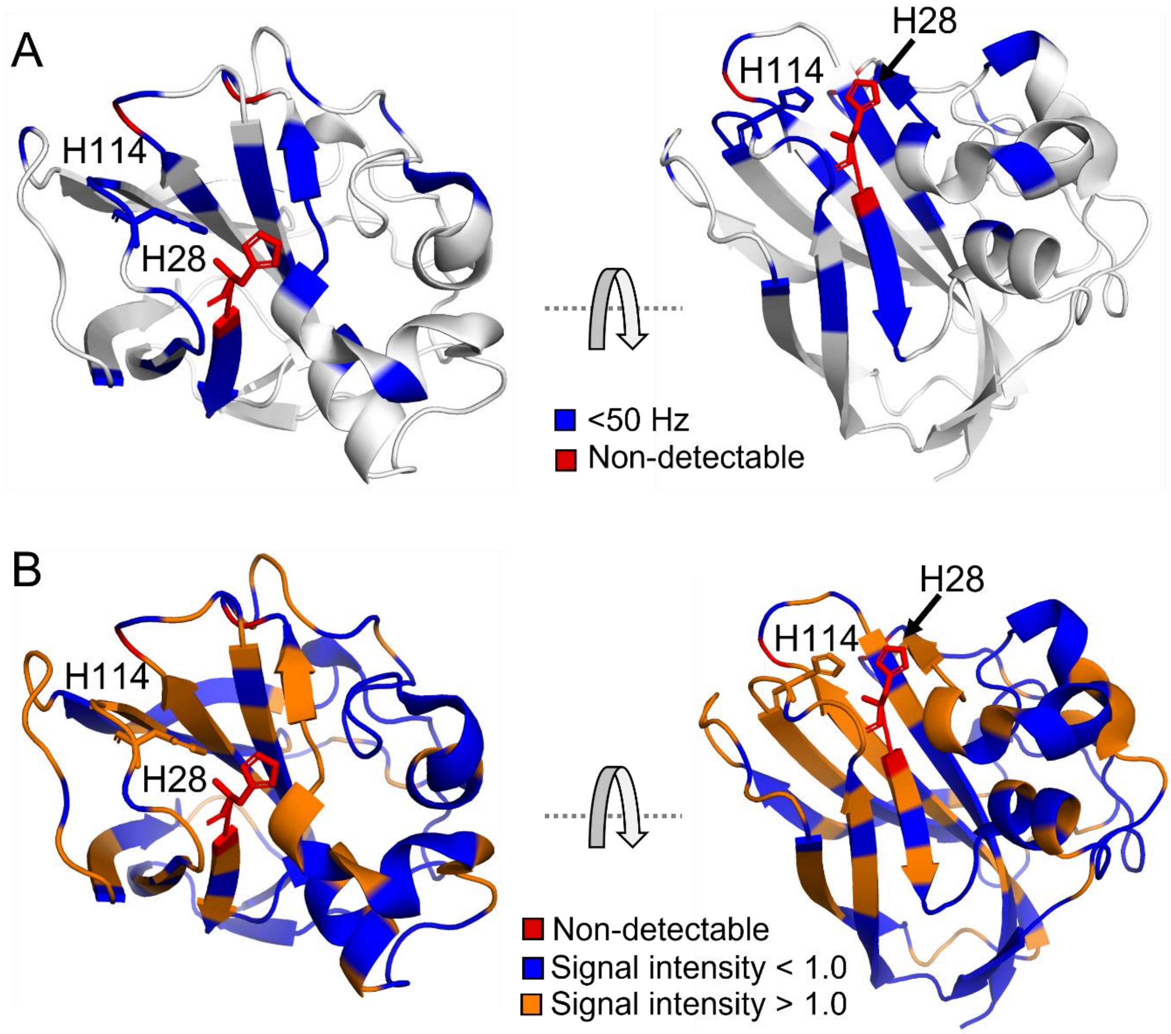


**Figure S19**: Immediate (t= 0h) effect of incubating *holo*-*Sm*LPMO10A with 10 mM ascorbic acid under aerobic conditions in the presence of β-chitin particles (treatment series 2). **A)** Chemical Shift Perturbations (CSPs) in response to addition of ascorbic acid (at t= 0h) and copper in the presence of chitin compared to *apo*-*Sm*LPMO10A highlighted on the X-ray crystal structure of *Sm*LPMO10A (PDB ID: 2BEM). The magnitude of the change in Hz is shown by the color scheme. Non-detectable residues are shown in red. **B)** Narrower linewidths in response to addition of ascorbic acid (at t= 0h) and copper reduction in the presence of chitin compared with *apo*-*Sm*LPMO10A highlighted on the X-ray crystal structure of *Sm*LPMO10A. An increase in signal intensity is shown in orange, while a decrease is shown in blue. Non-detectable residues are shown in red.

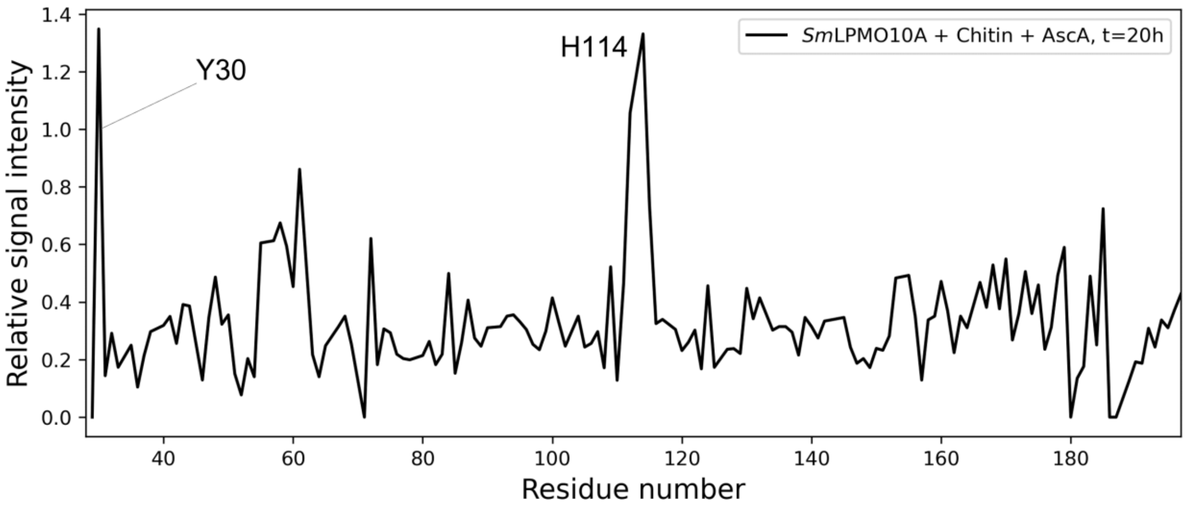


**Figure S20:** The relative intensities of $^{1}H$-$^{15}N$ signals in the $^{15}N$-HSQC spectra of *holo*-*Sm*LPMO10A incubated with 10 mM ascorbic acid for t= 20h in the presence of chitin ($^{15}N$-HSQC 11) and the intensities of $^{1}H$-$^{15}N$ signal intensities in the $^{15}N$-HSQC spectrum of *apo*-*Sm*LPMO10A in the absence of chitin ($^{15}N$-HSQC 1). Only Y30 and H114 show an increase in signal intensity.

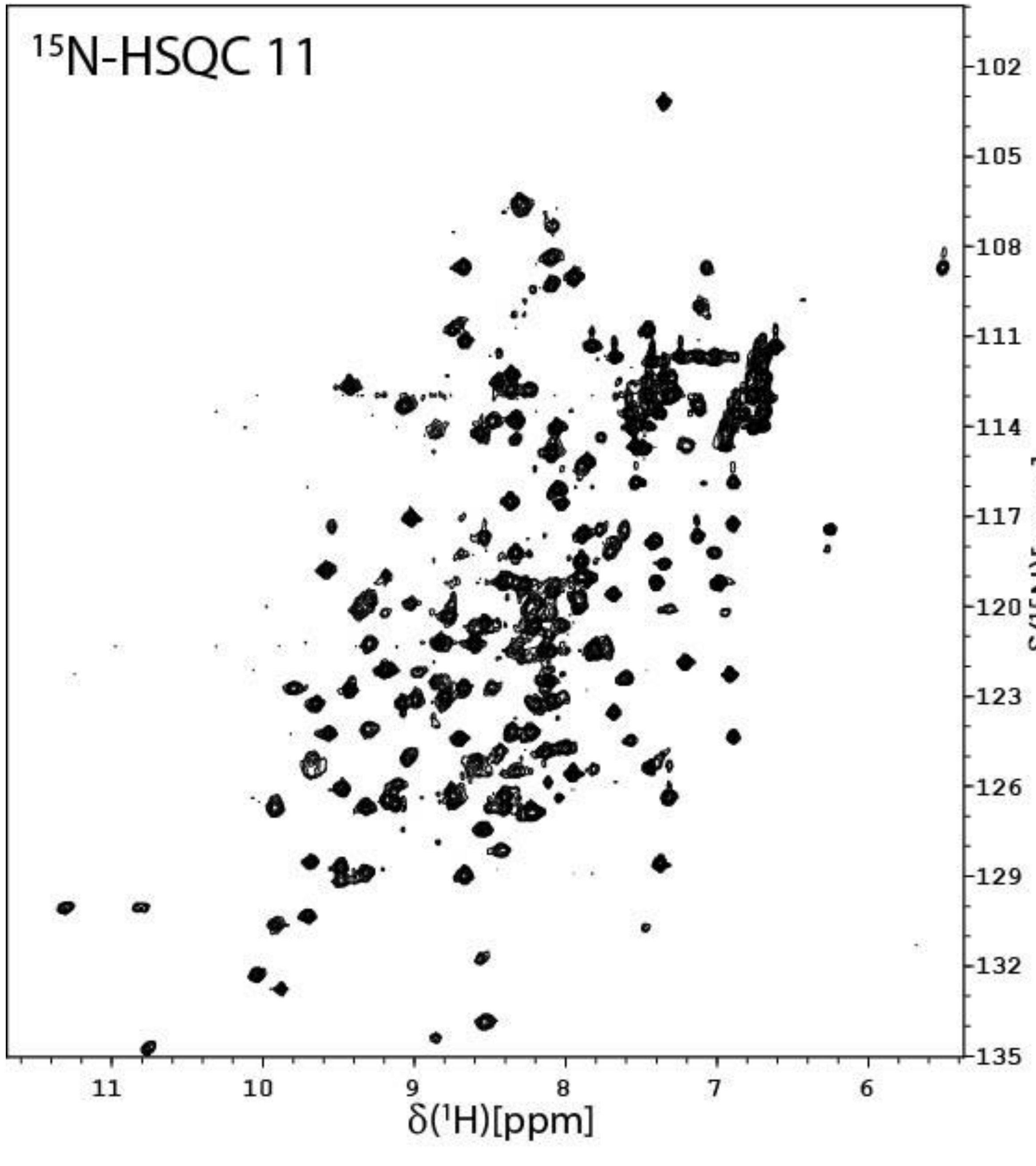


**Figure S21:** The $^{15}$N-HSQC of *holo*-*Sm*LPMO10A incubated with 10 mM ascorbic acid for t= 20h in acetate buffer (25 mM sodium-acetate, 10 mM NaCl, pH 5.0) in the presence of 10 mg β-chitin particles with a size of ~ 0.5 mm (1h preincubation) at 25 °C (treatment series 2, $^{15}$N-HSQC 11).

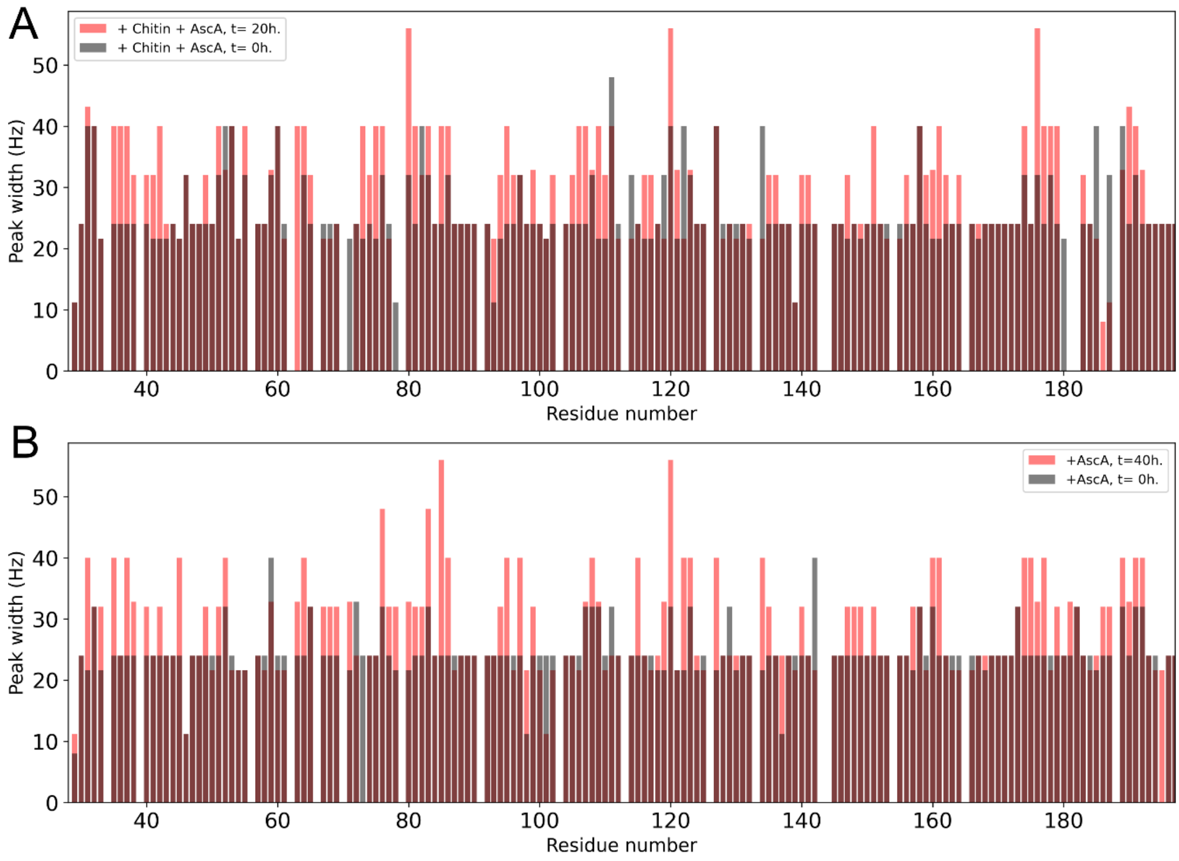


**Figure S22: A)** Changes in the $^{1}$H resonance linewidth of all residues of *holo*-*Sm*LPMO10A in response to t= 0h (treatment series 2, $^{15}$N-HSQC 10) and t= 20h (treatment series 2, $^{15}$N-HSQC 11) incubation periods with ascorbic acid in the presence of chitin (1h pre-incubation with chitin alone). **B)** Changes in the $^{1}$H resonance linewidth of all residues of *holo*-*Sm*LPMO10A in response to t= 0h (treatment series 1, $^{15}$N-HSQC 3) and t= 40h (treatment series 1, $^{15}$N-HSQC 5) incubation periods with ascorbic acid.

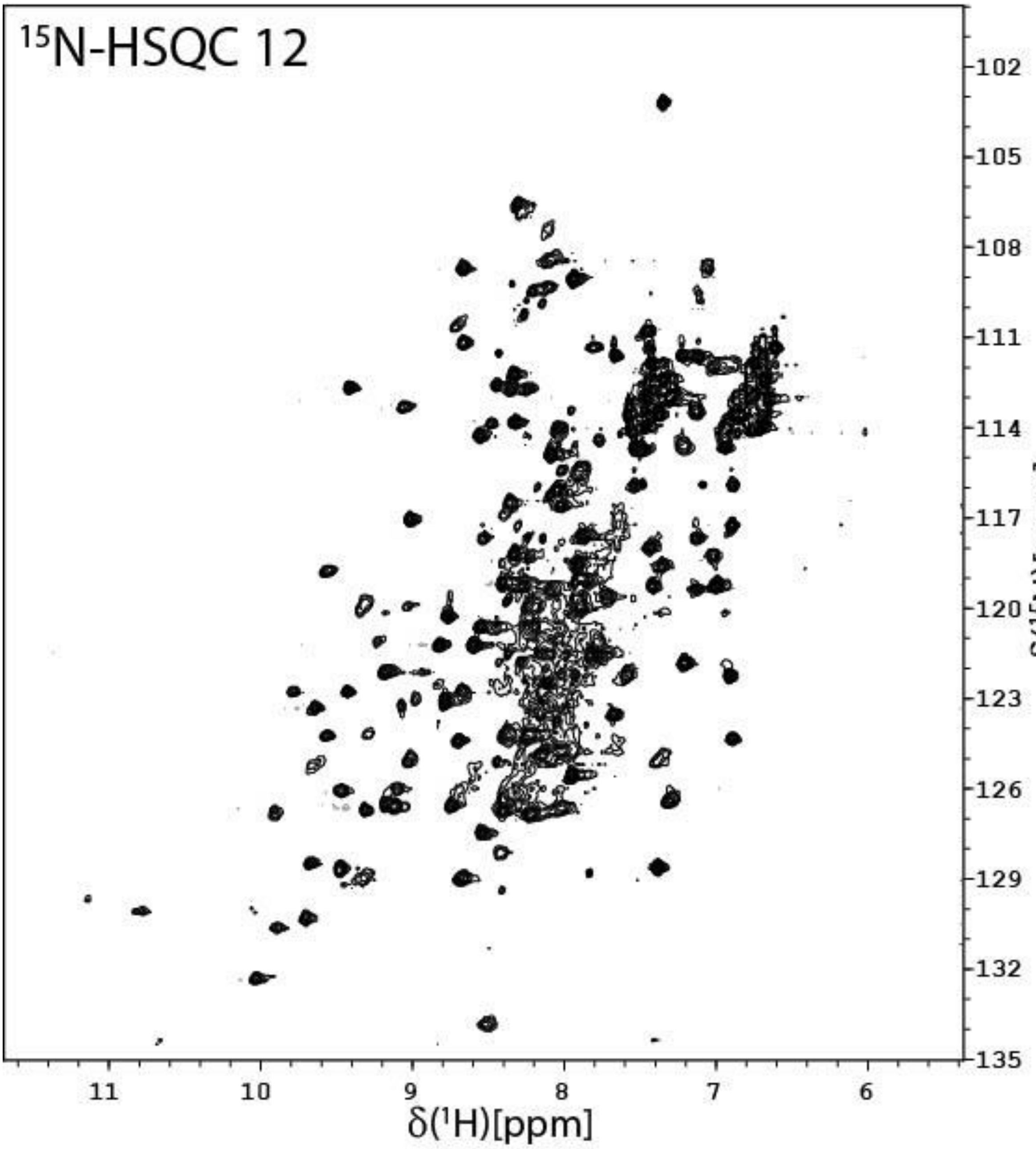


**Figure S23:** The $^{15}$N-HSQC of *holo*-*Sm*LPMO10A incubated with 10 mM ascorbic acid for 20h and, subsequently, with 5 mM ascorbic acid and 2 mM $H_2O_2$ for t'= 0h in acetate buffer (25 mM sodium-acetate, 10 mM NaCl, pH 5.0) in the presence of 10 mg β-chitin particles with a size of ~ 0.5 mm (1h preincubation) at 25 °C (treatment series 2, $^{15}$N-HSQC 12).

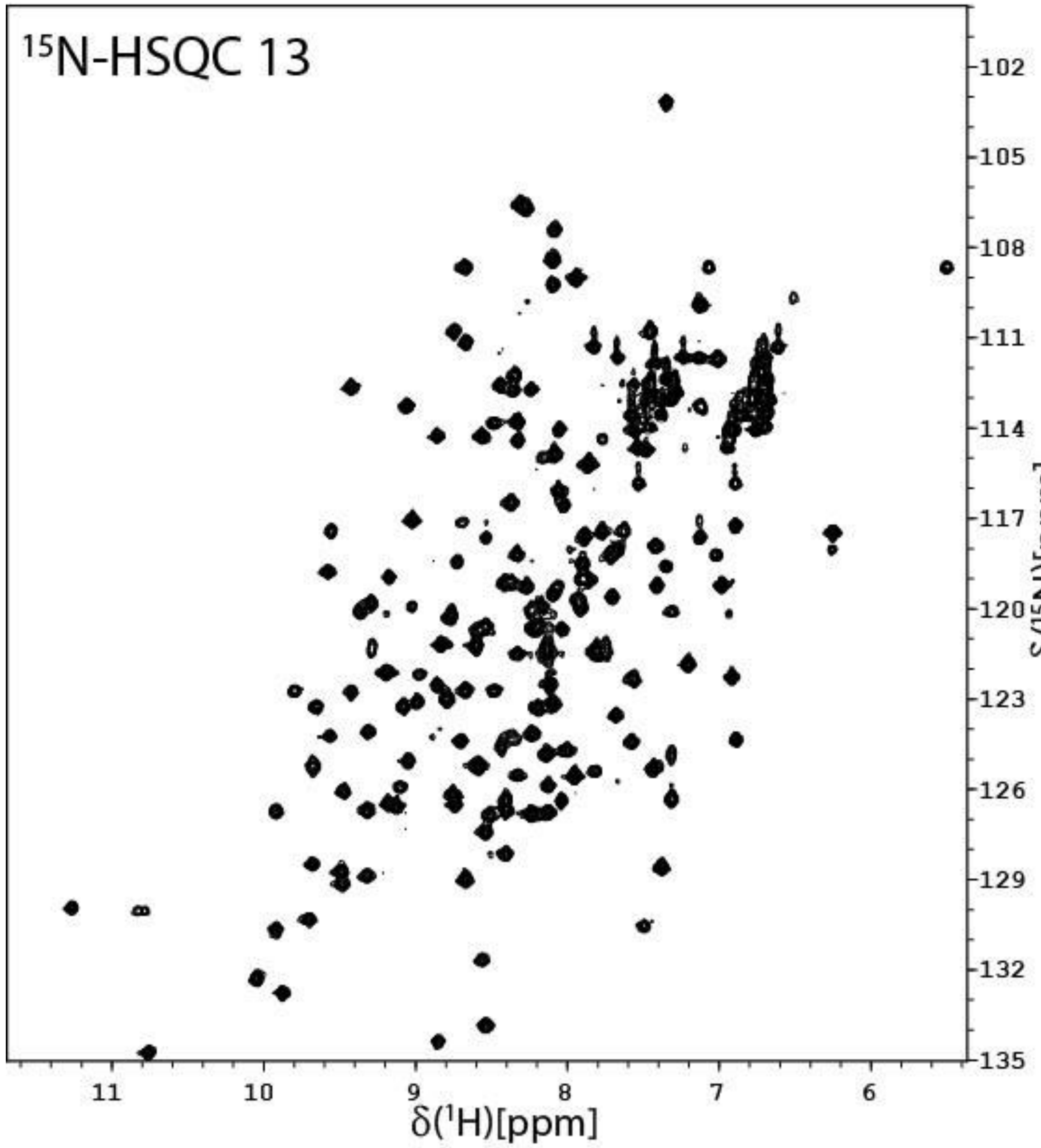


**Figure S24:** The $^{15}N$-HSQC of *holo*-*Sm*LPMO10A incubated with 10 mM ascorbic acid for 20h and, subsequently, with 5 mM ascorbic acid and 2 mM $H_2O_2$ for t'= 20h in acetate buffer (25 mM sodium-acetate, 10 mM NaCl, pH 5.0) in the presence of 10 mg β-chitin particles with a size of ~ 0.5 mm (1h preincubation) at 25 °C (treatment series 2, $^{15}N$-HSQC 13).

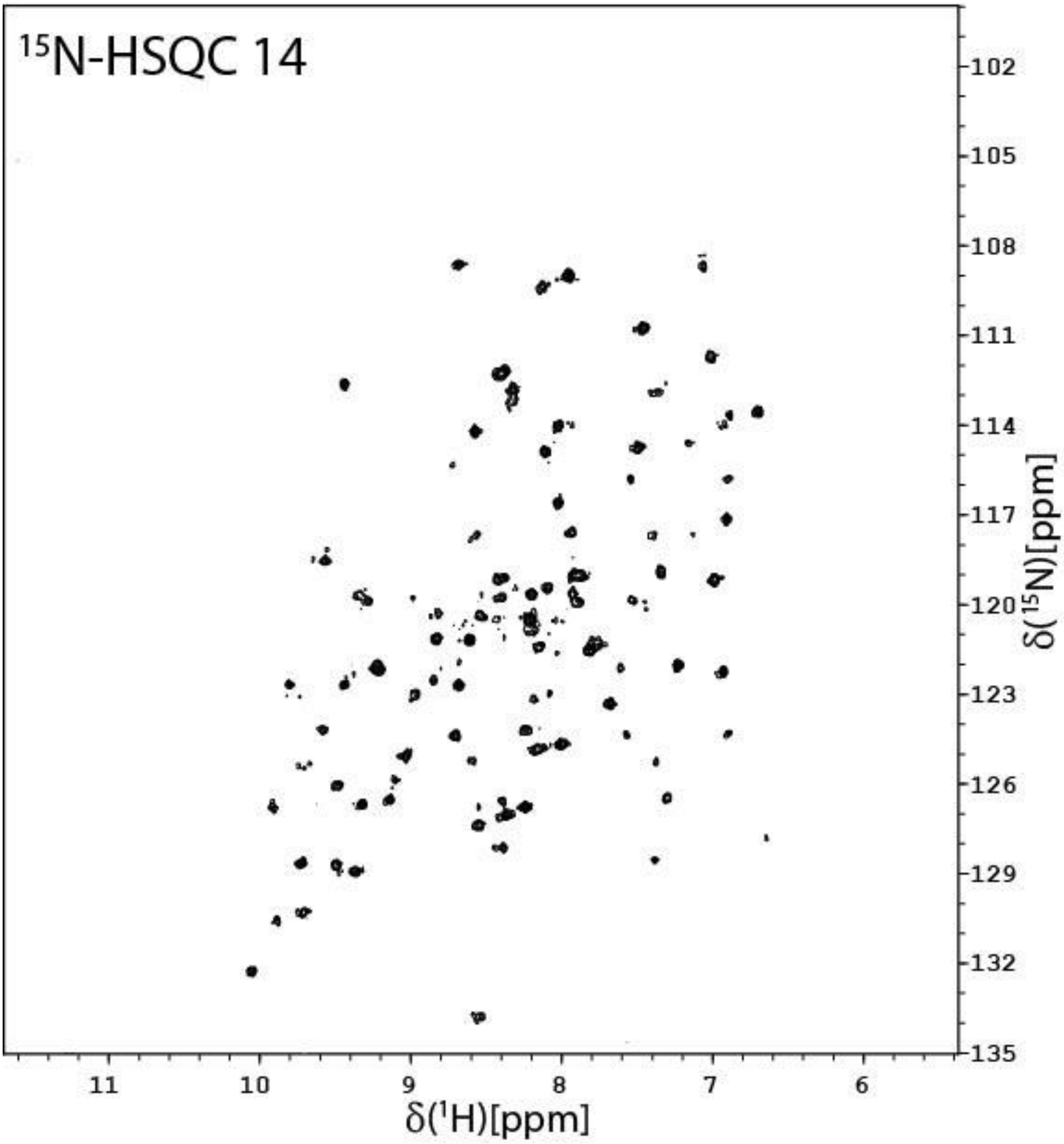


**Figure S25:** The $^{15}$N-HSQC of *holo*-*Sm*LPMO10A in acetate buffer (25 mM sodium-acetate, 10 mM NaCl, pH 5.0) in the presence of 10 mg β-chitin particles with a size of ~ 0.5 mm (24h preincubation) at 25 °C (treatment series 3, 15N-HSQC 14).

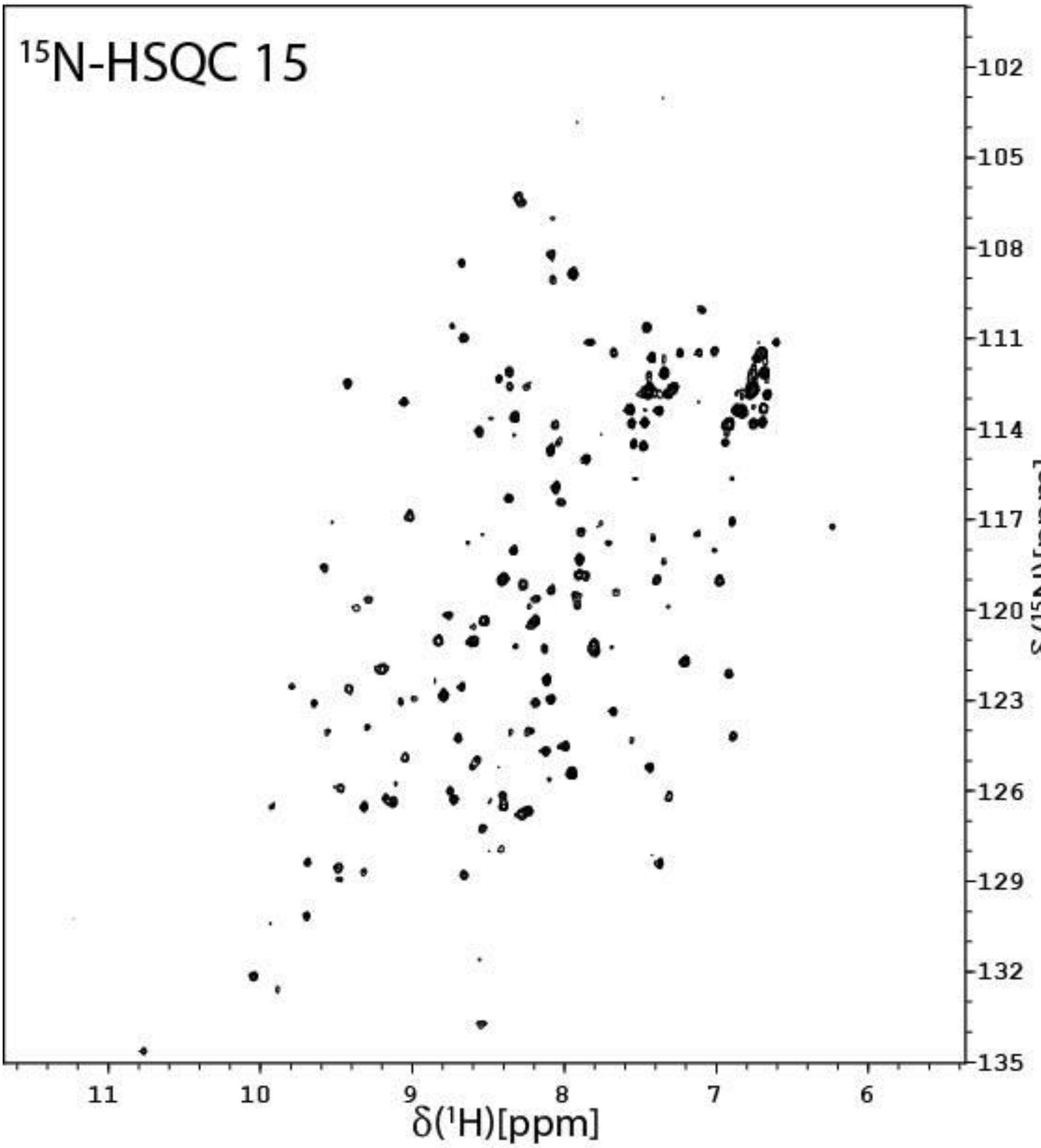


**Figure S26**: The $^{15}$N-HSQC of *holo*-*Sm*LPMO10A incubated with 10 mM ascorbic acid for t= 0h in acetate buffer (25 mM sodium-acetate, 10 mM NaCl, pH 5.0) in the presence of 10 mg β-chitin particles with a size of ~ 0.5 mm (24h preincubation) at 25 °C (treatment series 3, $^{15}$N-HSQC 15).

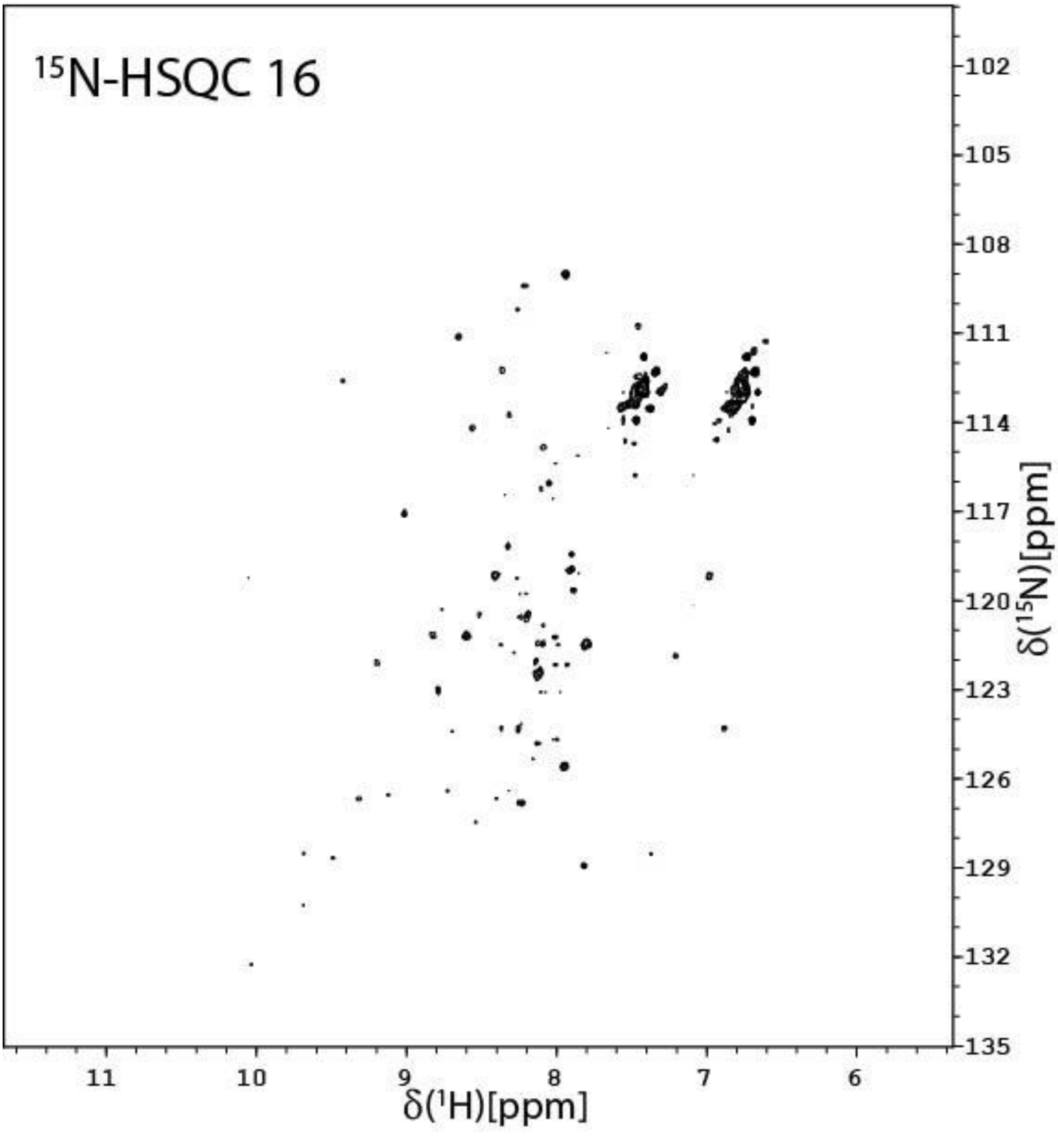


**Figure S27:** The $^{15}$N-HSQC of *holo*-*Sm*LPMO10A incubated with 10 mM ascorbic acid for t= 24h in acetate buffer (25 mM sodium-acetate, 10 mM NaCl, pH 5.0) in the presence of 10 mg β-chitin particles with a size of ~ 0.5 mm (24h preincubation) at 25 °C (treatment series 3, $^{15}$N-HSQC 16).

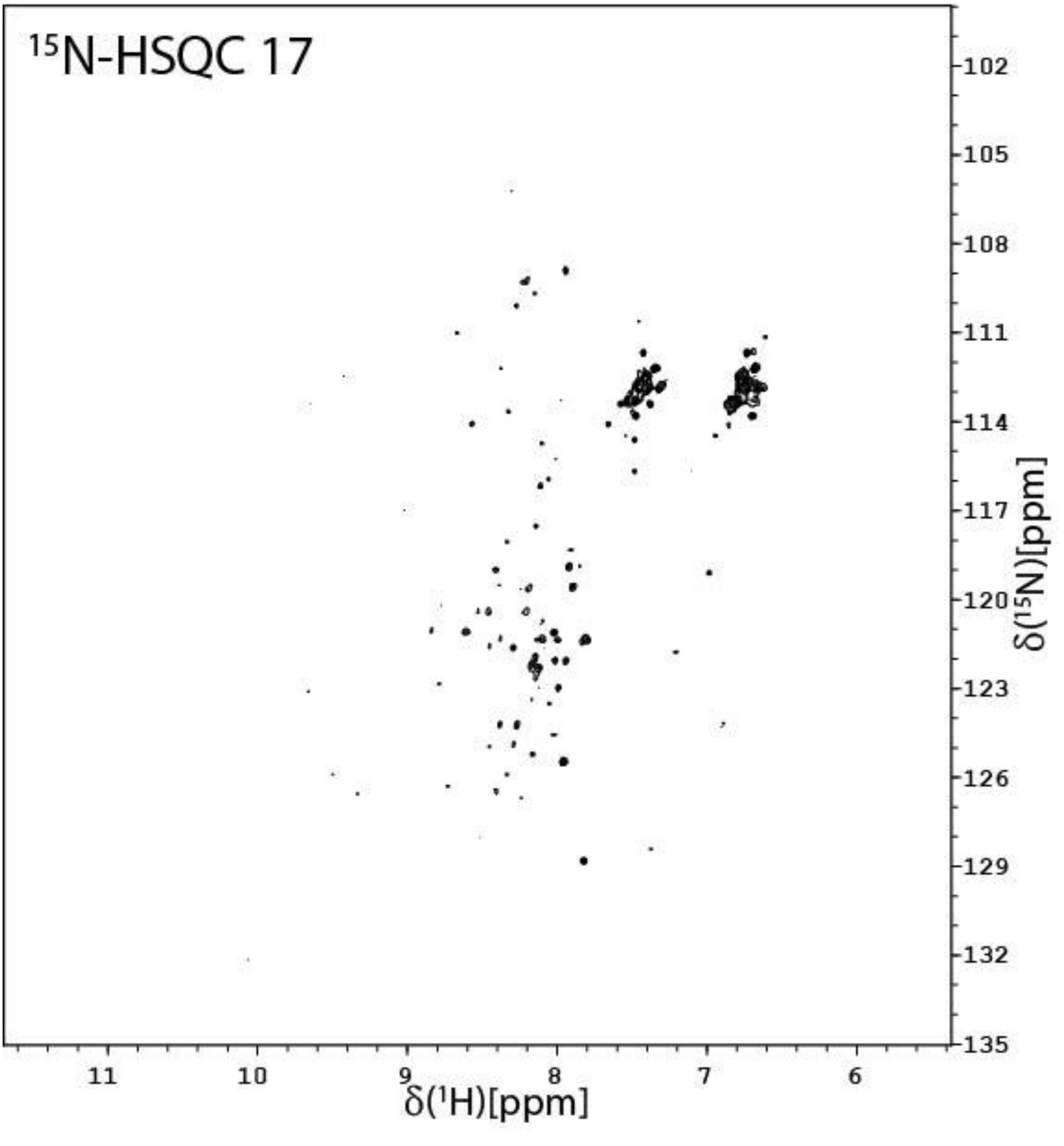


**Figure S28:** The $^{15}$N-HSQC of *holo-Sm*LPMO10A incubated with 10 mM ascorbic acid for t= 48h in acetate buffer (25 mM sodium-acetate, 10 mM NaCl, pH 5.0) in the presence of 10 mg β-chitin particles with a size of ~ 0.5 mm (24h preincubation) at 25 °C (treatment series 3, $^{15}$N-HSQC 17).

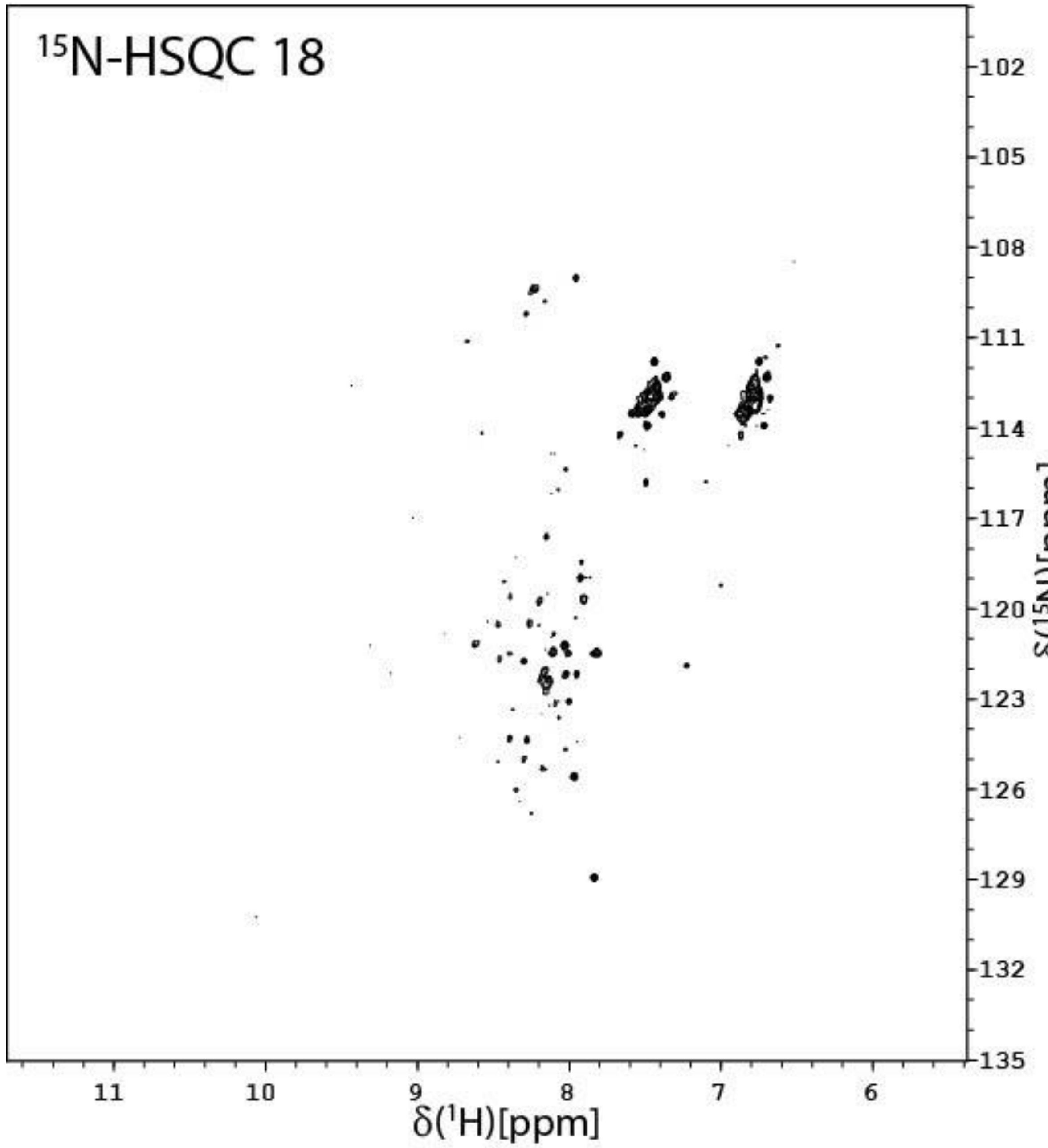


**Figure S29:** The $^{15}$N-HSQC of *holo-Sm*LPMO10A incubated with 10 mM ascorbic acid for t= 78h in acetate buffer (25 mM sodium-acetate, 10 mM NaCl, pH 5.0) in the presence of 10 mg β-chitin particles with a size of ~ 0.5 mm (24h preincubation) at 25 °C (treatment series 3, $^{15}$N-HSQC 18).

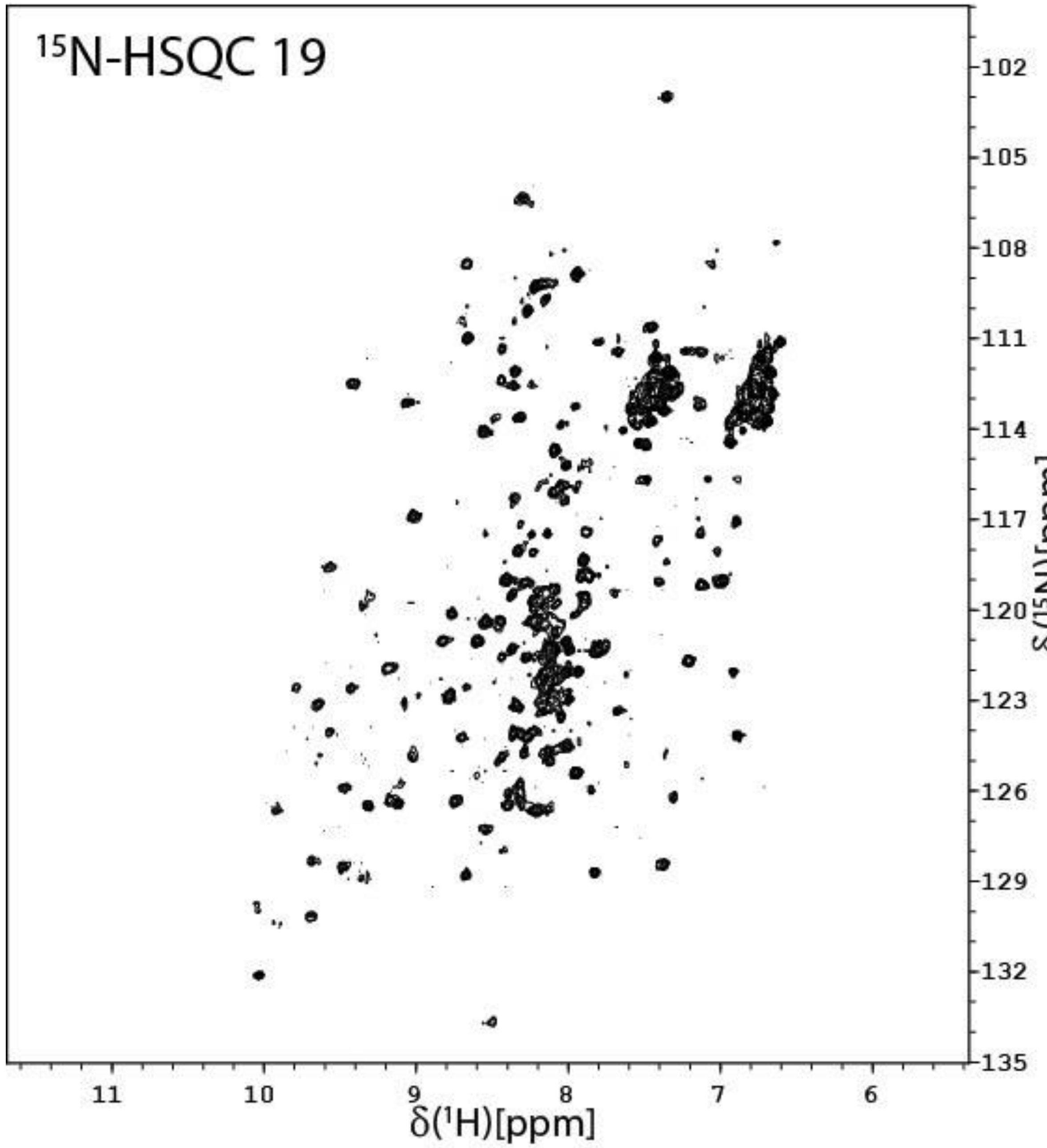


**Figure S30:** The $^{15}$N-HSQC of *holo*-*Sm*LPMO10A incubated with 10 mM ascorbic acid for 78h and then 5 mM ascorbic acid and 2 mM $H_2O_2$ for t'= 0h in acetate buffer (25 mM sodium-acetate, 10 mM NaCl, pH 5.0) in the presence of 10 mg β-chitin particles with a size of ~ 0.5 mm (24h preincubation) at 25 °C (treatment series 3, $^{15}$N-HSQC 19).

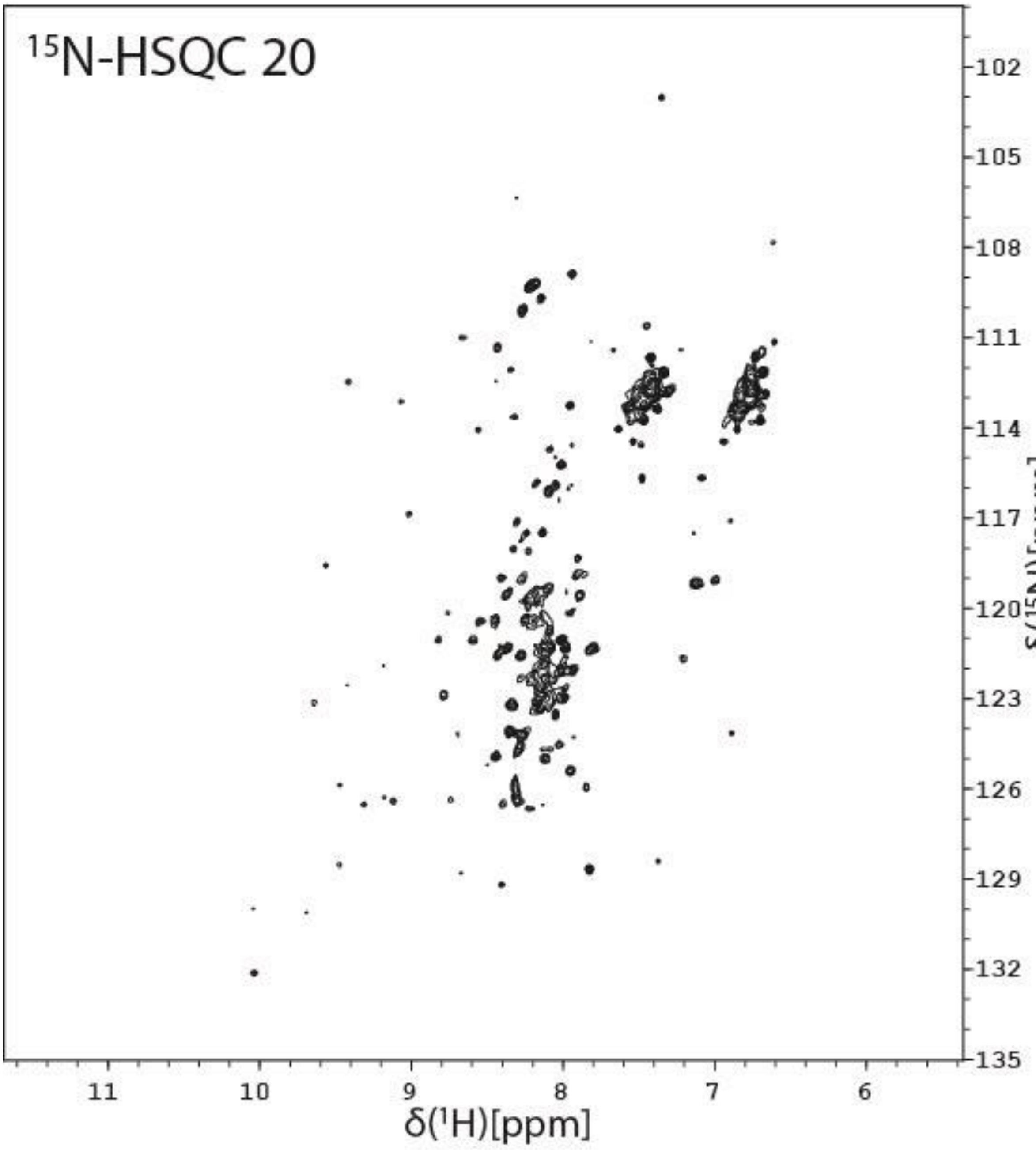


**Figure S31:** The $^{15}N$-HSQC of *holo*-*Sm*LPMO10A incubated with 10 mM ascorbic acid for 78h and then 5 mM ascorbic acid and 2 mM $H_2O_2$ for t'= 24h in acetate buffer (25 mM sodium-acetate, 10 mM NaCl, pH 5.0) in the presence of 10 mg β-chitin particles with a size of ~ 0.5 mm (24h preincubation) at 25 °C (treatment series 3, $^{15}N$-HSQC 20).

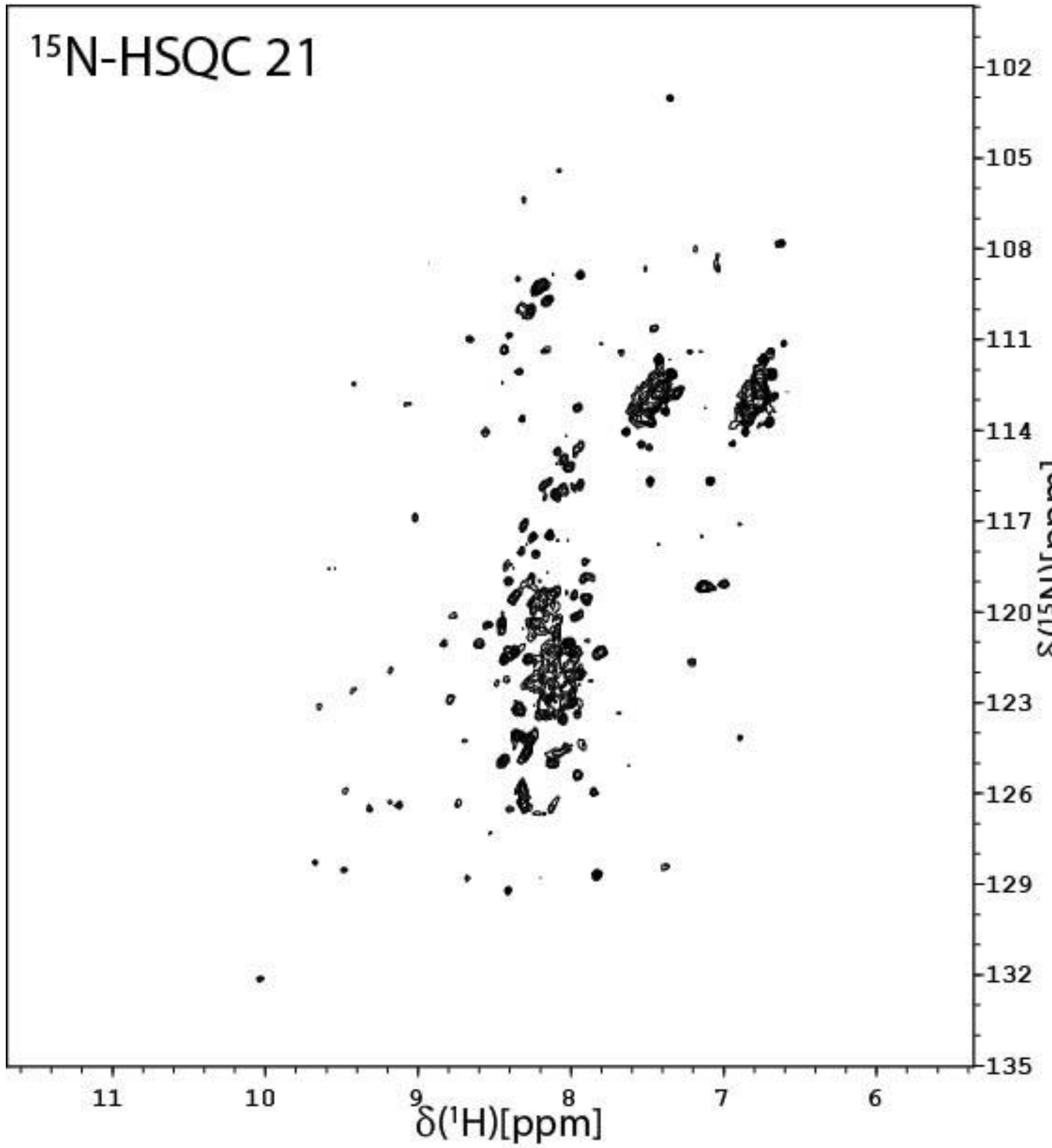


**Figure S32:** The $^{15}$N-HSQC of *holo*-*Sm*LPMO10A incubated with 10 mM ascorbic acid for 78h and then 5 mM ascorbic acid and 2 mM $H_2O_2$ for t'= 48h in acetate buffer (25 mM sodium-acetate, 10 mM NaCl, pH 5.0) in the presence of 10 mg β-chitin particles with a size of ~ 0.5 mm (24h preincubation) at 25 °C (treatment series 3, $^{15}$N-HSQC 21).

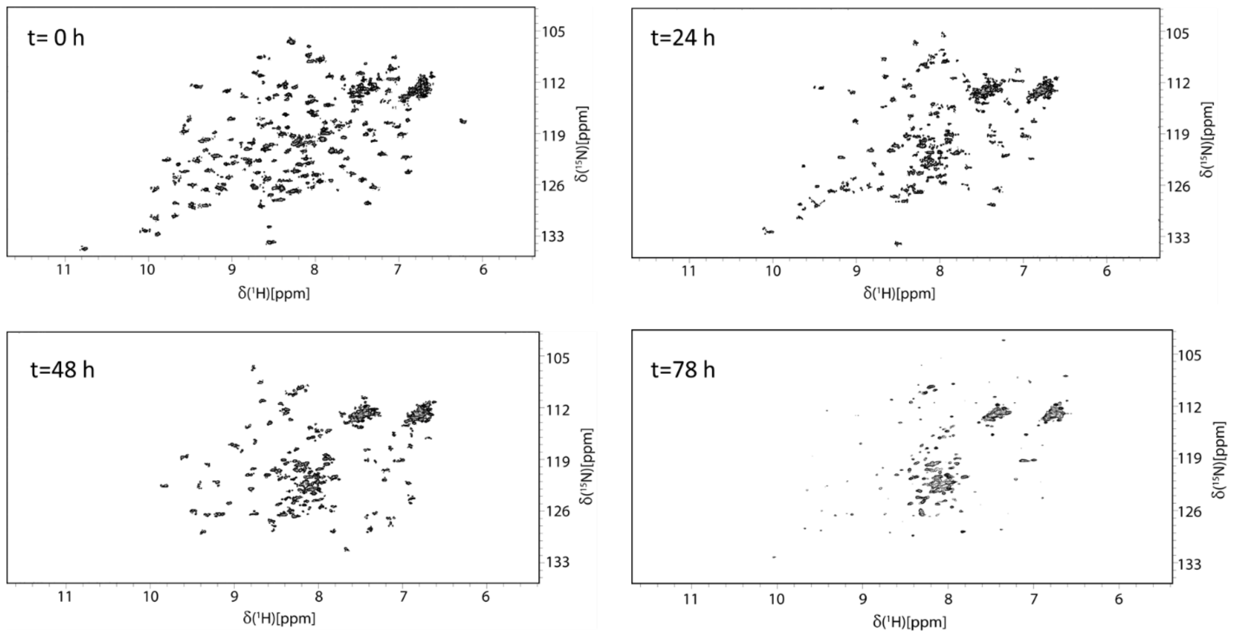


**Figure S33**: $^{15}$N-HSQC spectra of *holo-Sm*LPMO10A (treatment series 3, $^{15}$N-HSQCs 15-18) incubated with 10 mM ascorbic acid and 10 mg milled β-chitin for t= 0h, t= 24h, t= 48h, and t= 78h in acetate buffer (25 mM sodium-acetate, 10 mM NaCl, pH 5.0). Both reduced signal dispersion and signal line broadening, increased over time, indicating protein degradation. The enzyme was pre-incubated with the β-chitin particles for 24h before the addition of ascorbic acid.

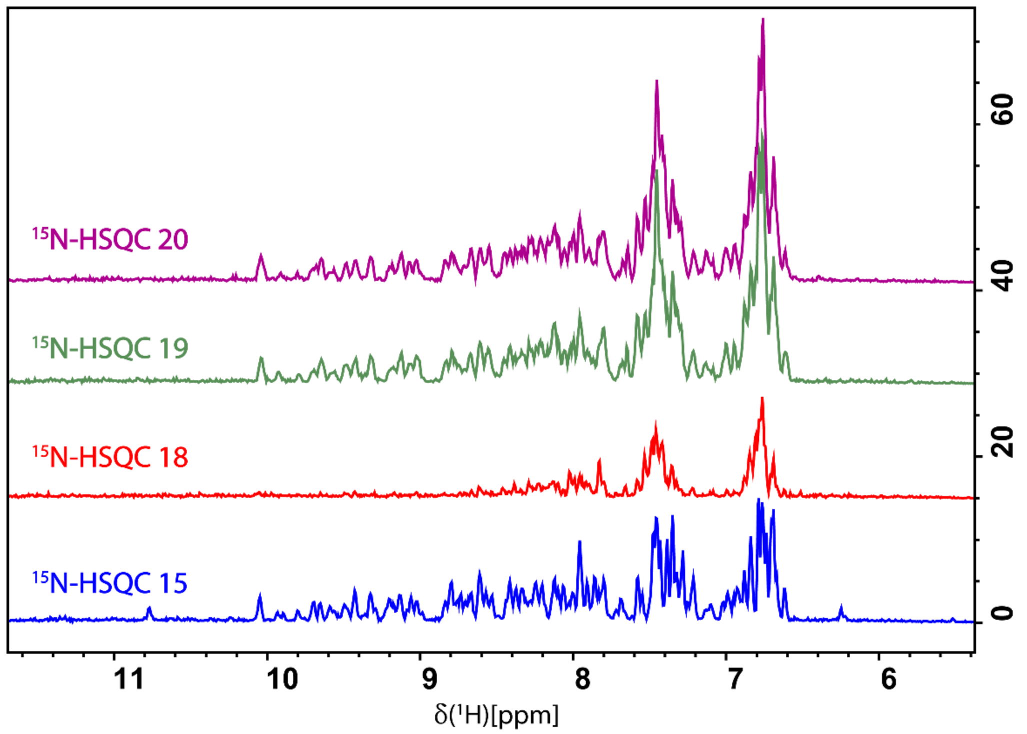


**Figure S34:** $^{1}H$ projections from the 2D $^{15}N$-HSQC spectra of *holo*-*Sm*LPMO10A (pre-incubated with chitin for 24h in treatment series 3) incubated with 10 mM ascorbic acid for t= 0h ($^{15}N$-HSQC 15), t= 78h ($^{15}N$-HSQC 18), and, subsequently, 5 mM ascorbic acid + 2 mM $H_2O_2$ for t'= 0h ($^{15}N$-HSQC 19) and t'= 24h ($^{15}N$-HSQC 20).

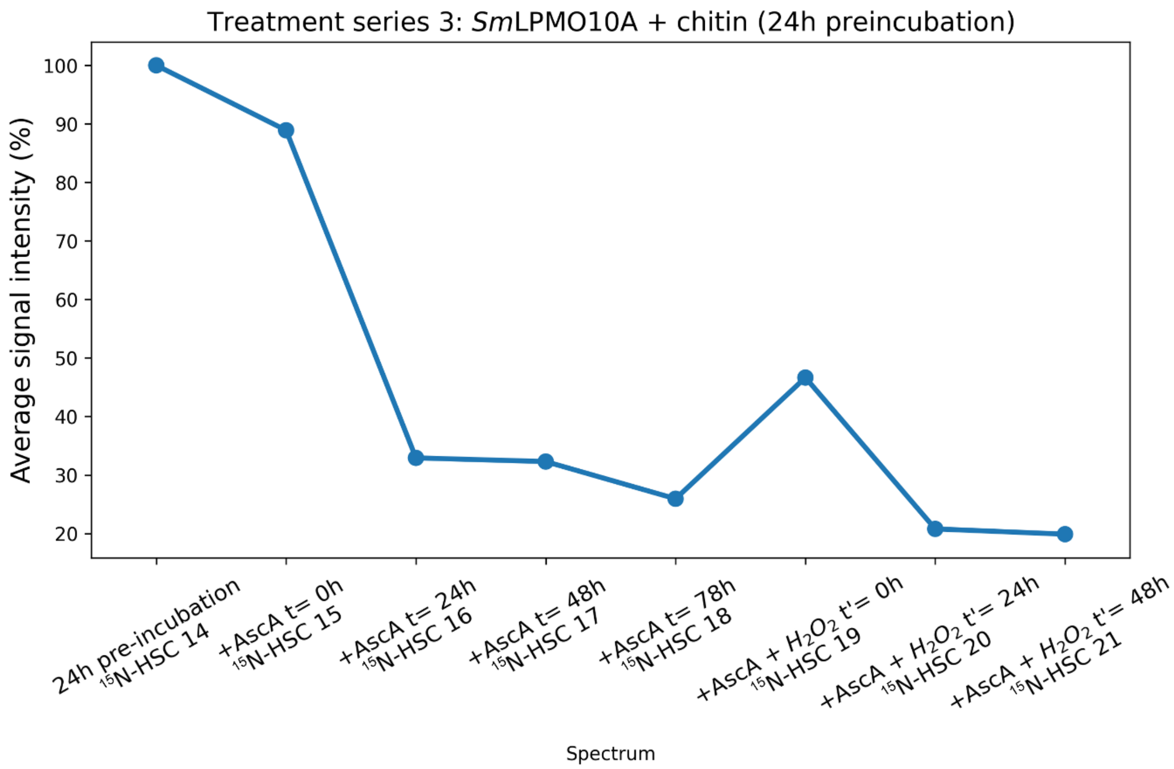


**Figure S35**: Average $^{1}$H-signal intensities in $^{15}$N-HSQC spectra 14-21 recorded for treatment series 3. *holo*-*Sm*LPMO10A was pre-incubated for 24 h with chitin, after which it was first exposed to 10 mM ascorbic acid alone for 78 h, and then to 5 mM ascorbic acid and 2 mM $H_2O_2$ for 48 h. The $^{1}$H intensities of all the $^{1}$H-$^{15}$N signals in $^{15}$N-HSQC spectra 15-19 were integrated using CARA [46].
